# Profiling a single-stranded DNA region within an rDNA segment that is a loading site for bacterial condensin

**DOI:** 10.1101/2021.06.17.448897

**Authors:** Koichi Yano, Hideki Noguchi, Hironori Niki

## Abstract

Bacterial condensin preferentially loads to single-stranded DNA (ssDNA) in vitro and loads onto rDNA in vivo to support proper chromosome compaction. Thus, the actively transcribing rDNA would provide the ssDNA region for the topological loading of bacterial condensin. We attempted to detect the ssDNA region in the *rrnI* gene in situ. Non-denaturing sodium bisulfite treatment catalyzed the conversion of cytosines to thymines via uracils (CT-conversion) at locally melted DNA of a bacterial genome. Using next-generation sequencing, we generated an average of 11,000 reads covering each cytosine on the PCR-amplified rDNA segment to obtain the actual CT-conversion rate. In principle, the CT-conversion rate is an accurate guide to detect the formation of the ssDNA segment. We expected that an increment of the CT-conversion rate would reflect a trend toward ssDNA accumulation at a given site within the rDNA. We detected multiple ssDNA segments throughout the rDNA. The deletion mutations of the rDNA that affect the bacterial-condensin loading hindered the ssDNA formation only at the 100–500 bp segment downstream of the promoter. These data support the idea that the ssDNA segment plays a crucial role as the bacterial condensin-loading site and suggest the mechanism of condensin loading onto rDNA.

## Introduction

Proper organization of chromosomal DNA is crucial for the segregation of replicated chromosomes into daughter cells. The structural maintenance of chromosomes (SMC) proteins are major players in chromosomal DNA organization in prokaryotes and eukaryotes (Dame et al., 2019; Hirano, 2016; Nasmyth and Haering, 2005; Uhlmann, 2016; Yatskevich et al., 2019). The SMC proteins are a core of the two distinct complexes, cohesin and condensin, that manage chromosome cohesion and condensation. A two-armed structure of the SMC dimer forms a ring that holds DNA topologically. The topological DNA-binding activity is regulated by non-SMC subunits that interact with an ATP-binding cassette (ABC)-like domain at the distal end of each arm. In addition, the SMC proteins bind to not only double-stranded DNA (dsDNA), but also single-stranded DNA (ssDNA). The physiological significance of the ssDNA-binding activity is currently not clear. However, several reports indicate that the ssDNA binding of the SMC proteins contributes to topological DNA binding on genomic DNA (Hirano and Hirano, 1998, 2006; Keyamura and Hishida, 2019; Murayama et al., 2018; Niki and Yano, 2016; Sutani and Yanagida, 1997; Sutani et al., 2015).

The *B. subtilis* Smc-ScpAB complex consists of the core of the SMC protein Smc and the non-SMC subunits ScpA/B and functions as a bacterial condensin (Moriya et al., 1998; Soppa et al., 2002). The SMC protein specifically binds to a highly transcribed region such as rDNA and also binds to the *parS-*binding sites (Gruber and Errington, 2009). Moreover, the Smc-ScpAB complex topologically binds to rDNA (Yano and Niki, 2017). The topological binding of the Smc-ScpAB complex requires a full-length rDNA that is actively transcribed (**Figure 1**). Although transcription is still highly activated, only the 50 bp deletion in the 5S rRNA gene hinders the topological loading activity. The full-length rDNA encodes 16S, 23S, 5S, and tRNA. These transcribed rRNAs form a higher-order structure. In particular, the ssDNA formation coupled with transcription might be a leading cause of the topological loading of bacterial condensin, because *E. coli* condensin MukB topologically binds ssDNA preferentially rather than double-stranded DNA (dsDNA) (Niki and Yano, 2016). Thus, it is conceivable that a higher-order structure of the transcripts and template DNA strand or R-loop is involved in the topological loading of bacterial condensin.

**Figure 1.**
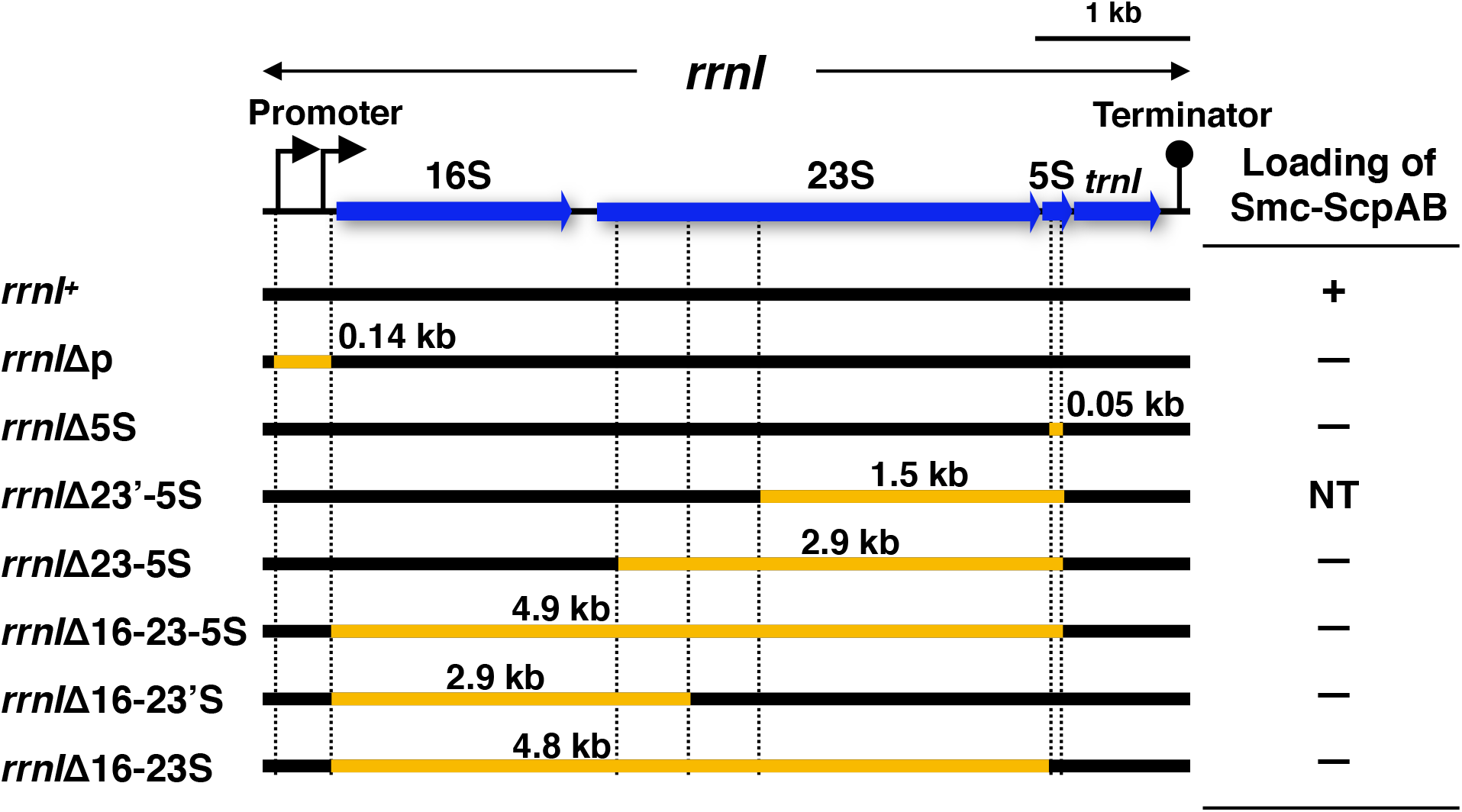
The structure of rDNA segments defective in Smc-ScpAB complex loading. The rDNA segment contains two promoters, the 16S-23S-5S rRNA, tRNA (*trnI*), and transcription terminator. The rDNA segment of the wild-type is shown at the top (*rrnI^+^*). The deleted DNA segments of the rDNA mutants are indicated in orange. The loading efficiencies of the Smc-ScpAB complex against the rDNA mutants are reported in Yano and Niki (2017).

Chromosomal DNA is ordinarily maintained as dsDNA and is partially melted as ssDNA during DNA replication and active transcription. It is conventionally considered that the ssDNA formation is detrimental to the maintenance of chromosome integrity (Zhao et al., 2010). However, recent studies indicate that non-B DNA structures play physiological roles, including roles in transcription regulation, absorption of torsional stress, and DNA replication inhibition (Edwards et al., 2009; Ginno et al., 2012; Zhao et al., 2010). In addition, dsDNA locally “breathes” to melt one to several base pairs and is used for the homology probing during homologous recombination (Renkawitz et al., 2014). Several methods have been used to detect ssDNA, such as probing with an anti-DNA-RNA hybrid antibody against the R-loop and nuclease mapping for the ssDNA segments. In addition, bisulfite genomic sequencing, a gold-standard method to detect DNA methylation, also helps determine the ssDNA segments in genomic DNA (Frommer et al., 1992; Gentry and Hennig, 2016; Hayatsu, 2008; Shah et al., 2020; Yu et al., 2003). Bisulfite selectively deaminates cytosine residues of denatured DNA and catalyzes unpaired cytosines to uracils (Hayatsu et al., 1970; Shapiro et al., 1970). PCR amplification of the chromosomal segment of interest converts the uracils into thymines. Non-denaturing sodium bisulfite sequencing is suitable for detecting the ssDNA segments in bacterial genomes because of the low methylation and poor chromatin structure of the chromosomal DNA (Leela et al., 2013). Thus, it is possible in principle to detect the ssDNA segment in situ based on the CT conversion rate. In this study, we quantified the levels of ssDNA formation in the bacterial rDNA at single-cytosine resolution by using non-denaturing sodium bisulfite treatment combined with deep sequencing analysis.

## Results

Based on the fact that bacterial condensin preferentially loads to single-stranded DNA (ssDNA) in vitro and loads onto rDNA in vivo to support proper chromosome compaction, we hypothesized that the rDNA provides the ssDNA region for the topological binding of the Smc-ScpAB complex. Therefore, we attempted to detect the ssDNA region on the rDNA segment in a living bacterial cell. Using next-generation sequencing (NGS), we generated 11,000 reads, on average, that cover each cytosine on the PCR-amplified rDNA segment to obtain the precise CT conversion rate. The results showed that the CT conversion rates were higher than the frequency of error generation by NGS. Therefore, we expected that an increment of the CT conversion rate would reflect a trend toward ssDNA accumulation at a given site within the rDNA segment.

### In situ detection of the ssDNA region in rDNA on the chromosome

The wild-type cell of *B. subtilis* has ten copies of rDNAs. By contrast, the “2 rrn strain” has only two copies of the rRNA gene due to deletions in the other eight rDNAs. The specific rDNA segment of the 2 rrn strain was easily amplified by PCR. The 2 rrn strain exhibited slight defects in cell growth (Yano et al., 2013) and had nucleoids that were separated and constricted in a dumbbell shape, as seen in the wild-type cell. Moreover, we analyzed the topological binding of the Smc-ScpAB complex to the deletion mutation rDNA segment in the 2 rrn strain, which has only two copies of the rRNA gene, *rrnA* and *rrnI* (Yano and Niki, 2017). Then we used a series of the 2 rrn strains harboring the deletion mutation rDNA segments to detect the ssDNA region on the rDNA segment (**Figure 1**).

First, we analyzed the CT conversion rates of the wild-type *rrnI* gene of the 2 rrn mutant without the sodium bisulfite treatment. We determined the CT conversion rates of each cytosine at the top and the bottom strand in the 5’ half of the *rrnI* gene. The CT conversion rates of the top strand were similar to those of the bottom strand (**Figure 2A**). We performed a sliding window analysis with the window size fixed at 64 bp and moving away from the promotor region of *rrnI* in one base step. The average CT conversion rate of each window was mapped on the *rrnI* gene to compensate for the irregular cytosine distributions in the gene. There were no characteristic profiles of the CT conversion rates at either strand. The average CT conversion rate was 0.16% (**Table S1**). When cells were not exposed to sodium bisulfite, the low CT conversion rates were well consistent with the error rate caused by the misreading of NGS (0.1~0.3%).

**Figure 2.**
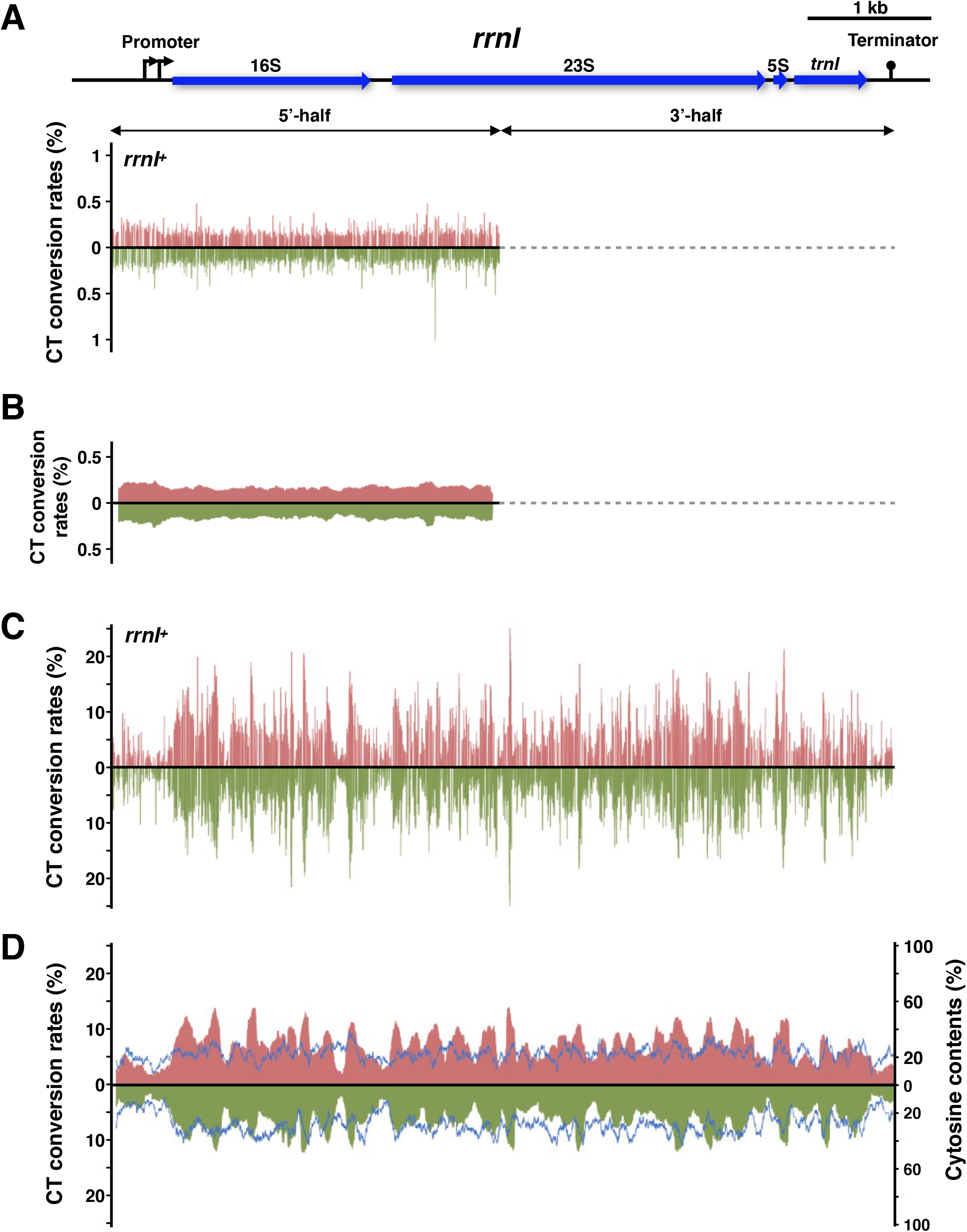
The CT conversion rates of the wild-type rDNA segment (*rrnI*) A. The CT conversion rates at each cytosine of the *rrnI* segment without sodium bisulfite treatment. The CT conversion rates of the top (red) and bottom (green) strands are represented. The lines with arrowheads indicate the DNA segments amplified by PCR, i.e., the 5’ half and the 3’ half of the *rrnI* gene.
B. The CT conversion rates of the *rrnI* segment without sodium bisulfite treatment are represented as the results of the sliding window analysis (window size of 64 bp).
C. The CT conversion rates at each cytosine of the *rrnI* segment with sodium bisulfite treatment.
D. The CT conversion rates of the *rrnI* segment with sodium bisulfite treatment are represented as the results of the sliding window analysis (window size of 64 bp). Cytosine content at each window is shown in blue.

See also Figures S1 and S2 and Tables S1-S7.

By contrast, the top and bottom strand CT conversion rates in the 5’ half of the *rrnI* gene were about 40-fold increased when cells were exposed to sodium bisulfite (**Table S1**). The average CT conversion rate was 6.89%. The CT conversion rates fluctuated dramatically at a single-nucleotide resolution throughout the *rrnI* gene; the lowest rate was 0.81%, and the highest rate was 21.5% (**Figure 2C**). We further analyzed the CT conversion rates of the 3’ half of the *rrnI* gene and then characterized the CT conversion rates throughout the *rrnI* gene. Similar results were obtained (**Table S1** and **Figure 2C**). The CT conversion rates fluctuated dramatically throughout the *rrnI* gene. The average CT conversion rate of the *rrnI* gene was 6.77%, and the highest rate was 25.6%. The nucleotide positions corresponding to the higher CT conversion rates (>15%) on the top and the bottom strand are listed with the neighbor sequences in **Tables S2 and S3**. In addition, nucleotides with higher CT conversion rates were simultaneously found on both the top and the bottom strand of the DNA segment (**Figures S1 and S2**). The nucleotides with higher CT conversion rates were positioned within relatively higher GC content, except for bp position 669 at the top strand, which was located at a DNA segment with relatively high AT content (**Table S2**). Thus, the higher CT conversion frequencies in the *rrnI* gene might not be caused by high AT content at the DNA segment.

### Relationship between the CT conversion rates and the C content

The sliding window analysis showed that the CT conversion rates fluctuated dramatically throughout the whole *rrnI* gene. We conjectured that the cytosine content within the sliding window affected the fluctuation of the CT conversion rates because the cytosine content within the window size used (64 bp) is not constant in the *rrnI* gene (**Figure 2D**). We analyzed the correlation coefficient between the CT conversion rates and the cytosine contents (**Figure 3A**). Weak positive correlations were observed in the top and the bottom strand. The correlation coefficients were 0.41 and 0.53, respectively.

**Figure 3.**
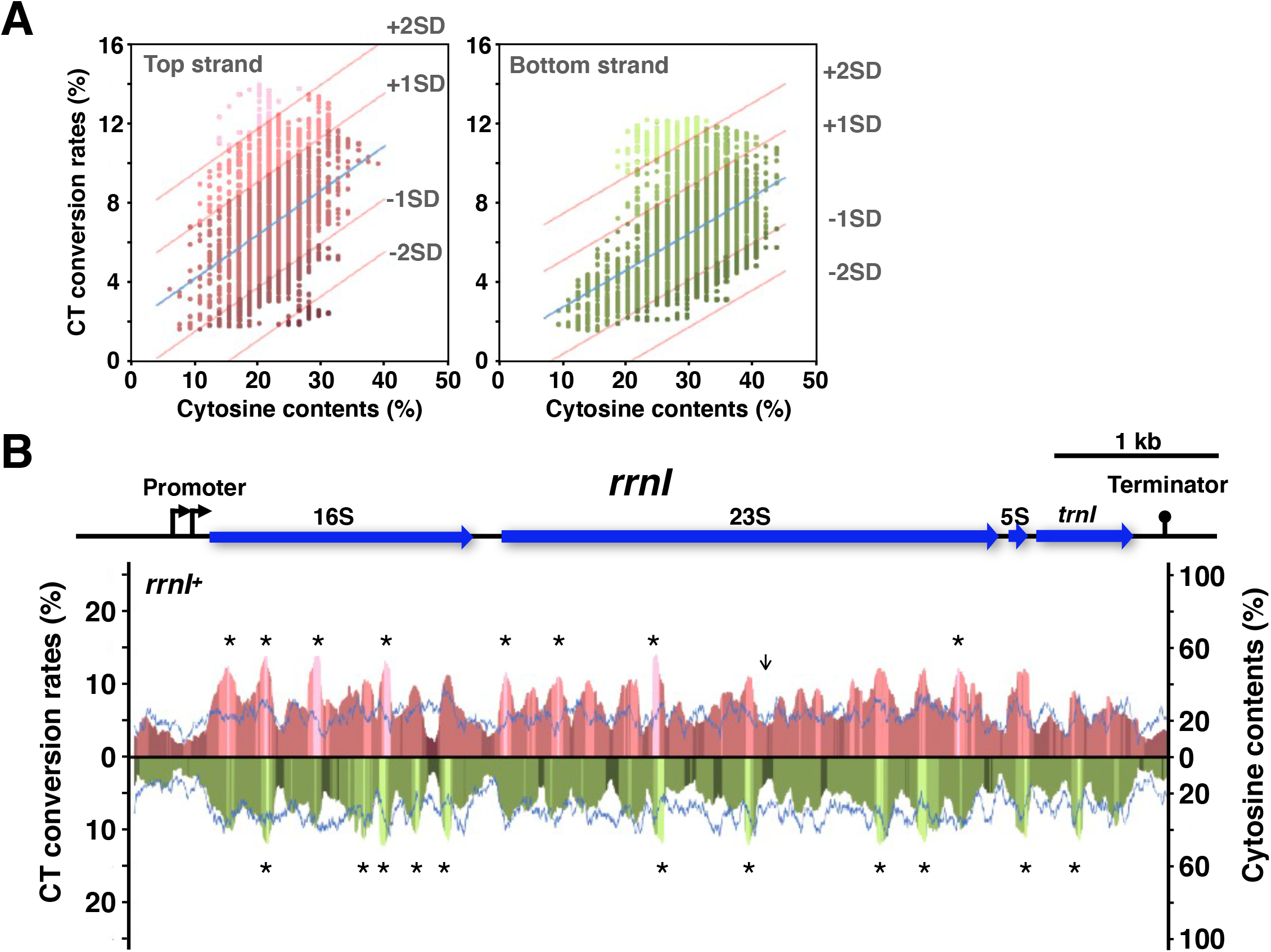
Relationship between the CT conversion rates and the cytosine content. A. Scatter diagrams of the correlation between the CT conversion rates and the cytosine content, the top strand of the *rrnI* gene (left), and the bottom strand of the *rrnI* gene (right). The CT conversion rates and the cytosine content were calculated by sliding window analysis (window size of 64 bp). Regression lines are shown in blue. The higher and lower CT conversion rates are categorized using standard deviations (±1SD and ±2SD) of the CT conversion rates. The graph points of the CT conversion rates are categorized as follows: >+2SD, +2SD< & >+1SD, +1SD< & > −1SD, −1SD < & > −2SD, < −2SD. The graph points within each category are shown in red (top strand) or green (bottom strand) hues of different brightness.
B. The graph points of the higher and the lower CT conversion rates are mapped on the *rrnI* segment, as shown in Figure 2D. The CT conversion rates on the top strand (five red hues) and bottom strand (four green hues), and the cytosine content (blue) are shown. The red and green hues correspond with those in Figure 3A. The *rrnI* map, including transcription start sites at the promoter region, 16S, 23S, 5S rRNA genes, tRNA gene (*trnI*), and transcription terminator, are indicated above the graph. See also Figures S3 and S4.

However, several segments were plotted apart from the main correlation lines. Indeed, some windows showed that the CT conversion rates were relatively high even though the cytosine content (C content) was low. Conversely, other windows showed that the CT conversion rates were relatively low even though the cytosine content was high. We selected the sliding windows with the higher and lower CT conversion rates (more and less than 1SD, respectively) from **Figure 3A** and plotted them on the *rrnI* gene map along with the CT conversion rates (**Figures 3B, S3, and S4, Tables S4, S5, and S6**). We confirmed that the CT conversion rates at the specific segments of the *rrnI* gene increased independently of the cytosine content. These results suggest that the *rrnI* gene contained the DNA segments that tend to form ssDNA.

### Relation between CT conversion and transcription at the rDNA

Next, we examined the effect of active transcription on the CT conversion rates. We analyzed the CT conversion rates of the *rrnI*Δp gene in which the promoter region was deleted (**YAN12675; Figure 4A**). The transcriptional activity of the *rrnI*Δp gene was reduced by 99% compared with that of the wild-type *rrnI* gene (Yano and Niki, 2017). However, a remarkable CT conversion of *rrnI*Δp was detected in the *rrnI*Δp gene. Figure 4 shows the profiles of the CT conversion rates of the single nucleotides and the sliding window analysis. The whole profiles were strikingly similar in appearance to the profiles of the wild-type *rrnI* gene. However, we considered that the marked decline in transcription activity caused some differences in the CT conversion rates between the wild-type and the *rrnI*Δp. We then subtracted the *rrnI*Δp CT conversion rates from the wild-type CT conversion rates for further analysis (**Figure 4C**). We found that, on the whole, the wild-type CT conversion rates were higher than the *rrnI*Δp conversion rates. The transcription activity from the *rrnI* promoter does not directly affect DNA melting in the segment upstream of the *rrnI* promoter. Certainly, the CT conversion rate differences between the wild-type and the *rrnI*Δp were slight at the segment upstream of the *rrnI* promoter. The average difference in the CT conversion rate at the promoter’s upstream segment was 0.52% (**Table S7**). On the other hand, the average difference in the CT conversion rate was a 2.06 percentage poin in the promoter’s downstream segment. Moreover, several segments showed differences of more than a 4 percentage poin in the CT conversion rate.

**Figure 4.**
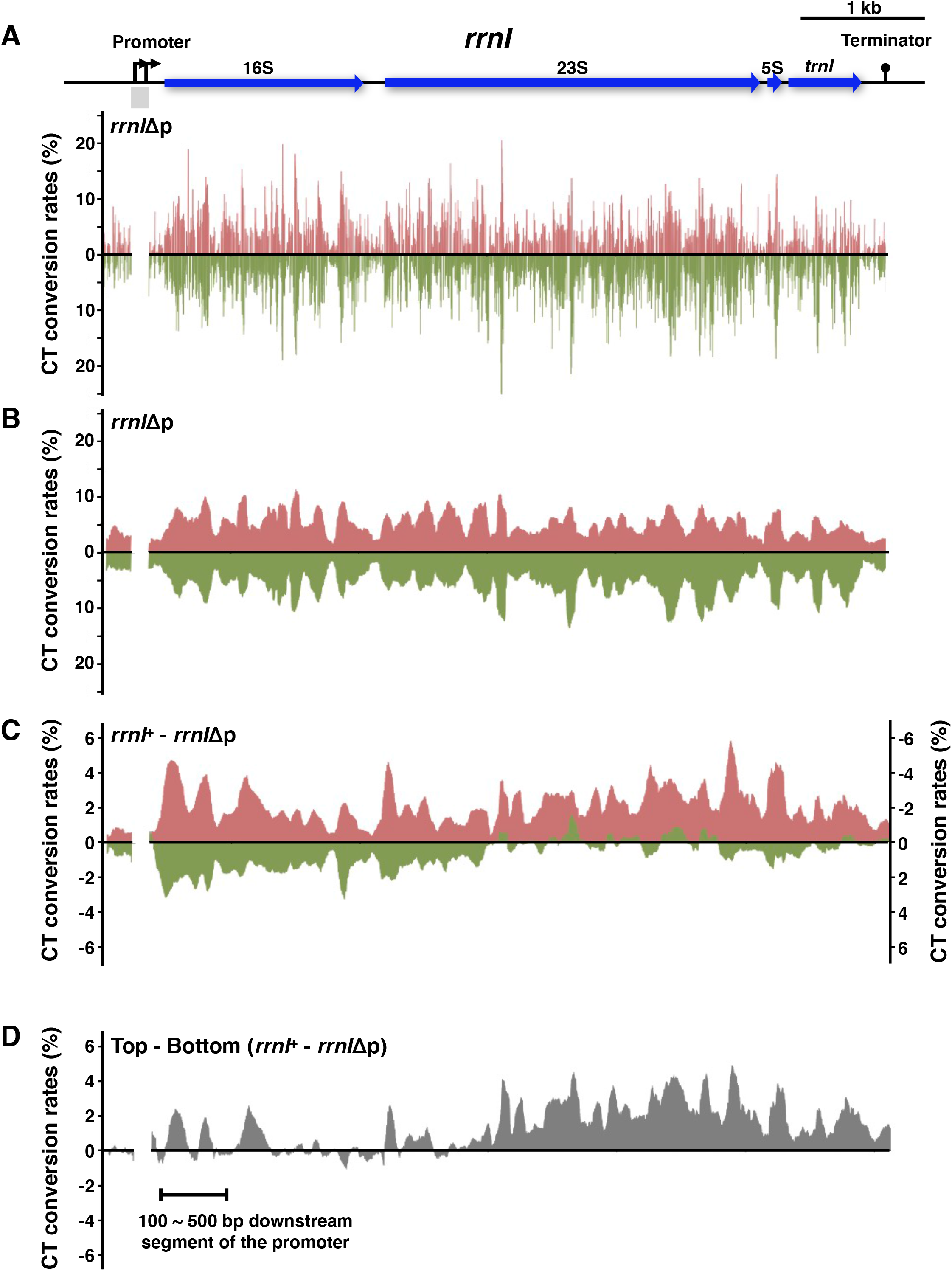
The CT conversion rates of the *rrnI*Δp segment. A. The CT conversion rates at each cytosine of the *rrnI*Δp segment with sodium bisulfite treatment. The CT conversion rates of the top (red) and the bottom (green) strand of the *rrnI*Δp segment are represented. The deleted region is indicated with a gray box in the *rrnI* gene map.
B. The CT conversion rates of the *rrnI*Δp segment with sodium bisulfite treatment are represented as the results of sliding window analysis (window size of 64 bp).
C. The differences of the CT conversion rates between the *rrnI*Δp segment and the wild-type *rrnI* segment. The left vertical line shows the differences in the CT conversion rates for the top strand, and the right vertical axis shows those for the bottom strand.
D. The differences of the CT conversion rates between the top and the bottom strand as shown in Figure 4C (*rrnI* - *rrnI*Δp). The solid line shows the segment 100 to 500 bp downstream of the promoter.

Notably, the differences in the CT conversion rates were particularly large in the top strand, i.e., the coding DNA strand. On the other hand, the differences in the CT conversion rates in the bottom strand, i.e., the template DNA strand, were mainly found at the former part of the *rrnI* gene but not at the latter part.

Next, we subtracted the CT conversion rates of the top strand from those of the bottom strand (**Figure 4D**). The degree of influence caused by reducing the transcription activity was substantial in the top strand of the *rrnI* gene, and the influence was especially remarkable at the latter part of the top strand. The two peaks of the biased CT conversion rates at the segment 100 to 500 bp downstream of the promoter indicate that the R-loop formation was affected by the reduction in transcription activity.

### The CT conversion of the *rrnI*Δ5S mutant

We carried out a further analysis to specify the portion of the ssDNA region within the *rrnI* gene that affects the topological loading of Smc-ScpAB bacterial condensin. We analyzed the CT conversion rate of the *rrnI*Δ5S gene (**YAN12698**), in which the 51 bp segment of the 5S rRNA coding region (116 bp) was deleted (**Figure 5A**). Although the mutated *rrnI*Δ5S gene results in a defect in the topological loading of the Smc-ScpAB, it is actively transcribed (Yano and Niki, 2017). The whole profiles of the CT conversion of *rrnI*Δ5S shown by both the single nucleotides and the sliding window analysis were strikingly similar in appearance to those of the wild-type *rrnI* gene (**Figure 5A and B**). As in the analysis of *rrnI*Δp, the CT conversion rate of the *rrnI*Δ5S gene was subtracted from that of the wild-type *rrnI*. As a result, we found marked differences in the CT conversion rates in the segment 100 to 500 bp downstream of the promoter but not in the other segments (**Figure 5C**). The CT conversion rates within the segment 100 to 500 bp downstream of the promoter were higher than the average CT conversion rate of the *rrnI* segment. Similar results were obtained from the bottom strand.

**Figure 5.**
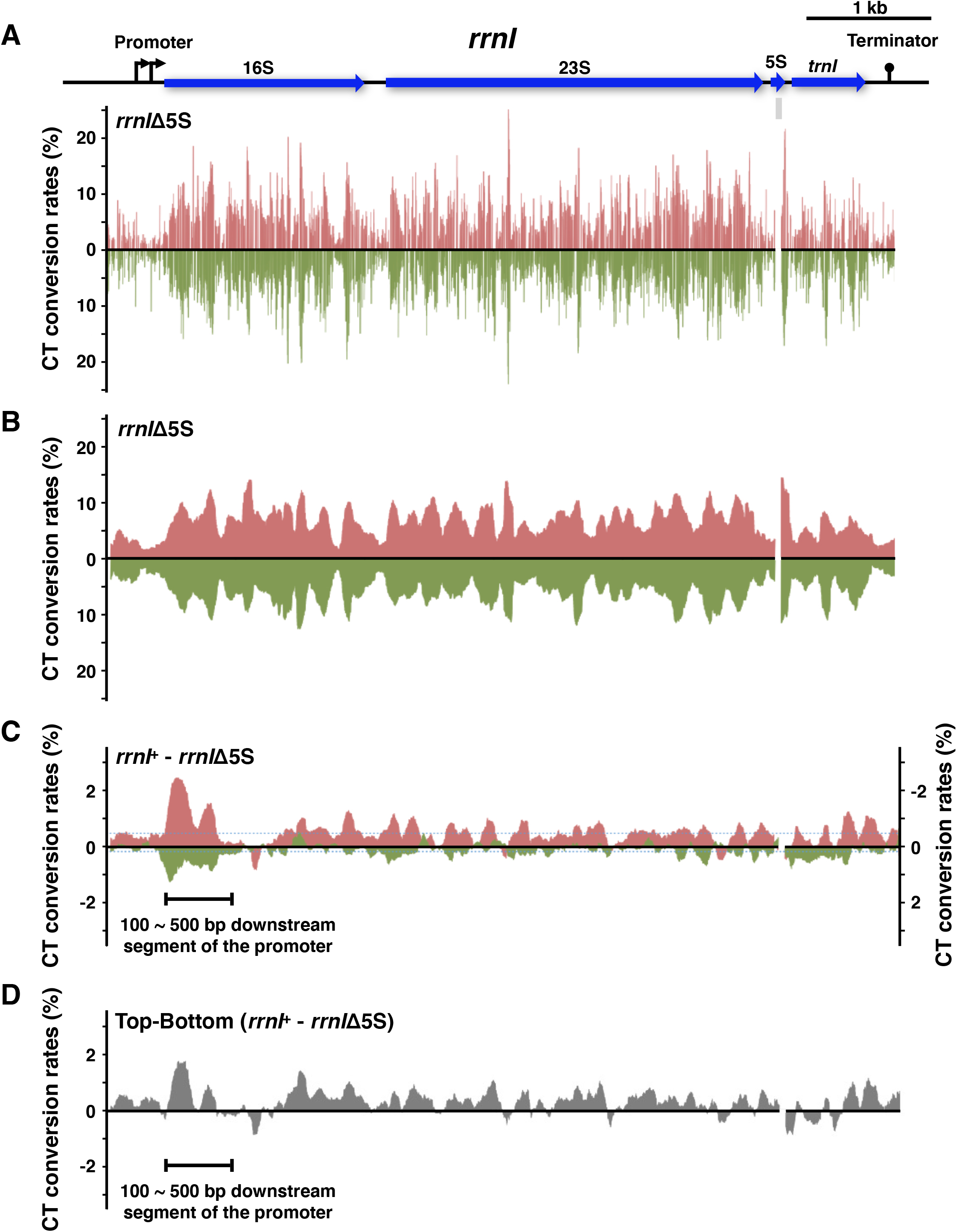
The CT conversion rates of the *rrnI*Δ5S segment. A. The CT conversion rates at each cytosine of the *rrnI*Δ5S segment with sodium bisulfite treatment. The CT conversion rates of the top (red) and the bottom (green) strand of the *rrnI*Δ5S segment are represented. The deleted region is indicated with a gray box in the *rrnI* gene map.
B. The CT conversion rates of the *rrnI*Δ5S segment with sodium bisulfite treatment are represented as the results of sliding window analysis (window size of 64 bp).
C. The differences of the CT conversion rates between the *rrnI*Δ5S segment and the wild-type *rrnI* segment. The left vertical line shows the differences in the CT conversion rates for the top strand, and the right vertical axis shows those for the bottom strand.
D. The differences of the CT conversion rates between the top and the bottom strand as shown in Figure 5C (*rrnI* - *rrnI*Δ5S). The solid line shows the segment 100 to 500 bp downstream of the promoter.

We subtracted the CT conversion rates of the bottom strand from those of the top strand (**Figure 5D**). There were two peaks in the CT conversion rates of the top strand in the segment 100 to 500 bp downstream of the promoter, as shown in Figure 4D. The deletion of the 51 bp segment at the 5S rRNA coding region specifically affected the CT conversion rates of the top strand. Thus, the efficiency of ssDNA formation within the *rrnI*Δ5S gene was reduced only at the downstream segment of the promoter.

### The CT conversion of an rDNA mutant defective in the loading of the Smc-ScpAB complex

Next, we analyzed other deletion mutants of the *rrnI* gene in the 2 rrn strain (**Figure 1**). These mutants of the *rrnI* gene maintained active transcription but lost their activity for loading the Smc-ScpAB complex. The whole profiles of the CT conversion of the mutated *rrnI* genes are shown in **Figures 6 and S5**. As shown by the sliding window analyses of *rrnI*Δ23’-5S and *rrnI*Δ23-5S, the whole profiles of the CT conversion of the remaining segments were similar in appearance to those of the wild-type *rrnI* gene (**Figure 2D**). We found that the CT conversion rates were affected in the segment 100 to 500 bp downstream of the promoter when we subtracted the CT conversions of *rrnI*^Δ23’-5S^ from the wild-type or the CT conversions of *rrnI*^Δ23-5S^ from the wild-type. Other deletion mutants of the *rrnI* gene lost the region containing the segment 100 to 500 bp downstream of the promoter. The remaining segment was similar to that of the wild-type *rrnI* gene, and the differences were very small between the wild-type and the *rrnI* mutant genes. The results for both *rrnI*Δ23’-5S and *rrnI*Δ23-5S supported the idea that the segment 100 to 500 bp downstream of the promoter had the greatest effect on the CT conversion rates throughout the *rrnI* segment.

**Figure 6.**
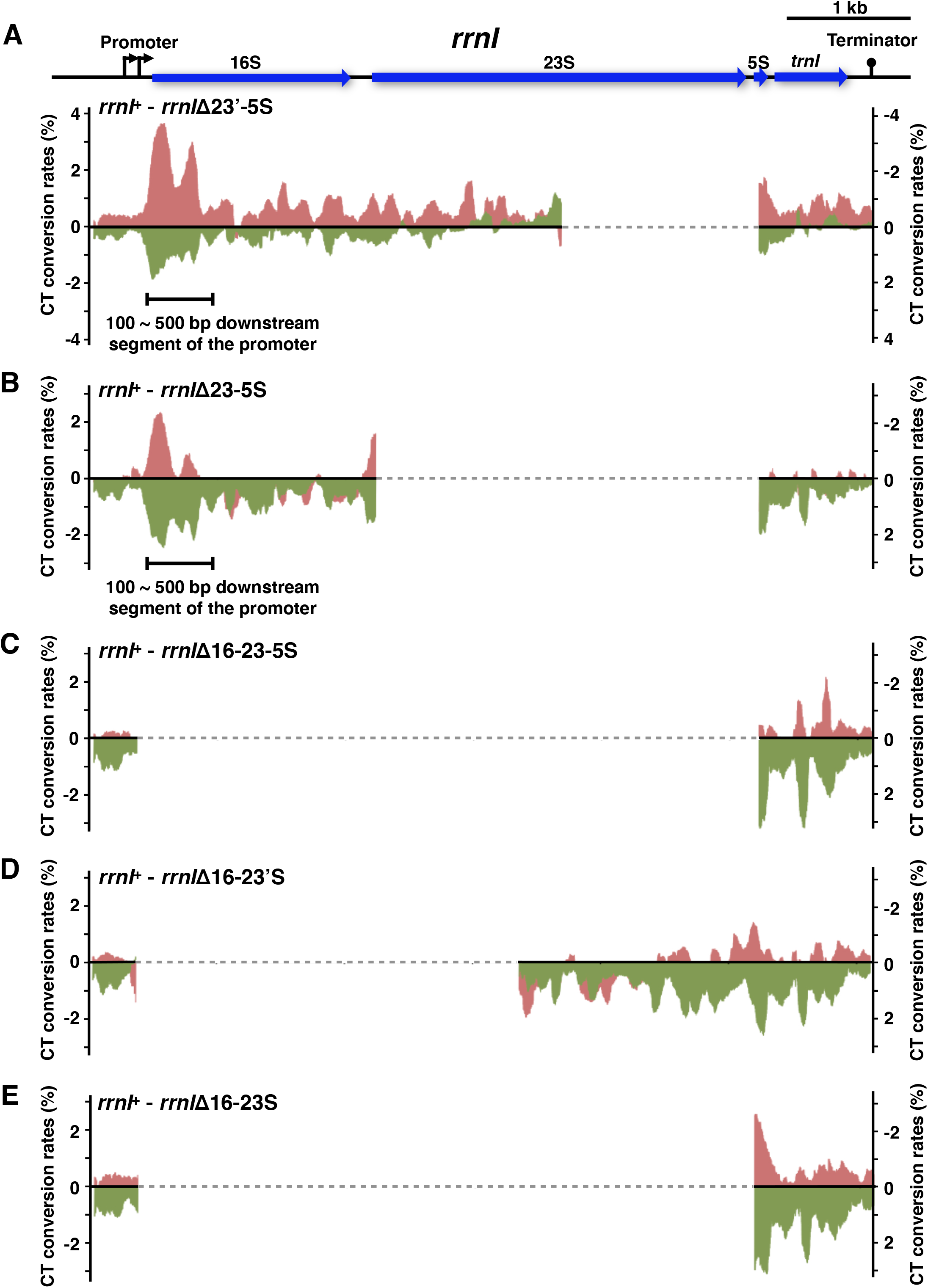
The CT conversion rates of the *rrnl* segments with various lengths of deletion mutation. The deleted regions are shown as dashed lines. The left vertical line shows the CT conversion rates for the top strand, and the right vertical axis shows the CT conversion rates for the bottom strand. (A-E) The differences in the CT conversion rates between the *rrnI*Δ23’-5S (A), *rrnI*Δ 23-5S (B), *rrnI*Δ16-23-5S (C), *rrnI*Δ16-23’S (D), and *rrnI*Δ16-23S (E) segments and the wild-type *rrnI* segment. (A and B) The solid line shows the segment 100 to 500 bp downstream of the promoter. See also Figure S5.

### The CT conversion of inactive genes during vegetative growth

We evaluated the difference in the effect of the non-denaturing sodium bisulfite treatment between the wild-type and the mutant *rrnI* genes. Because the mutant *rrnI* cells did not exhibit the same growth as the wild-type *rrnI* strains, and the mutant *rrnI* strains slowed cell growth, we analyzed the CT conversion rates of another chromosomal segment as an internal standard to consider the effect of cell growth on the CT conversion rates. The *spsA* and the *spsB* genes are inactive during vegetative growth and are activated by a sporulation-specific sigma factor, σ^K^, in response to the deterioration of nutrients in a growing environment (Eichenberger et al., 2004; Nicolas et al., 2012). Thus, the *spsAB* segment is suitable for analysis of the effect of the non-denaturing sodium bisulfite treatment on cell activities other than transcription.

We analyzed the CT conversion rates of the DNA segment, including the *spsAB* genes in the wild-type *rrnI* strain. Figures 7A and 7B show the whole profile of the CT conversion of the *spsAB* segment by both the single nucleotides and the sliding window analysis. The top and bottom strand nucleotides showed markedly higher CT conversion rates (more than 10%). In particular, the CT conversion rates were high in the region upstream of the promoter on the top strand. The average CT conversion rates of the *spsAB* segment were 4.20% on the top strand and 3.65% on the bottom strand (**Table S7**). These rates were less than those for the *rrnI* and *rrnI*Δp genes. Thus, the CT conversion of a chromosomal segment decreased due to inactivation in transcription.

**Figure 7.**
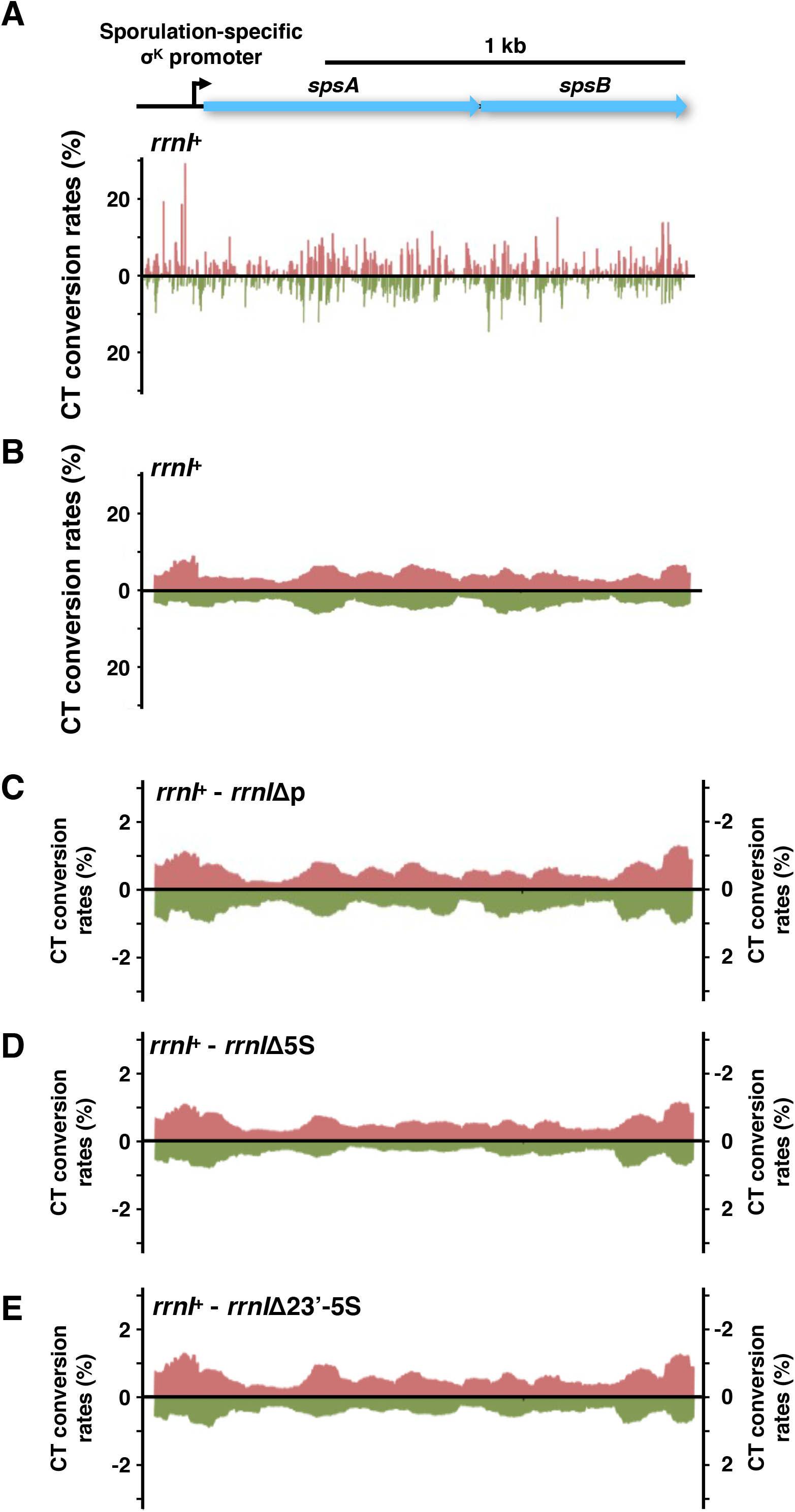
The CT conversion rates of the *spsAB* segment. (A) The CT conversion rates at each cytosine of the *spsAB* segment in cells with sodium bisulfite treatment.
(B) The CT conversion rates of the *spsAB* segment are represented using the sliding window analysis (window size of 64 bp).
(C-E) The differences of the CT conversion rates of the *spsAB* segment between the wild-type *rrnI* and the *rrnI* deletion mutants: *rrnI*Δp (C), *rrnI*Δ5S (D), and *rrnI*Δ23’-5S
(E). The map of the *spsA* and *spsB* genes, including the promoter, are indicated at the top.

See also Figures S6 and S7 and Tables S1 and S7.

Next, we analyzed the CT conversion rates of the *spsAB* gene in the 2 rrn strains harboring a series of the mutated rDNA segments. We obtained whole profiles of the CT conversion of the *spsAB* segment by sliding window analysis (**Figure S6**). The results showed that the *spsAB* segment profiles in the mutated rDNA strains were similar. Moreover, the profiles were also very similar to that in the 2 rrn strain harboring the wild-type *rrnI*. In fact, we did not find any apparent differences when the CT conversion rates in the mutated rDNA strains were subtracted from those of the 2 rrn strain harboring the wild-type *rrnI* (**Figures 7C, 7D, 7E, and S7**). The differences in the CT conversion rate between the wild-type and the mutated *rrnI* strains were less than a 0.7 percentage point. Thus, the deletion within the *rrnI* gene did not affect the CT conversion of the other genes on the genome.

## Discussion

We used a combination of biochemical and genomic approaches to detect where chromosome DNA segments melt into ssDNA in a bacterial cell. Non-denaturing sodium bisulfite treatment effectively catalyzed the conversion of unpaired cytosines to thymines via uracils without any DNA denaturation treatments. The CT conversion induced by non-denaturing sodium bisulfite treatment showed high performance for the detection of ssDNA segments in the genome by using NGS. Our analyses demonstrated that bacterial genomic DNA unpredictably becomes ssDNA in a living cell.

We applied the CT conversion rates of the *spsAB* segment to the internal standard. The differences in the CT conversion rate at the *spsAB* segment between the wild-type and the mutated *rrnI* (0.67%) strains were less than those at the *rrnI* segment (1.26%) (**Table S7**). Judging from the above, the reduction of the cell growth rate did not reflect the CT conversion rates. We carried out the non-denaturing sodium bisulfite treatment until the progression of the cytosine deamination was nearly saturated and then performed next-generation sequencing, with an average of 11,000 reads, covering each cytosine on the PCR-amplified rDNA segment. As a result, we were able to obtain the actual CT conversion rate to overcome perturbation by the cell growth rate. Therefore, we conclude that the CT conversion rate differences at the *rrnI* segment between the wild-type and the mutated *rrnI* were directly affected by the efficiency of the ssDNA formation, which deteriorates due to the deletion mutations.

When we compared the CT conversion rates between the wild-type and the mutated *rrnI* gene, the difference in the rates led us to discover ssDNA segments in the bacterial rDNA under active transcription. Multiple ssDNA segments were formed throughout the rDNA segment under active transcription. Temporary hybridization between a new RNA transcript and the template DNA strand reduced the CT conversion rate on the bottom strand. The biased CT conversion rates of the ssDNA segments indicate the RNA loop formation.

The ssDNA segment located at the segment 100 to 500 bp downstream of the rDNA promoter was essential for loading the Smc-ScpAB complex. Biases in the CT conversion rates between the top and the bottom strand were obvious in this segment. It is conceivable that the ssDNA segment was exposed by the R-loop formation in a manner dependent on the rRNA transcription. The results are consistent because the bacterial SMC proteins bind to ssDNA and preferentially load onto the actively transcribing rDNA. Therefore, we propose that the Smc-ScpAB complex is loaded to the ssDNA segment located 100 to 500 bp downstream of the *rrnI* promoter.

Malig et al. (2020) reported a sophisticated approach for identifying R-loops in the human genome by analyzing the CT conversion rates of consecutive cytosines (Malig et al., 2020). In contrast, in the present study we were able to detect R-loops of the bacterial genome straightforwardly. It is conceivable that the R-loop formation during transcription directly reflects differences in the CT conversion rates between the coding DNA strand and the template DNA strand. If so, this could be because the bacterial genome structure is simpler than the human genome structure. Modified cytosine bases in bacterial genomes are restricted to a specific motif of DNA. DNA methyltransferase of *B. subtilis* 168 (BsuM) modifies cytosine in the sequence 5’-YTCGAR-3′(Guha, 1988). Nine cytosines in the sequence were located in the *rrnI* genes. In addition, bacterial chromatin does not form a nucleosome or nucleosome-like structures. Thus, the properties of bacterial genomes make it easier to detect the R-loop formation during transcription by using the CT conversion rates (Leela et al., 2013). Our procedure used in this experiment could be applied for the analysis of the genome dynamics of prokaryotes.

The deletions in the latter half of the *rrnI* gene affected the ssDNA formation at the segment 100 to 500 bp downstream of the *rrnI* promoter. Even the 51 bp deletion in the 5S encoding segment was effective against reducing ssDNA formation at the downstream segment of the *rrnI* promoter. The transcription activity increases by 3.5-fold in the *rrnI*Δ5S gene (Yano and Niki, 2017). The ssDNA formation depended on the transcription from the *rrnI* promoter. Therefore, our results suggested that the latter half of the *rrnI* gene could somehow affect the transcription at the region upstream of the *rrnI* gene. The transcribed rRNA, including 5S rRNA, forms the secondary structure of RNA.

Moreover, the transcribed product develops a higher-order structure than the tertiary structure of transcribed products. It is a plausible explanation that the higher-order structure of the transcribed RNA, including 5S rRNA, would affect the R-loop formation at the ssDNA segment at 100 to 500 bp downstream of the *rrnI* promoter. Indeed, an R-loop was reported to be formed at the replication origin of the ColE1 plasmid and stabilized by the higher-order structure of a transcribed RNA (Masukata and Tomizawa, 1990).

High rates of CT conversion were detected in the *B. subtilis* genome, irrespective of transcription activity. The CT conversion rates by the single-nucleotide analysis fluctuated dramatically throughout the rDNA. The positions of the cytosines with the higher CT conversion rates were shared in common among the rDNA segments we analyzed. However, we could not find a prominent consensus motif in the flanking sequences of the cytosines with the higher CT conversion rates. In general, the AT-rich motifs tend to be locally melted with a complete disruption of base pairing. This was not the case in our present analysis. Instead, the flanking sequences of the cytosines with the higher CT conversion rates contained CC, CCC, CCCC, CCCCC, and CCCCCC motifs. The cytosines with the higher CT conversion rates were also detected in the *spsB* gene. Thus, melting of the rDNA was not necessarily dependent on AT content, even though the higher CT conversion was precisely reflected by the melting of dsDNA. In addition, transcription did not directly influence the higher CT conversion rates, based on the similar results for the *rrnI*Δp mutant. It may be that DNA replication was involved in the ssDNA formation of the cytosine-rich motifs. Alternatively, these ssDNA segments may have been formed due to DNA breathing by torsion of a specific conformation rather than by thermal fluctuations. Genome-wide analysis of the CT conversion rate may clarify the correlation between a DNA sequence and a tendency to form ssDNA segments.

Members of the SMC protein family possess the ssDNA-binding ability in addition to the dsDNA-binding ability. It is not yet clear how the ssDNA-binding ability of the SMC proteins contributes to the maintenance of chromosomal DNA. Bacterial SMC proteins utilize the ssDNA-binding ability for topological loading to specific sites on the genome DNA, such as the stable ssDNA segment. A thorough analysis of the ssDNA formation in the genome would help elucidate the function of the ssDNA binding of the SMC proteins for the topological loading.

## Supporting information

Yano et al_supplemental files

## Acknowledgments

We thank all members of the NIKI Laboratory for their helpful suggestions and comments. This research was supported by Grants-in-Aid for Scientific Research (18H02485 and 18K14627). This work was also supported by JSPS KAKENHI Grant Number 16H06279 (PAGS) and the Takeda Science Foundation.

## Author Contributions

KY and H. Niki conceived the study, analyzed the data, and wrote the manuscript. KY performed the experiments and computational analysis. HN assisted with the computational analysis.

## Declaration of Interests

The authors declare no competing interests.

## Supplementary Figure Legends

**Figure S1. The nucleotide sequence map of the highest CT conversion rates in the *rrnI* segment, related to Figure 2** Nucleotide sequences on the top and bottom are indicated in the 3,064 bp to 3,163 bp region with the CT conversion rates.

**Figure S2. The nucleotide sequence map of the CT conversion rates throughout the *rrnI* segment, related to Figure 2** Nucleotide sequences on the top and the bottom strands are shown every 100 bp.

**Figure S3. Enlargement of scatter diagrams of the correlation between the CT conversion rates and the cytosine content in the top strand, related to Figure 3** A scatter diagram of correlation for the top strand in Figure 3A in the left panel. The boxed areas for the CT conversion rates above the 2SD (+2SD) and below the 2SD (-2SD) are shown in the right panels, respectively. The data points are numbered, and their values are listed in Tables S4 and S5.

**Figure S4. Enlargement of scatter diagrams of correlation between the CT conversion rates and the cytosine content in the bottom strand, related to Figure 3** A scatter diagram of correlation for the bottom strand in Figure 3A is shown in the left panel. The boxed areas for the CT conversion rates below the 2SD (-2SD) are shown in the right panels. The data points are numbered, and their values are listed in Tables S6.

**Figure S5. The CT conversion rates of the *rrnI* segments with various lengths of deletion mutation, related to Figure 6** Dashed lines show the deleted regions. Transcription start sites at the promoter region, 16S, 23S, 5S rRNA genes, tRNA gene (*trnI*), and transcription terminator are indicated above the plots. The CT conversion rates of the *rrnI*Δ23’-5S (A), the *rrnI*Δ 23-5S (B), the *rrnI*Δ16-23-5S (C), the *rrnI*Δ16-23’S (D), and the *rrnI*Δ16-23S (E) segment with sodium bisulfite treatment are represented as the results of sliding window analysis (window size of 64 bp).

**Figure S6. The CT conversion rates of the *spsAB* segment, related to Figure 7** Transcription start sites at the promoter region, *spsA* and *spsB* genes are indicated in the *spsAB* gene map. The CT conversion rates of the *spsAB* segment with sodium bisulfite treatment are represented as the results of sliding window analysis (window size of 64 bp) in the 2 rrn strains harboring the *rrnI*Δ23’-5S (A), *rrnI*Δ23-5S(B), *rrnI*Δ 16-23-5S (C), *rrnI*Δ16-23’S (D), or *rrnI*Δ16-23S (E).

**Figure S7. Differences of the CT conversion rates at the *spsAB* segment between the wild-type *rrnI* and the *rrnI* mutants with various lengths of deletion, related to Figure 7** Transcription start sites at the promoter region, the *spsA* gene, and the *spsB* gene are indicated in the *spsAB* gene map. The differences of the CT conversion rates at the *spsAB* segment between the wild-type rrnI and the rrnI deletion mutant are shown: *rrnI*Δ23-5S (A), *rrnI*Δ16-23-5S (B), *rrnI*Δ16-23’S (C), and *rrnI*Δ16-23S (D). The left and right vertical axes show the CT conversion rates for the top strand and the bottom strand, respectively.

**Figure S8. The CT conversion rates at the *rrnI* segment with an increasing incubation time of bisulfite reaction, related to Methods Details** Results are shown for the top strand (red circle) and bottom strand (green circle).

**Figure S9. Schematic diagrams for PCR-amplified DNA segments from the *rrnI* genes and *spsAB* genes, related to Methods Details** The PCR-amplified DNA segments with primers are indicated in the gene map. The thick black lines indicate the region used to determine the CT conversion rates. The thick green lines indicate sequences complementary to primers. The numbers on the lines indicate base numbers.

A. *rrnI* gene. Transcription start sites at the promoter region, 16S, 23S, 5S rRNA genes, tRNA gene (*trnI*), transcription terminator, and *spc* gene are indicated.
B. *spsAB* genes. Transcription start sites, *spsA*, and *spsB* genes are indicated.

**Figure S10. Mapping efficiencies under different values in the score-min option for the Bismark mapping tool, related to Methods Details**

A. Mapping efficiencies with changing values of Y in the score-min option (L, -X, -Y).
B. Mapping efficiencies with changing values of X in the score-min option (L, -X, -Y).

## Methods

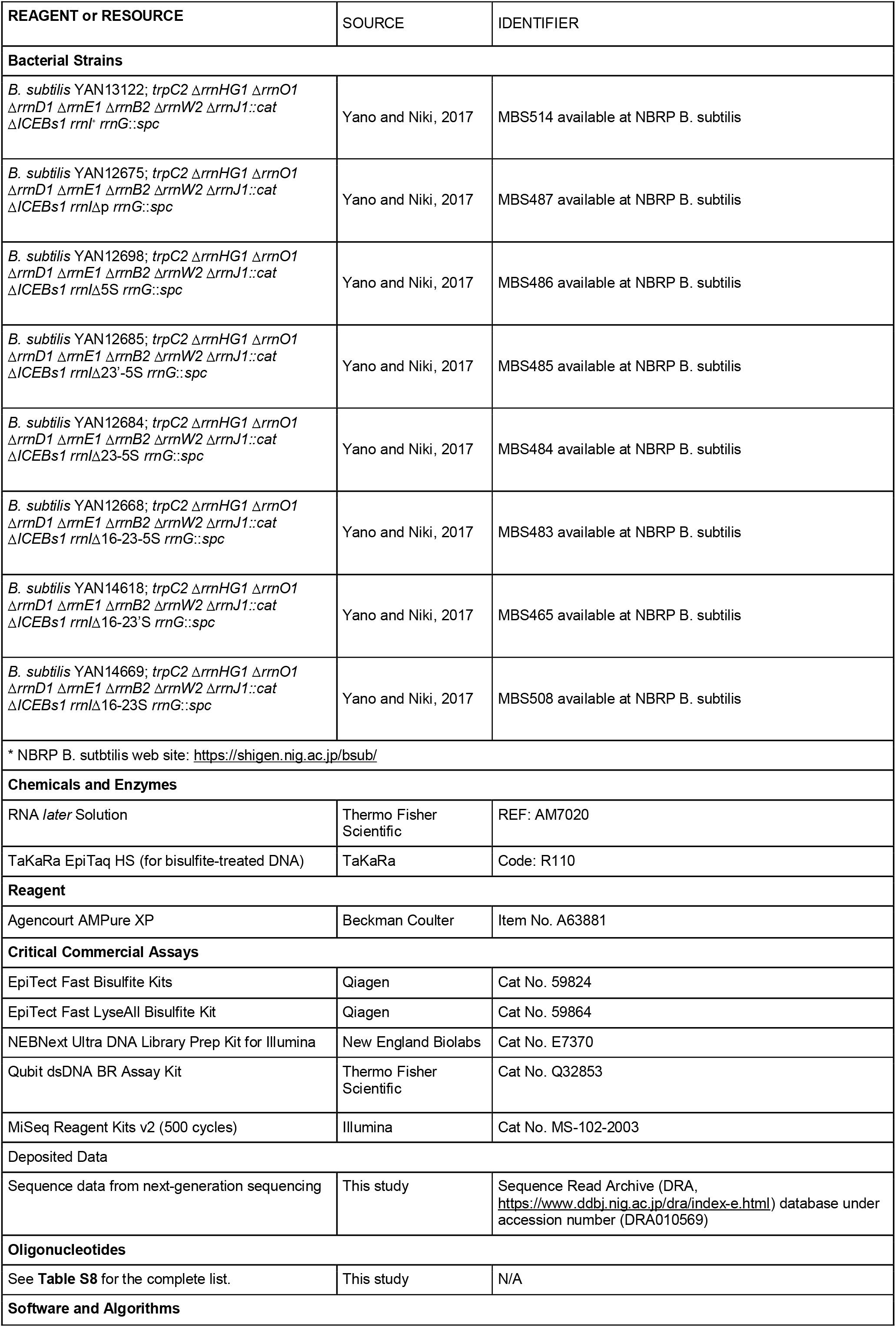
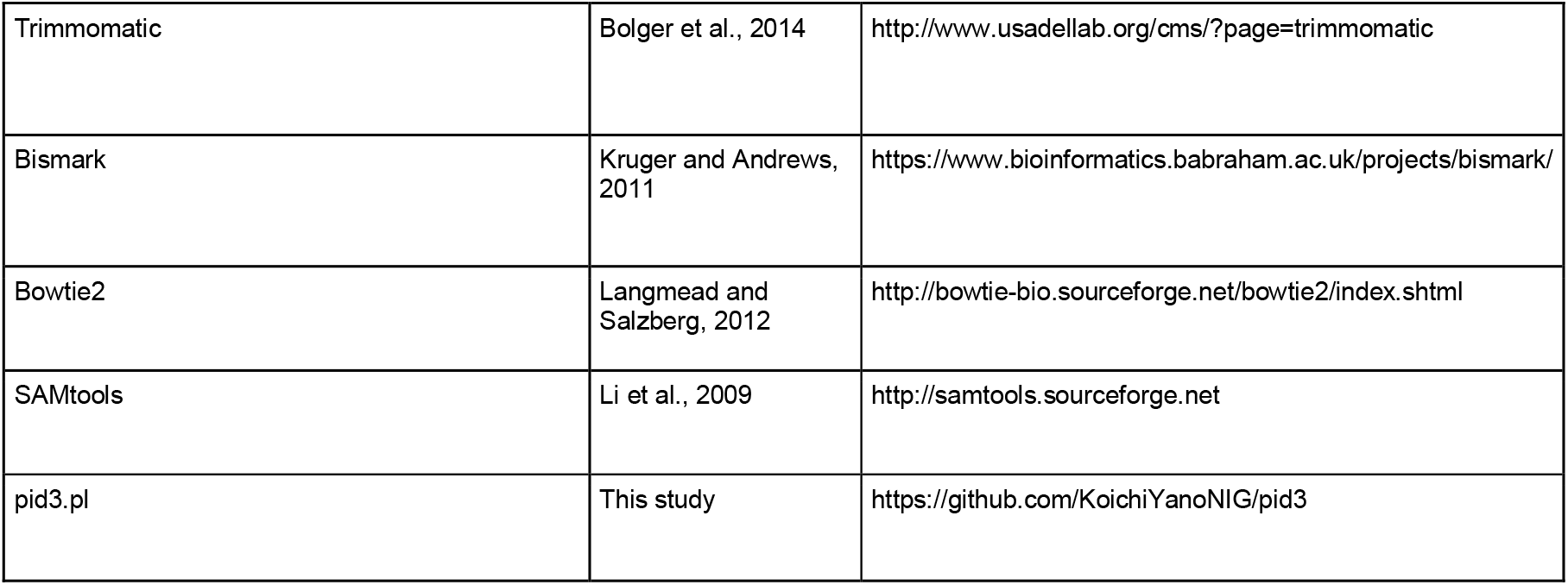
Resources Table.

### Materials availability

*Bacillus subtilis* strains used in this study are available from the National BioResource Project, *Bacillus subtilis*, JAPAN (http://www.shigen.nig.ac.jp/bsub/).

### Data and code availability

Datasets from next-generation sequencing are available at the DNA Data Bank of Japan, DDBJ (Accession No.: DRA010569). The computational script constructed in this study is available at GitHub ((https://github.com/KoichiYanoNIG/pid3).

### Experimental Model and Subject Details

#### Bacterial strains

All the strains used in this study are listed in the Key Resources Table. The 2rrn mutant, YAN13122, carries two copies of the rRNA gene (*rrnA* and *rrnI*). Various lengths of deletion in rDNA were introduced in the *rrnI* gene, as shown in **Figure 1**.

### Methods Details

#### Cultivation of cells for sodium bisulfite treatment

*B. subtilis* cells were grown in L medium at 37°C with shaking, and 1 ml of the culture was collected for the sodium bisulfite treatment. After the centrifugation, the supernatant was discarded. The cell pellet was resuspended in 500 μl of an RNA stabilizing reagent (RNA*later* Solution; Thermo Fisher Scientific) and incubated at room temperature for 10 min. After centrifugation and discarding of the supernatant, the cell pellet was frozen in liquid nitrogen and stored at −80°C until use.

### Sodium bisulfite treatment

Aqueous 80% methanol (methanol: water, 80:20 v/v) was directly added to the frozen cell pellet and left for 5 min, and then the cells were collected by centrifugation. After discarding the supernatant, the cells were lysed with lysozyme at concentrations of 0.4 μg/ml in a buffer (10 mM Tris-HCl, pH 8.0, 4 mM EDTA, 4% (w/v) sucrose) for 30 min at 37°C. Sodium bisulfite treatment was carried out using commercially available bisulfite conversion kits (EpiTect Fast Bisulfite Kit and EpiTect Fast LyseAll Bisulfite Kit; Qiagen) according to the manufacturer’s instructions except for the DNA denaturation step and sodium bisulfite reaction time. We carried out the sodium bisulfite reaction at 45°C without DNA denaturation, and incubated it for 120 min. The incubation time was determined according to the CT conversion rates after a series of the sodium bisulfite reactions with incubations ranging from 0 to 180 min (**Figure S8**). After the bisulfite-treated DNA was eluted from the DNA spin column, the DNA was precipitated with ethanol, rinsed with 70% ethanol, resuspended with Milli-Q water, and frozen at −80°C until use.

### PCR amplification of the rDNA and the *sps* segment

To determine CT conversion rates in the whole region of the *rrnI* gene, the region divided into 5’- and 3’-half were separately amplified by PCR using the primers listed in **Table S8**. To amplify the 5’-half of the *rrnI* gene, specific primers that hybridize to the region upstream of the *rrnI* promoter (#1) and the inside of the gene encoding 23S rRNA (#2) were used. To amplify the 3’-half of the *rrnI* gene, specific primers that hybridize to the inside of the gene encoding 23S rRNA (#3) and the spectinomycin-resistance gene (*spc*), which was inserted at the region downstream of the *trnI* gene (#4), were used. To determine CT conversion rates at the *spsAB* genes, specific primers that hybridize to the upstream region of the *spsA* promoter (#5) and the inside of the *spsB* gene (#6) were used. The amplified segments are shown in **Figure S9**.

### Library preparation and sequencing with MiSeq

To amplify the regions for sequencing, the target DNA fragments were amplified by PCR (TaKaRa EpiTaq HS; TAKARA). The PCR fragment was purified by precipitation with PEG 8000. The concentration of the DNA fragments was quantified by using a fluorescence dye and a fluorometer (Qubit dsDNA BR assay kit; Thermo Fisher Scientific). The amplified DNA fragments (100 ng) were sheared to the average length of 250 bp by using an ultrasonicator (Covaris focused-ultrasonicator S220; Covaris) and purified by using magnetic beads (Agencourt AMPure XP; Beckman Coulter). Adaptor and index primers were added to the sheared DNA fragments using a next-generation sequencing library preparation kit (NEBNext Ultra DNA Library Prep Kit for Illumina; NEB) according to the manufacturer’s instructions. PCR enrichment was carried out with nine cycles. The index-ligated fragments were mixed to construct a sequencing library and sequenced using the MiSeq system and its reagent kit (MiSeq Reagent Kits v2 (500 cycles); Illumina) according to the manufacturer’s instructions. Sequencing was carried out with 250 cycles and a paired-end mode.

### Determination of the parameters for mapping of the sequenced reads

Sequenced reads were first trimmed using a trimming tool, Trimmomatic (v0.33; Bolger et al., 2014). To trim the reads, the following parameters were used: accepted mismatches, 2; palindrome clip threshold, 30; simple clip threshold, 10; leading, 20; trailing, 20; window size, 4; average quality, 15; minimal length, 30. The trimmed reads were mapped to the sequence of the 5’ half of *rrnI* as a reference by using mapping software, Bismark (v0.20.0; Kruger and Andrews, 2011). The version of Bowtie2, the software required to run Bismark, was v2.2.6 (Langmead and Salzberg, 2012). In order to map as many reads as possible, we determined parameters for the minimum alignment function of Bismark (score-min option). The option consists of three parameters: (a) a function type, (b) a constant term, X, and (c) a coefficient, Y. We used a linear equation (L) for the function type and ran Bismark with score--min L, -X, and -Y, where X and Y were changed from 0 to 150 and 0 to 4, respectively. While Y was being changed, X was fixed as 0. While X was being changed, Y was fixed as 2. The mapping efficiencies were plotted against the increasing values of X or Y (**Figure S10**). The values of X and Y required to map as many reads as possible were determined as 100 and 2, respectively.

### Quality control of the mapped reads

Mapped reads below MAPQ⇐ one were removed using SAMtools (v1.9; Li et al., 2009), and reads with insertions and deletions were removed using an in-house Perl script pid3.pl.

### Calculation of the CT conversion rates

Depths of the coverage for each nucleotide position in the mapped reads were counted for four bases (A, T, G, C) by using SAMtools. CT conversion rates were calculated for each cytosine on both the top and bottom strands according to the following equations: CT conversion rates for top strand (%) = (Counts for T) / (Counts for C + Counts for T) × 100; CT conversion rates for bottom strand (%) = (Counts for A) / (Counts for G + Counts for A) × 100.

### Quantification and Statistical Analysis

The mean value and standard deviations of the mean were calculated. Simple regression analysis was done to draw regression lines in the correlation plots between cytosine contents and CT conversion rates. The equations of the regression line for the top and bottom strand are y = 22.405× + 1.9114 and y = 18.677× + 0.8789, respectively, where x is the cytosine content and y is the CT conversion rate.

## Notes

### Competing Interest Statement

The authors have declared no competing interest.

