## Supplementary material for "Profiling a single-stranded DNA region within an rDNA segment that is a loading site for bacterial condensin": Yano et al_supplemental files: Yano et al_FigureS2.pdf

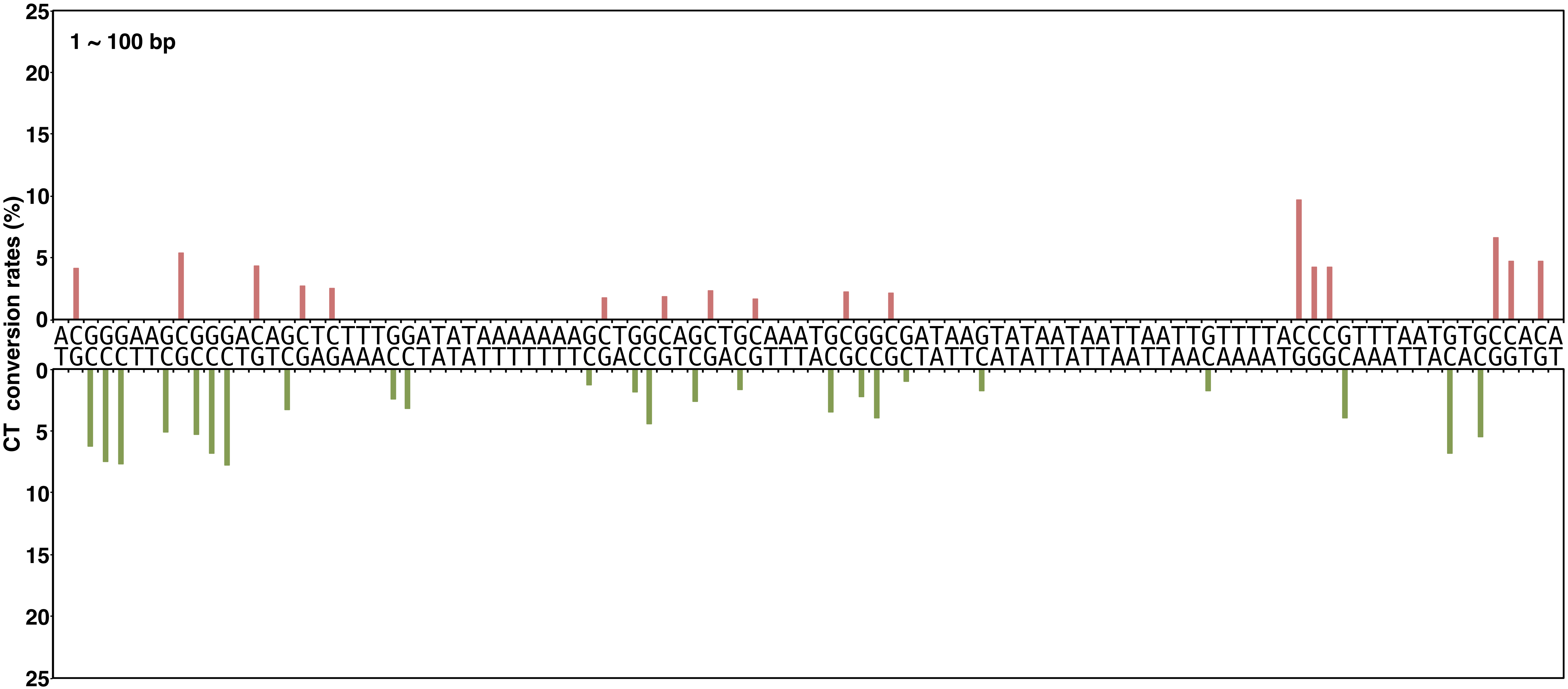


**Yano, Noguchi, and Niki Figure S2**

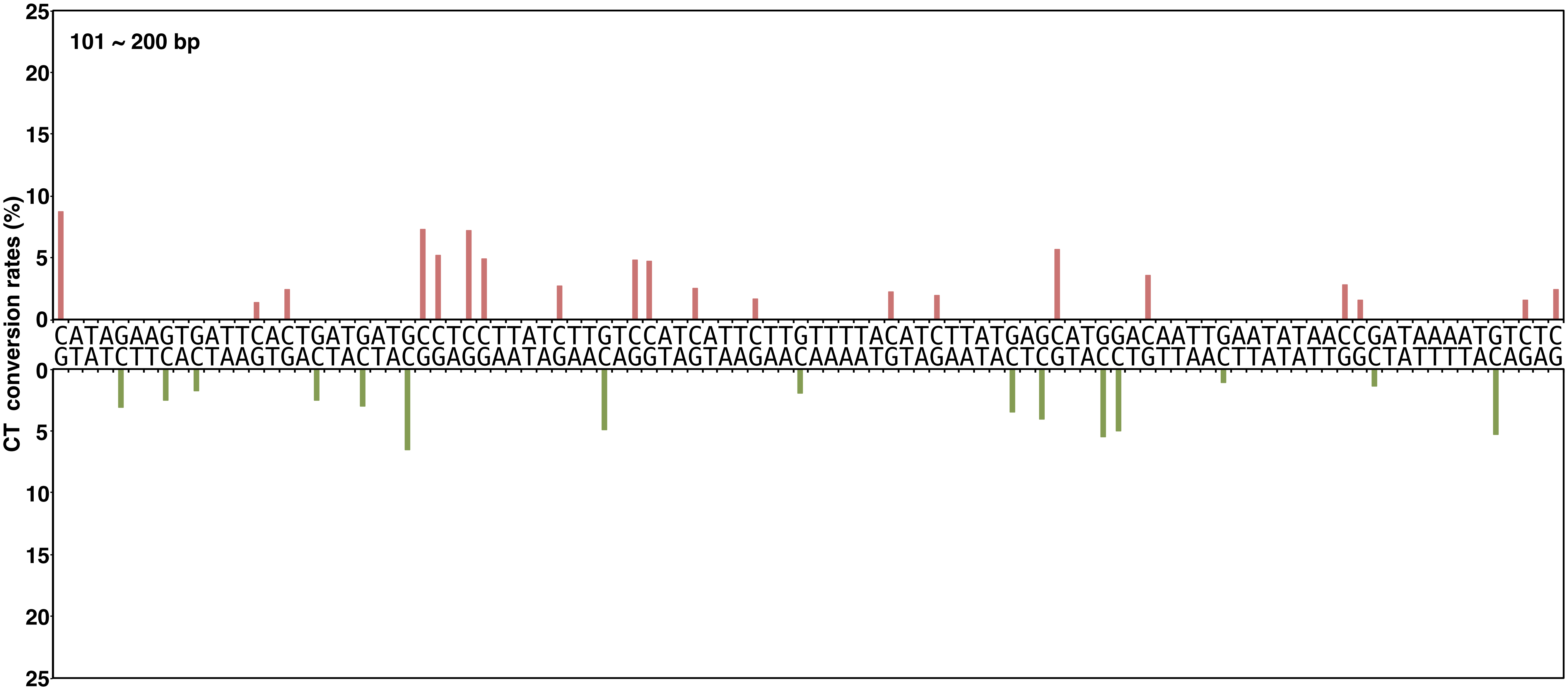


**Yano, Noguchi, and Niki Figure S2, continued**

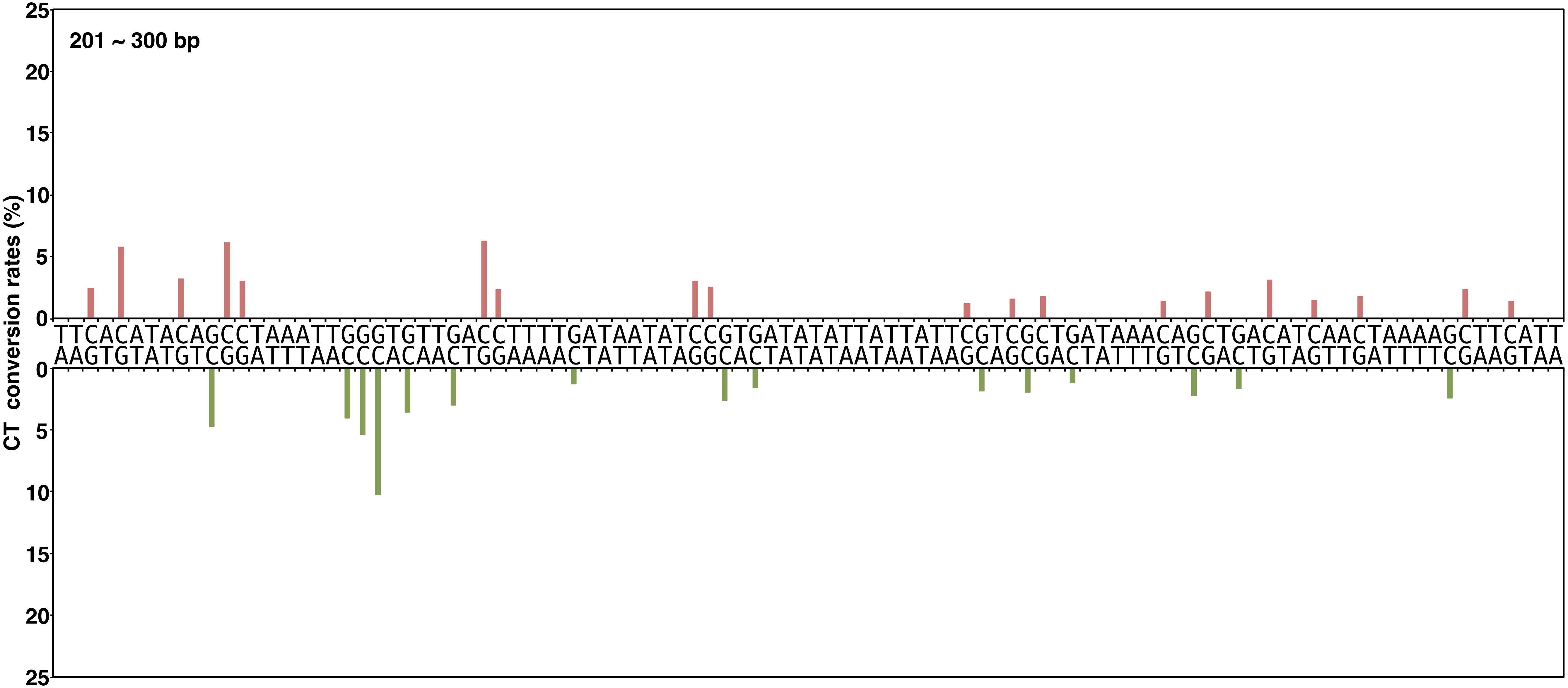


**Yano, Noguchi, and Niki Figure S2, continued**

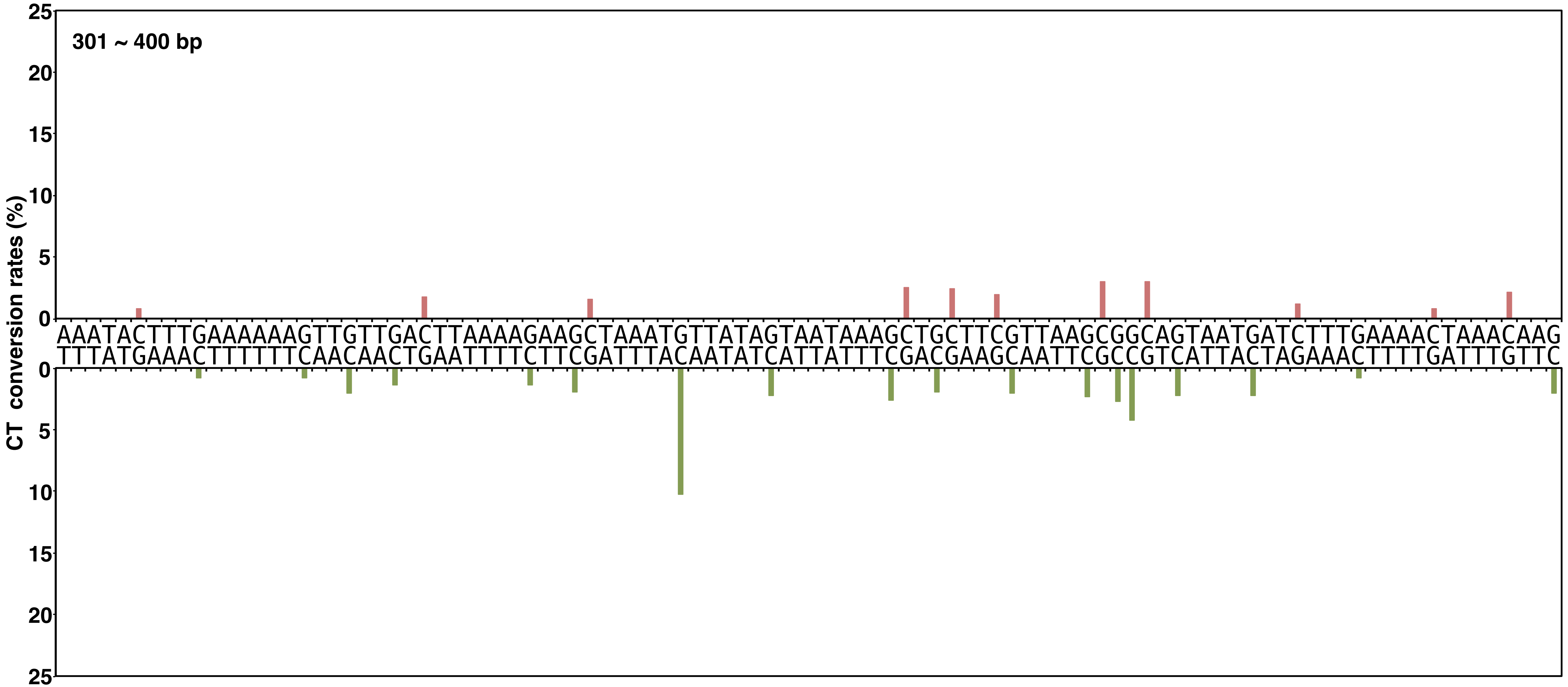


Yano, Noguchi, and Niki Figure S2, continued

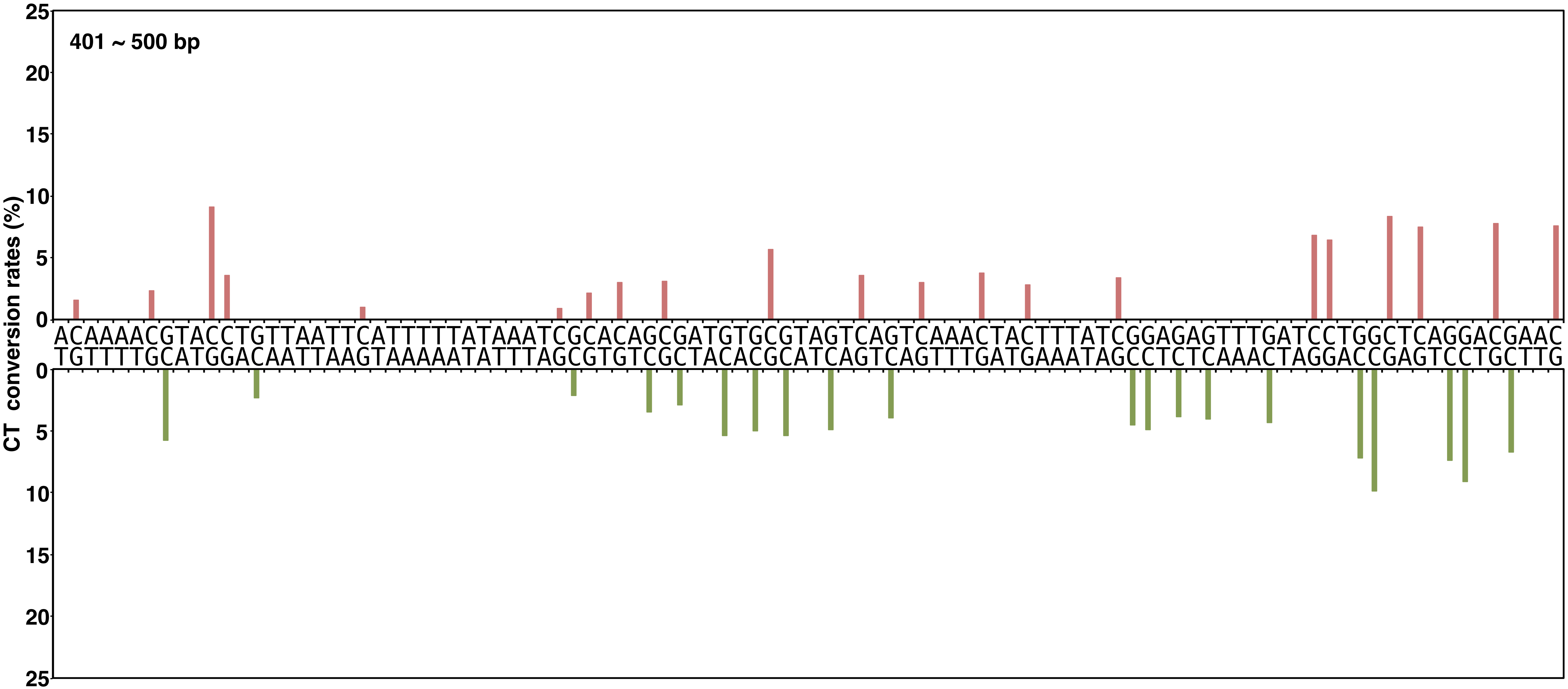


**Yano, Noguchi, and Niki Figure S2, continued**

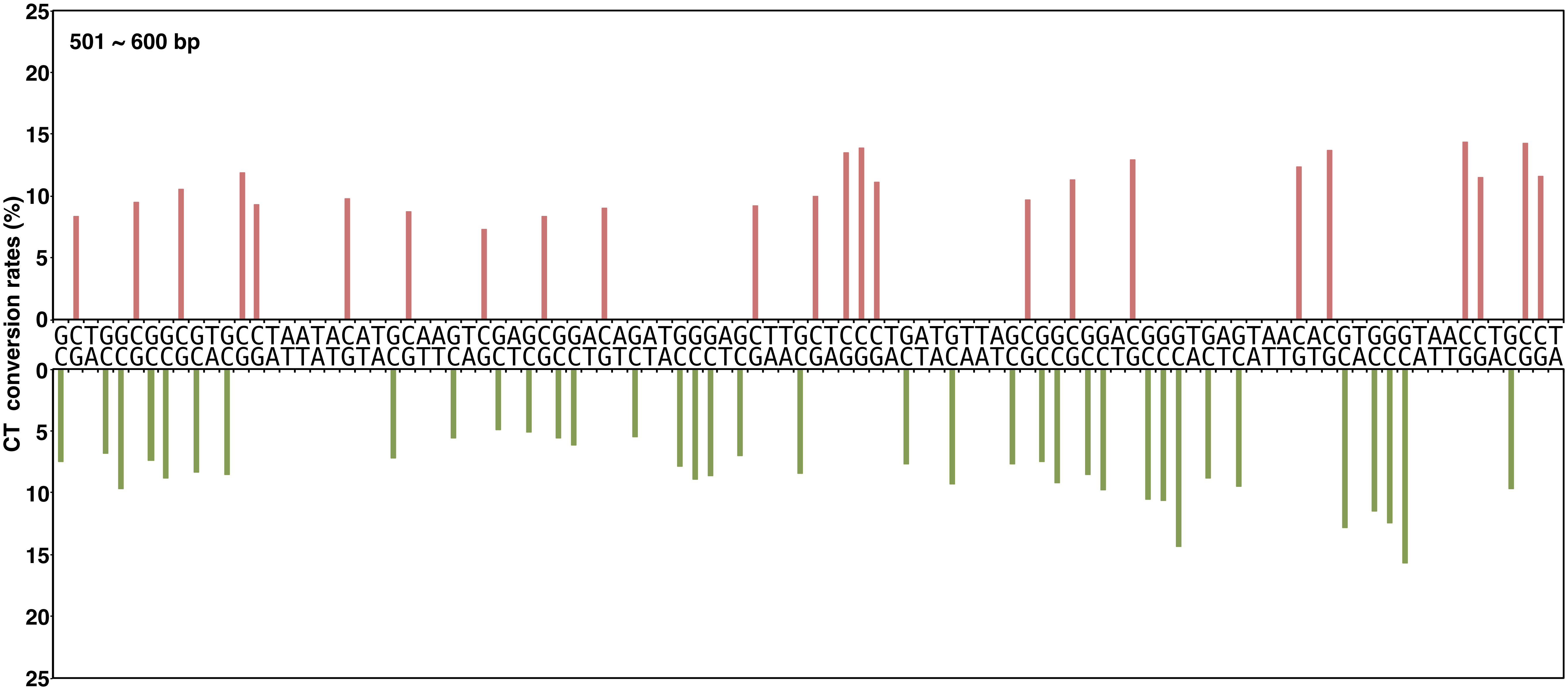


**Yano, Noguchi, and Niki Figure S2, continued**

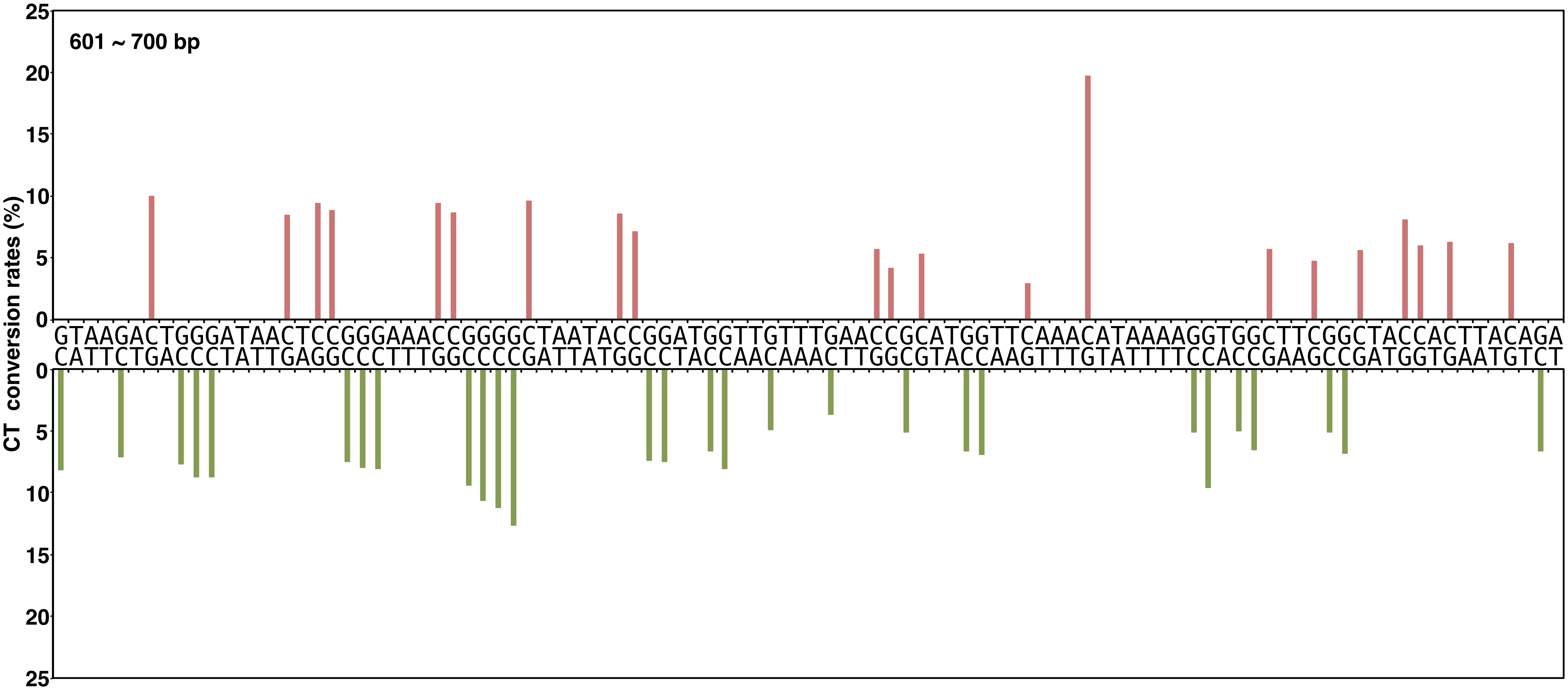


**Yano, Noguchi, and Niki Figure S2, continued**

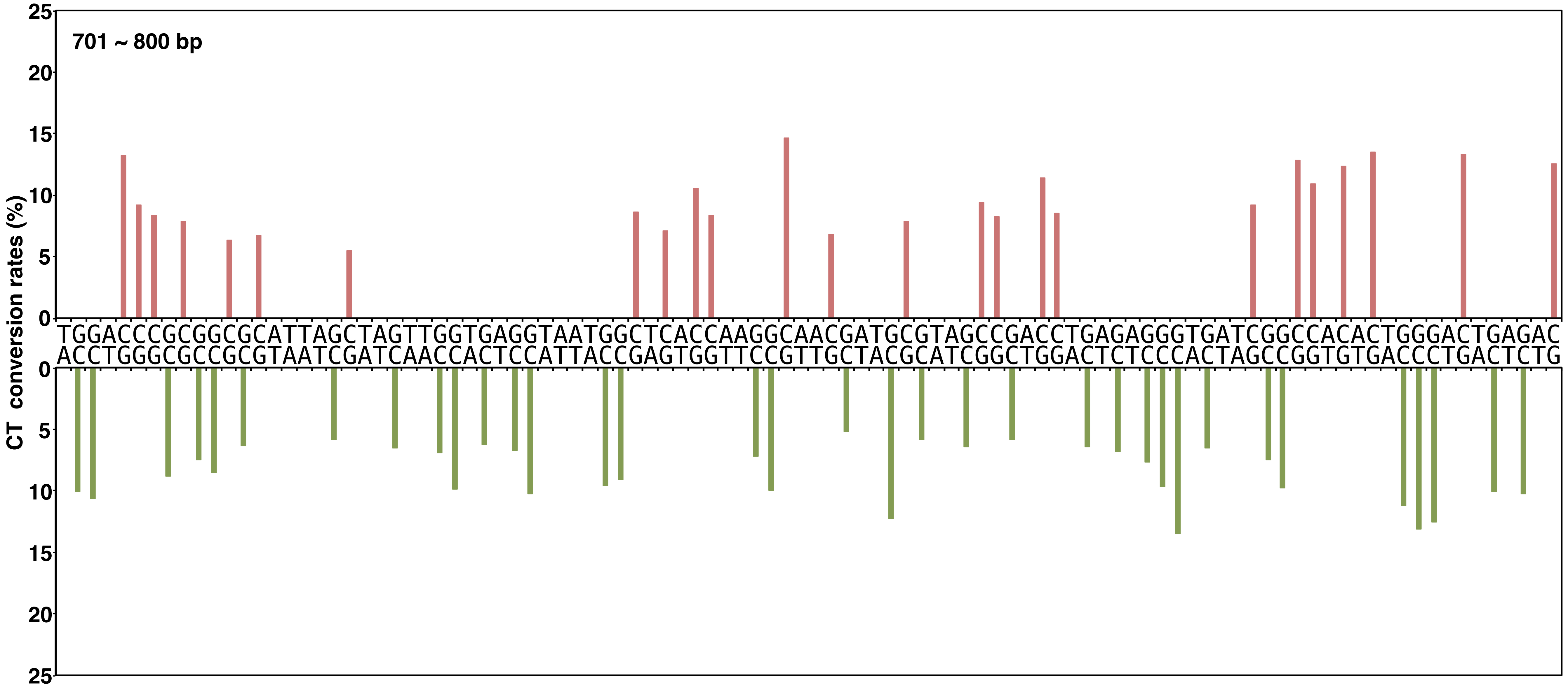


**Yano, Noguchi, and Niki Figure S2, continued**

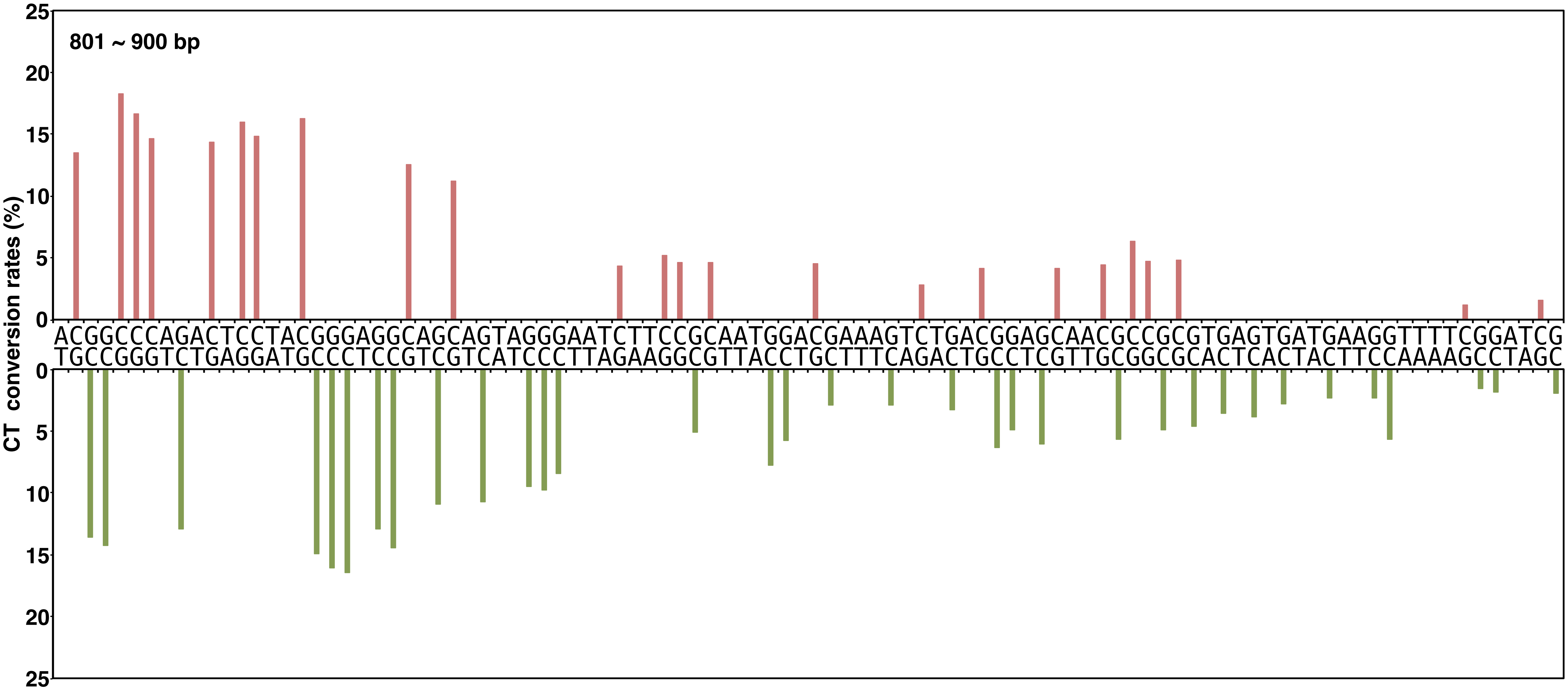


**Yano, Noguchi, and Niki Figure S2, continued**

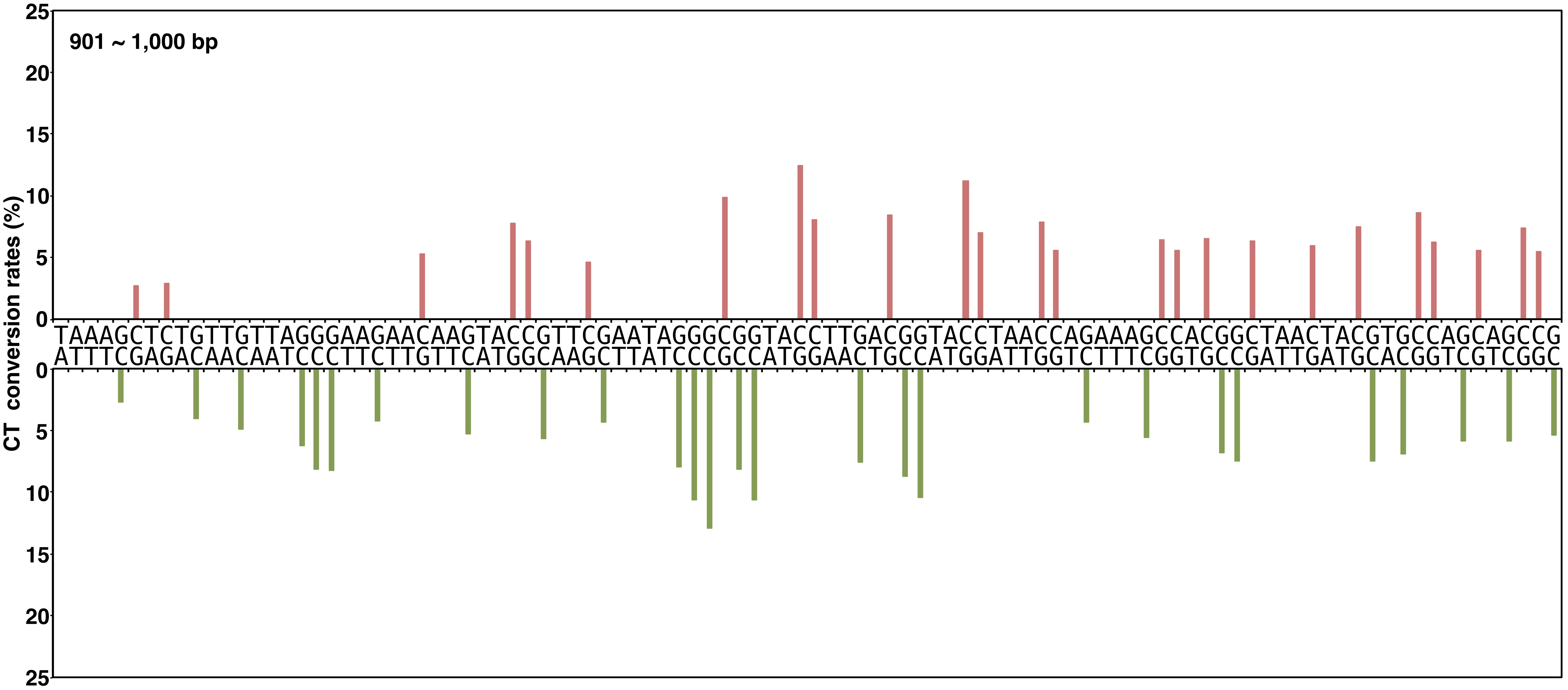


**Yano, Noguchi, and Niki Figure S2, continued**

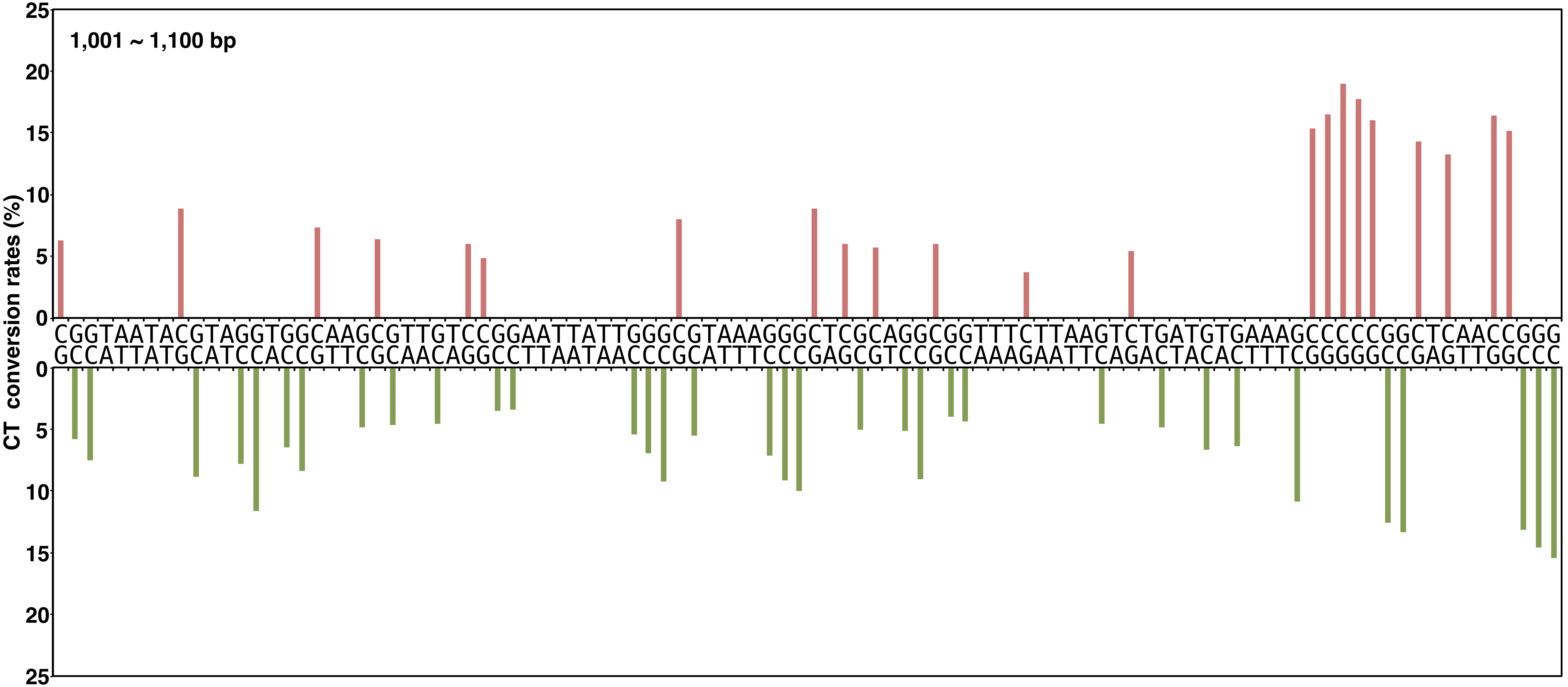


**Yano, Noguchi, and Niki Figure S2, continued**

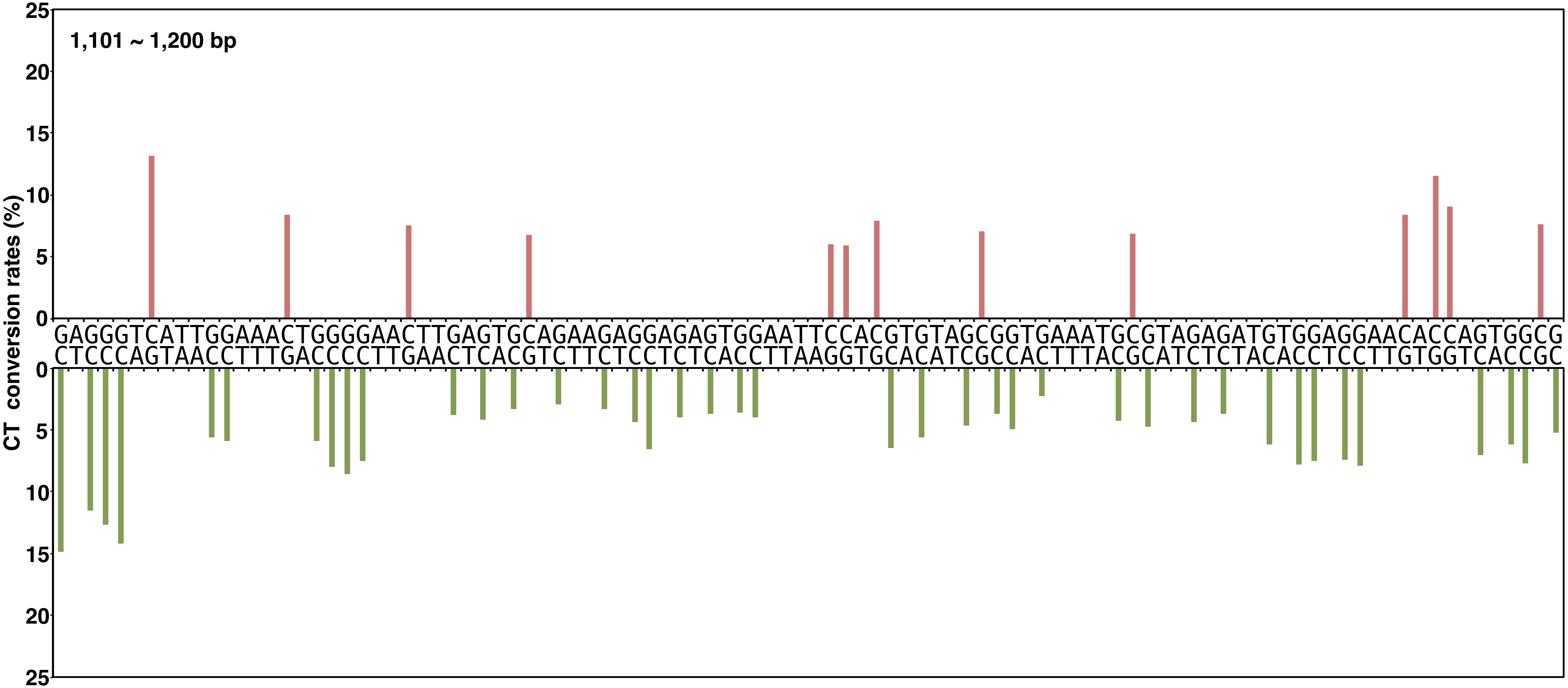


**Yano, Noguchi, and Niki Figure S2, continued**

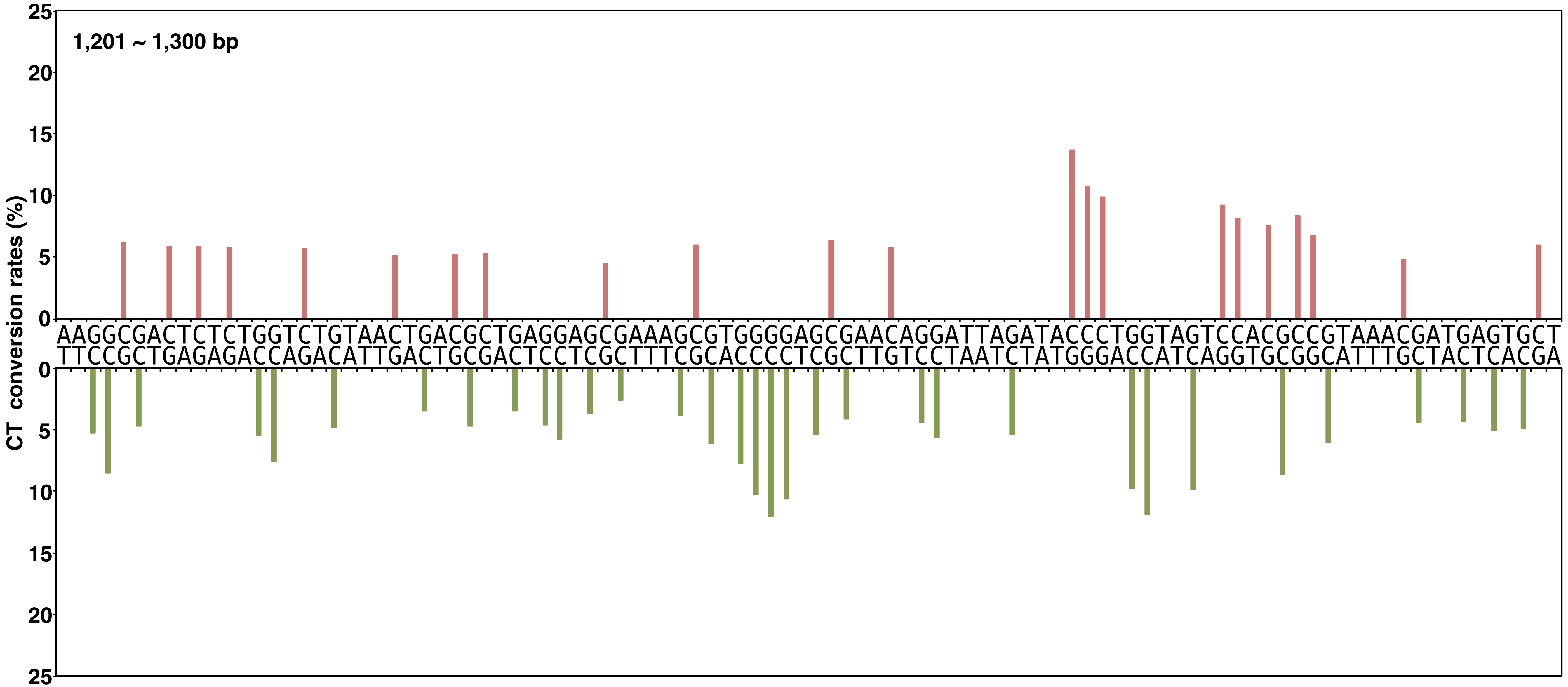


**Yano, Noguchi, and Niki Figure S2, continued**

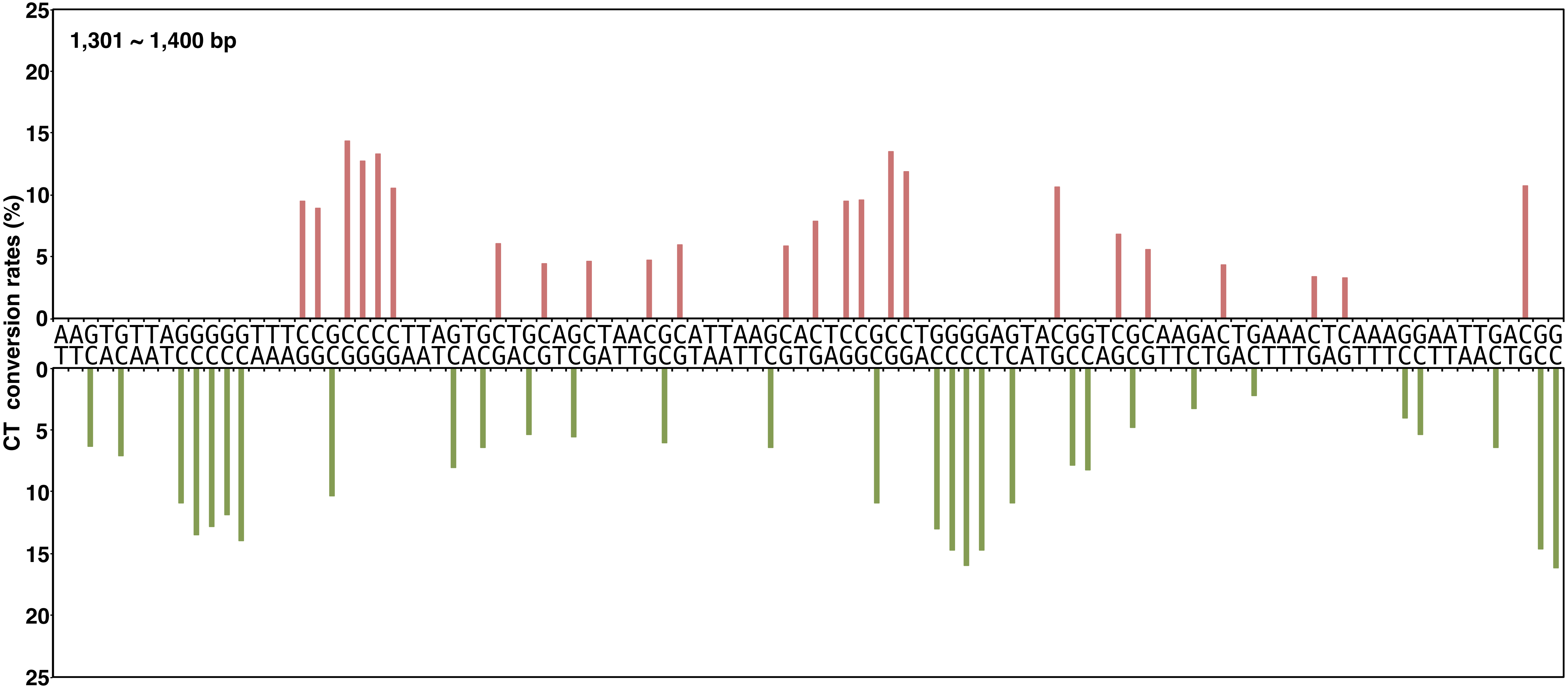


**Yano, Noguchi, and Niki Figure S2, continued**

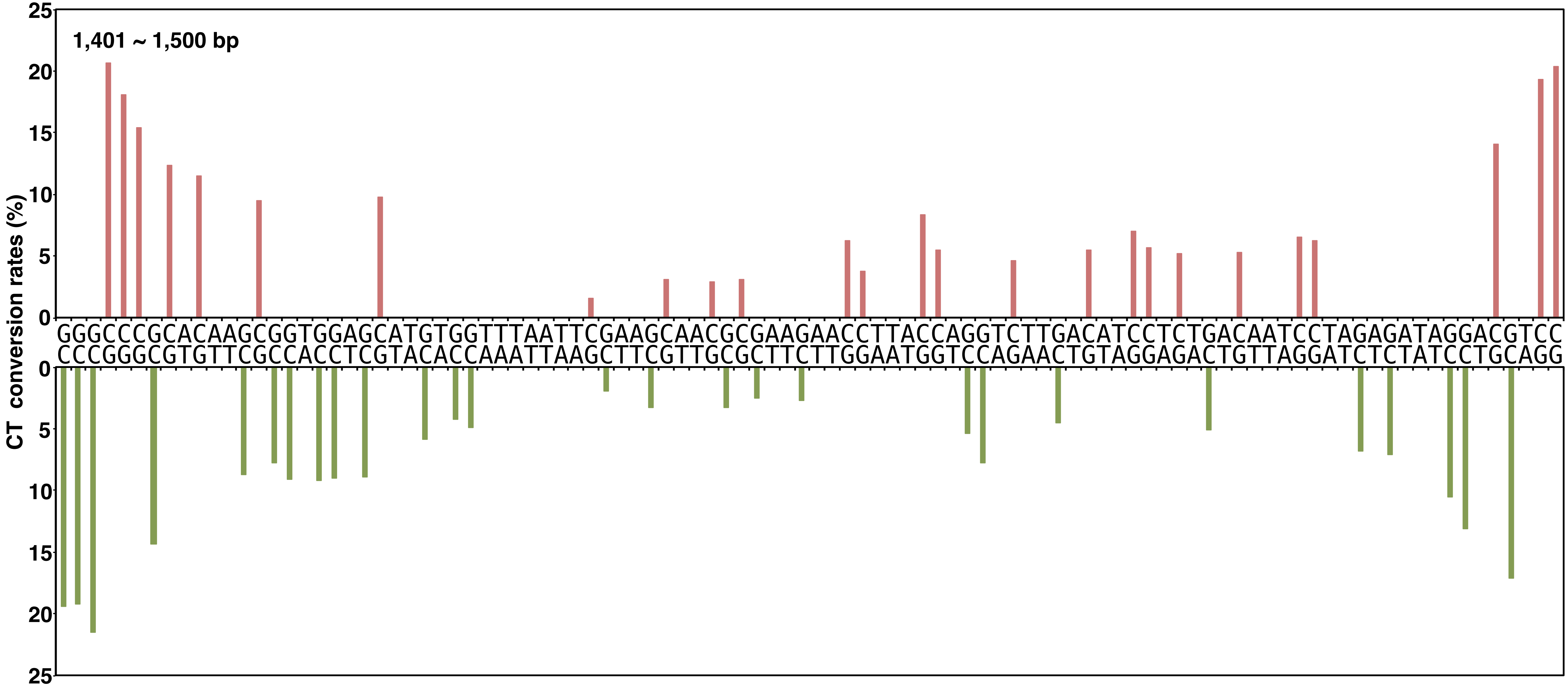


**Yano, Noguchi, and Niki Figure S2, continued**

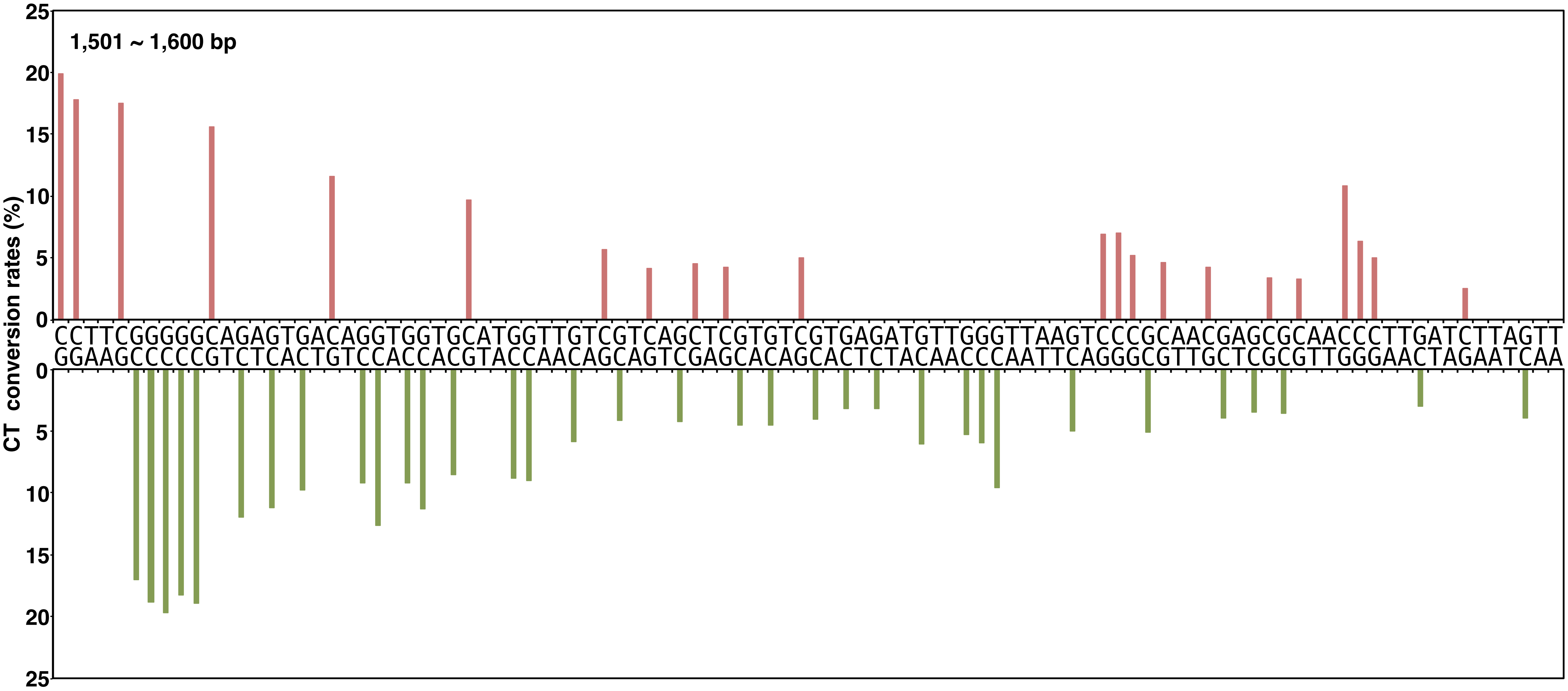


**Yano, Noguchi, and Niki Figure S2, continued**

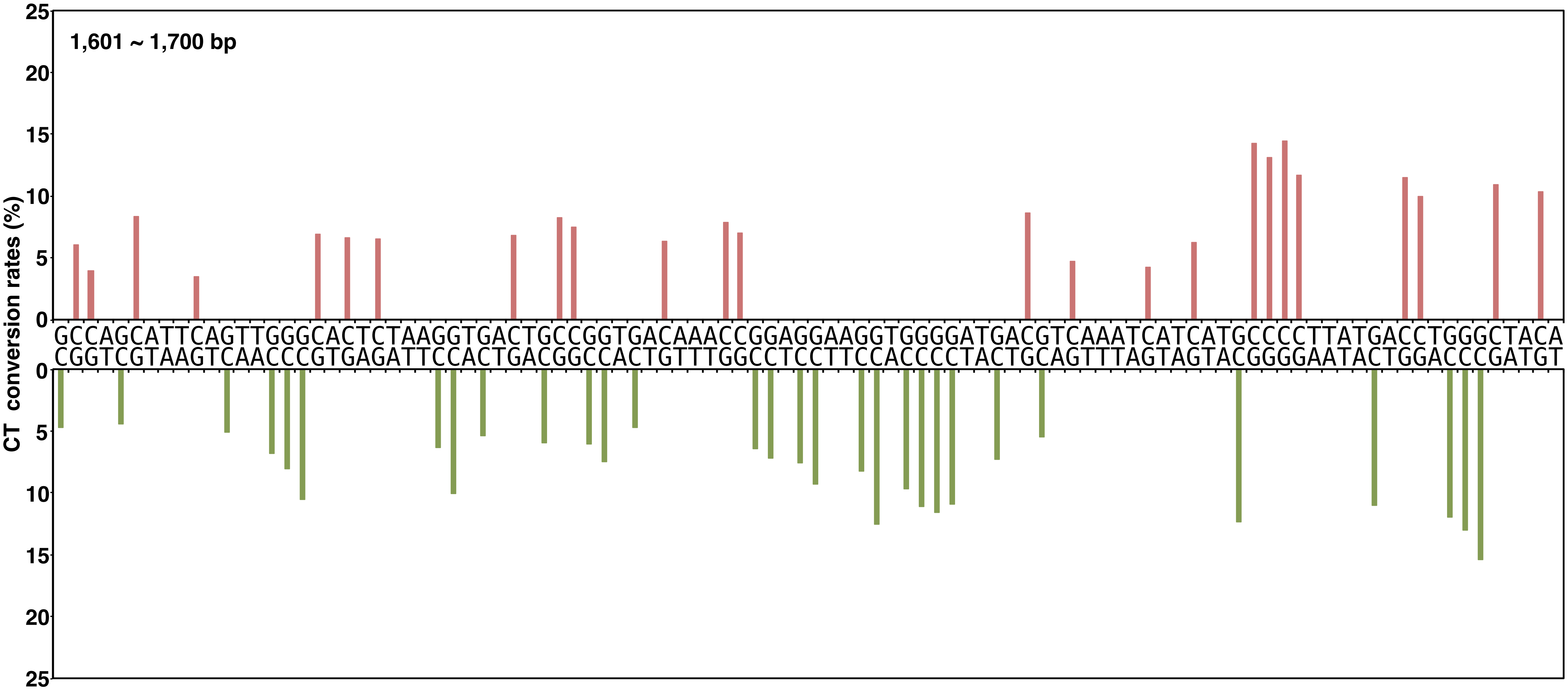


**Yano, Noguchi, and Niki Figure S2, continued**

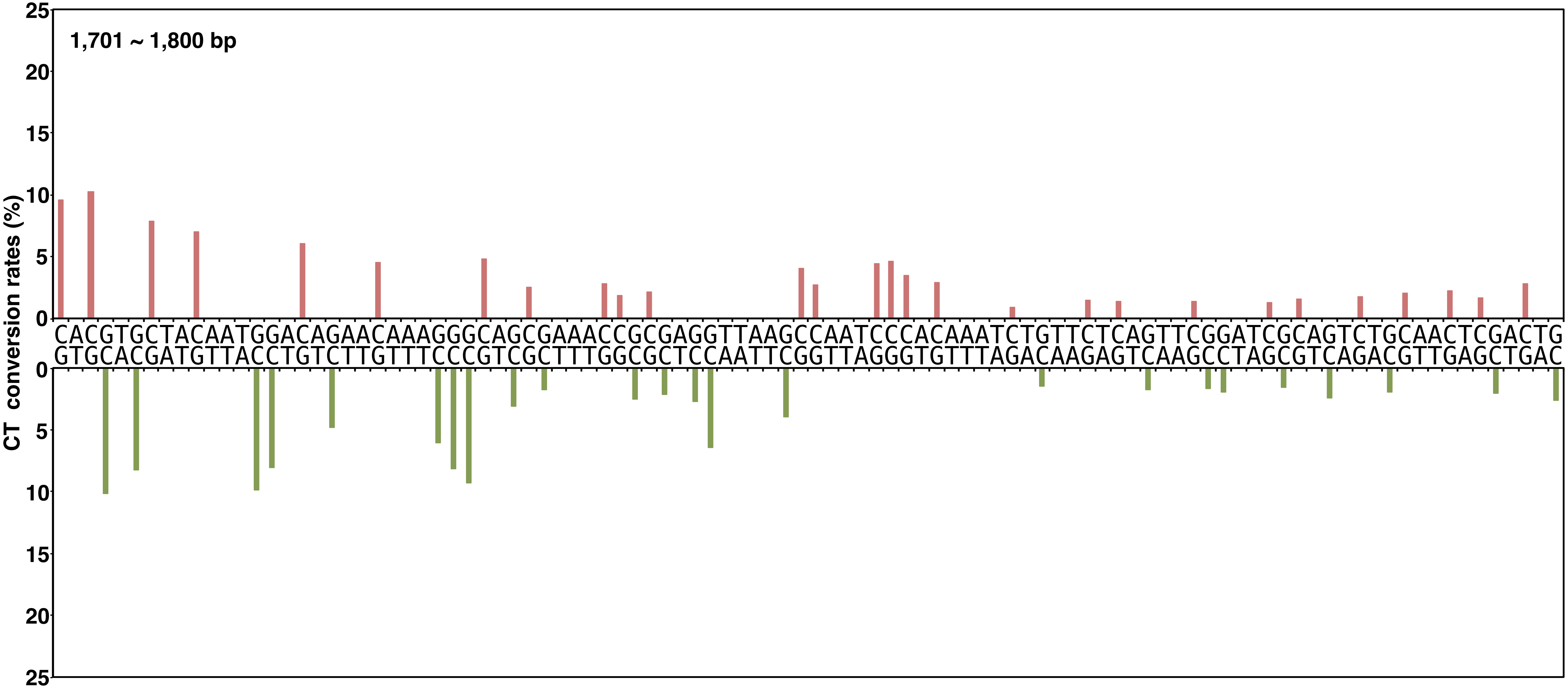


**Yano, Noguchi, and Niki Figure S2, continued**

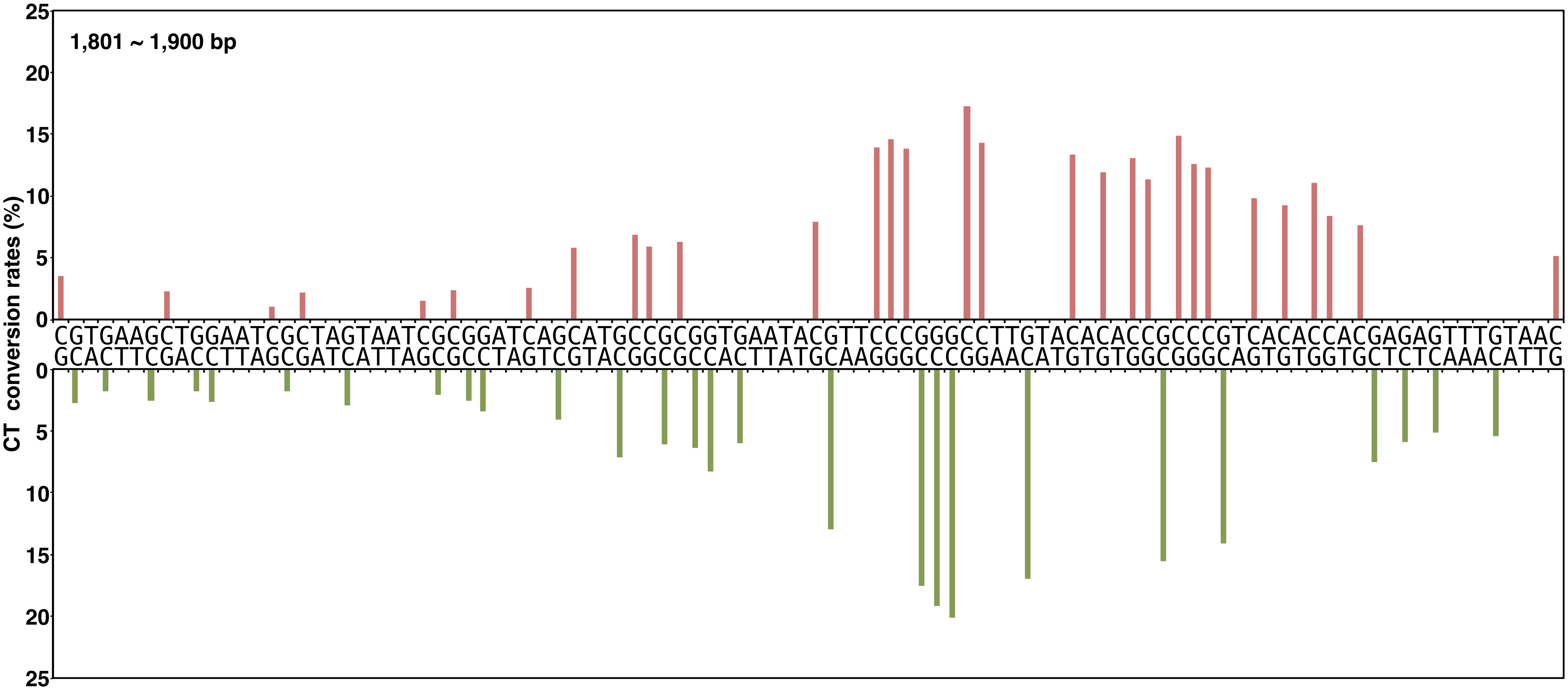


**Yano, Noguchi, and Niki Figure S2, continued**

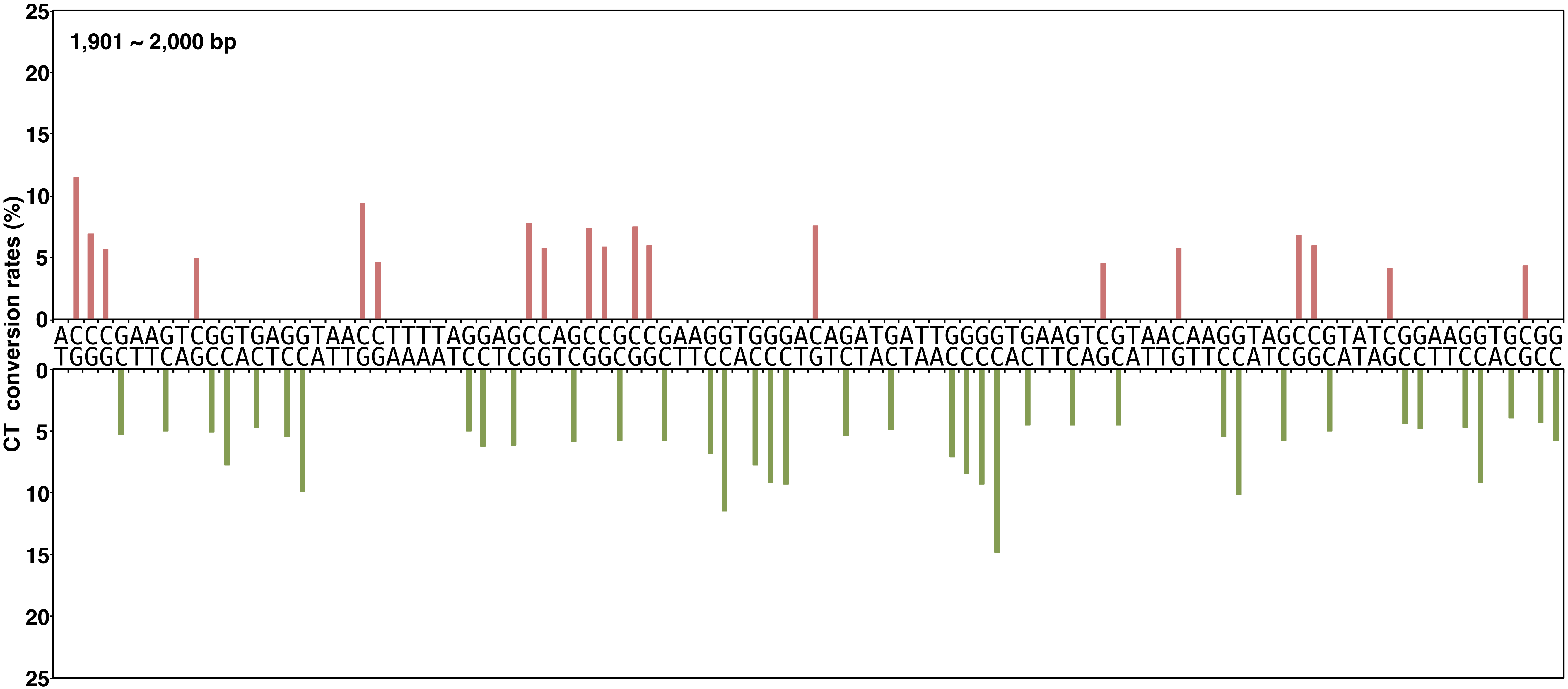


**Yano, Noguchi, and Niki Figure S2, continued**

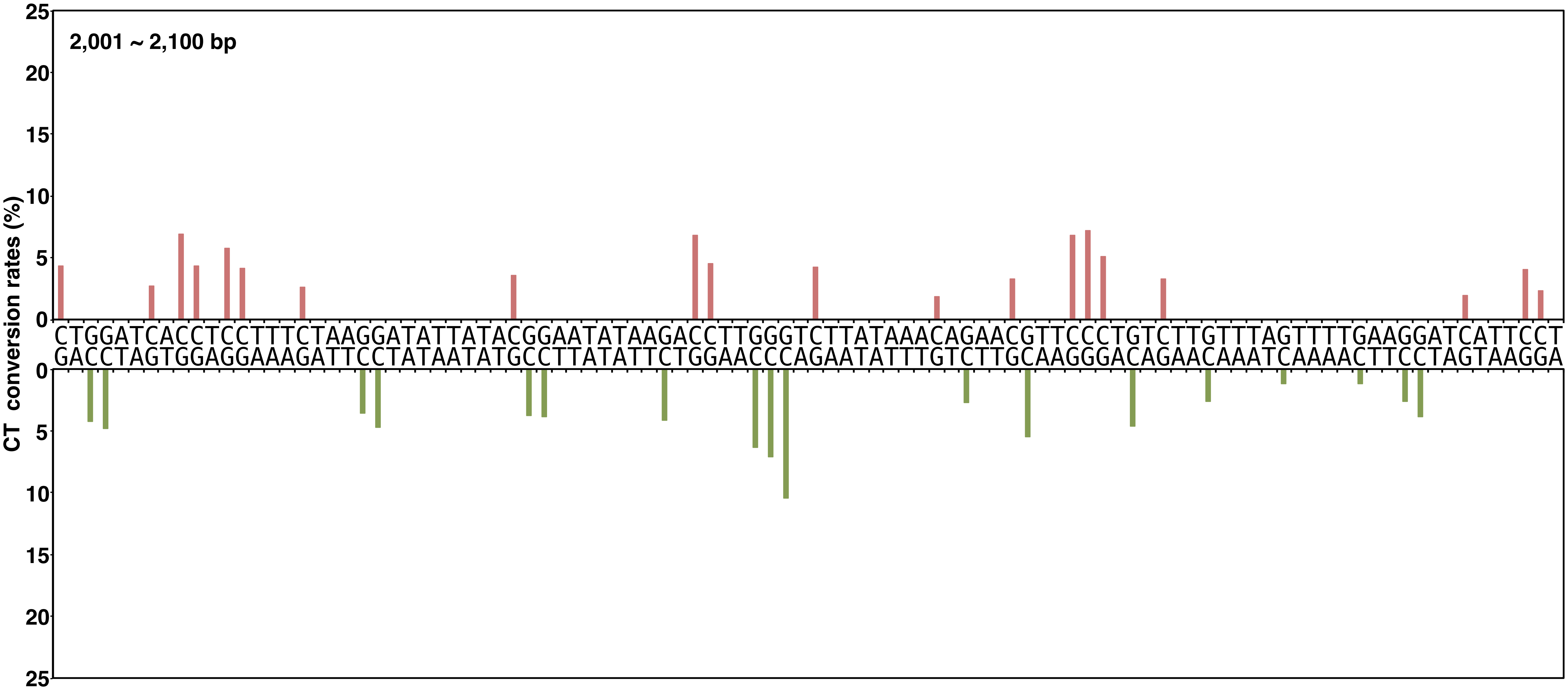


**Yano, Noguchi, and Niki Figure S2, continued**

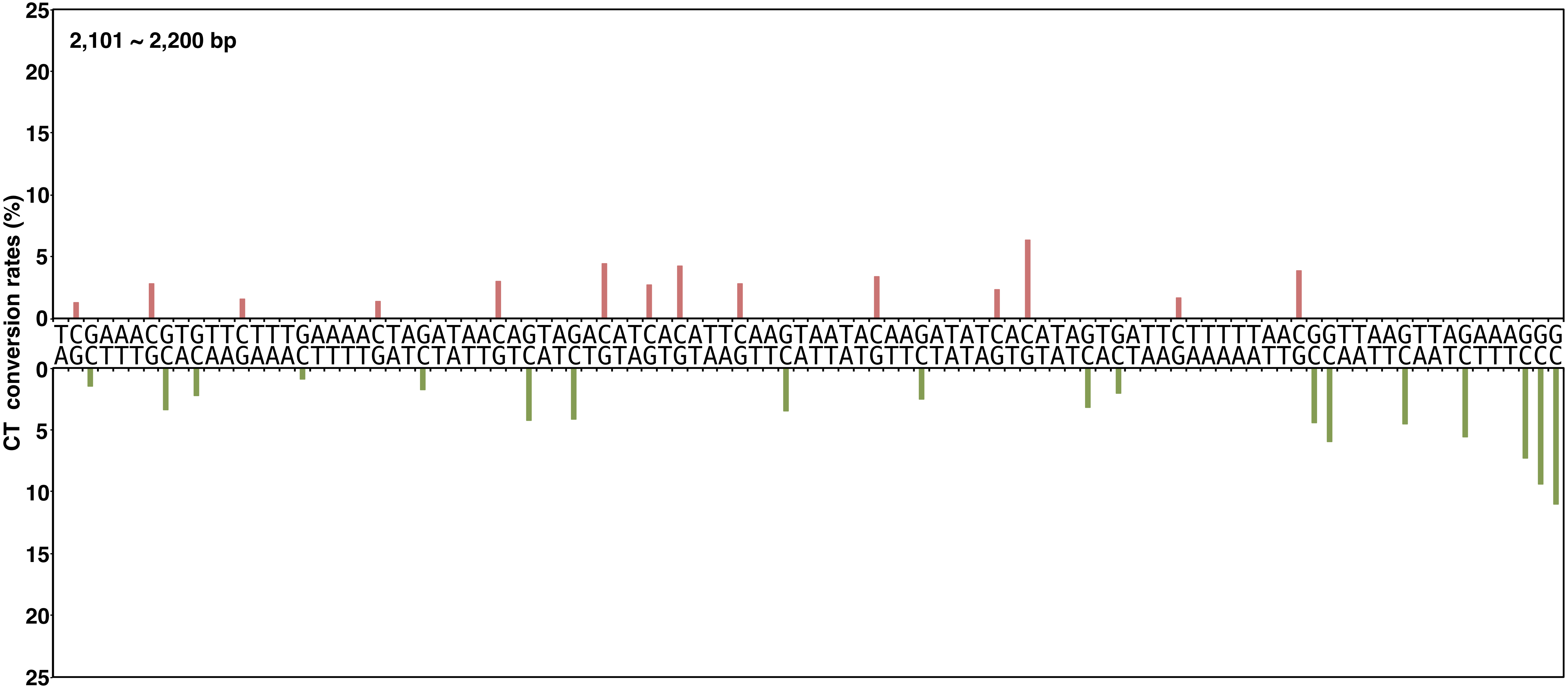


**Yano, Noguchi, and Niki Figure S2, continued**

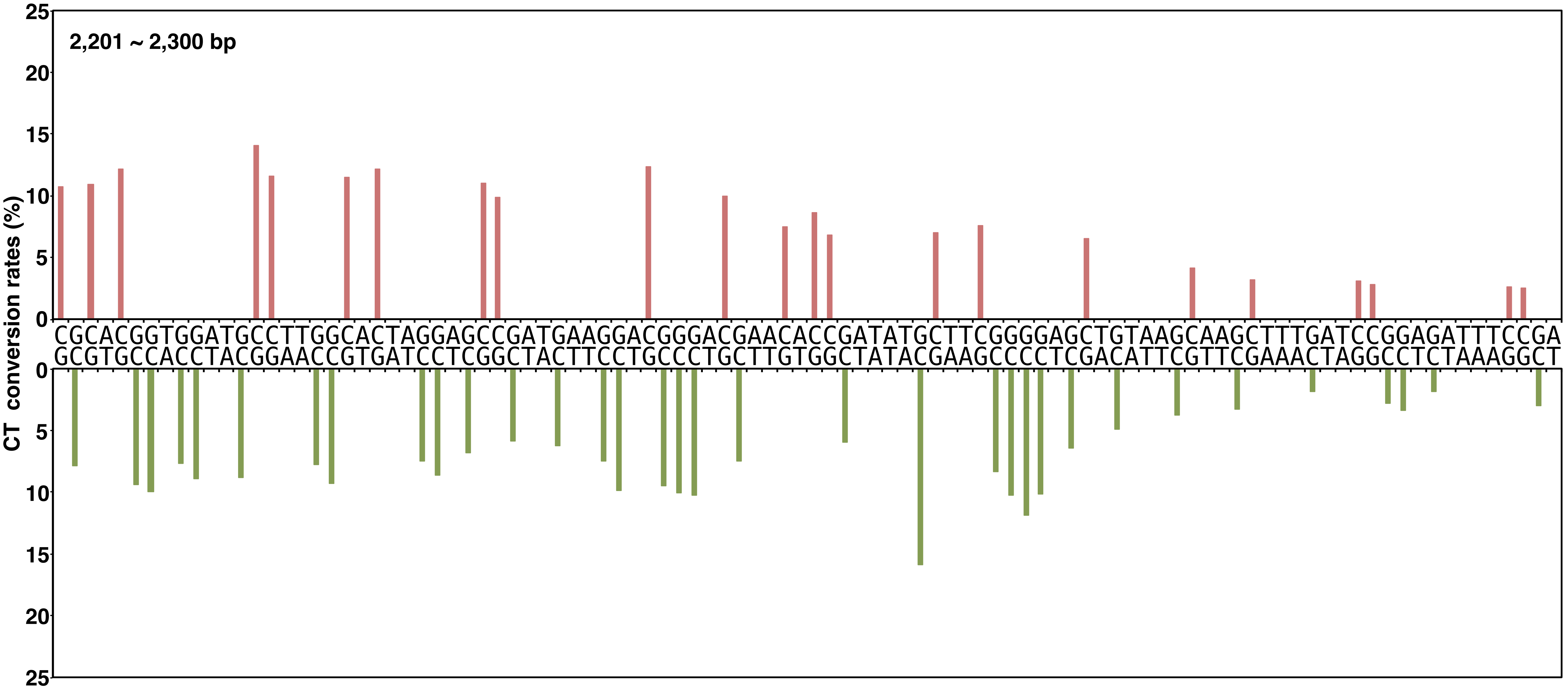


**Yano, Noguchi, and Niki Figure S2, continued**

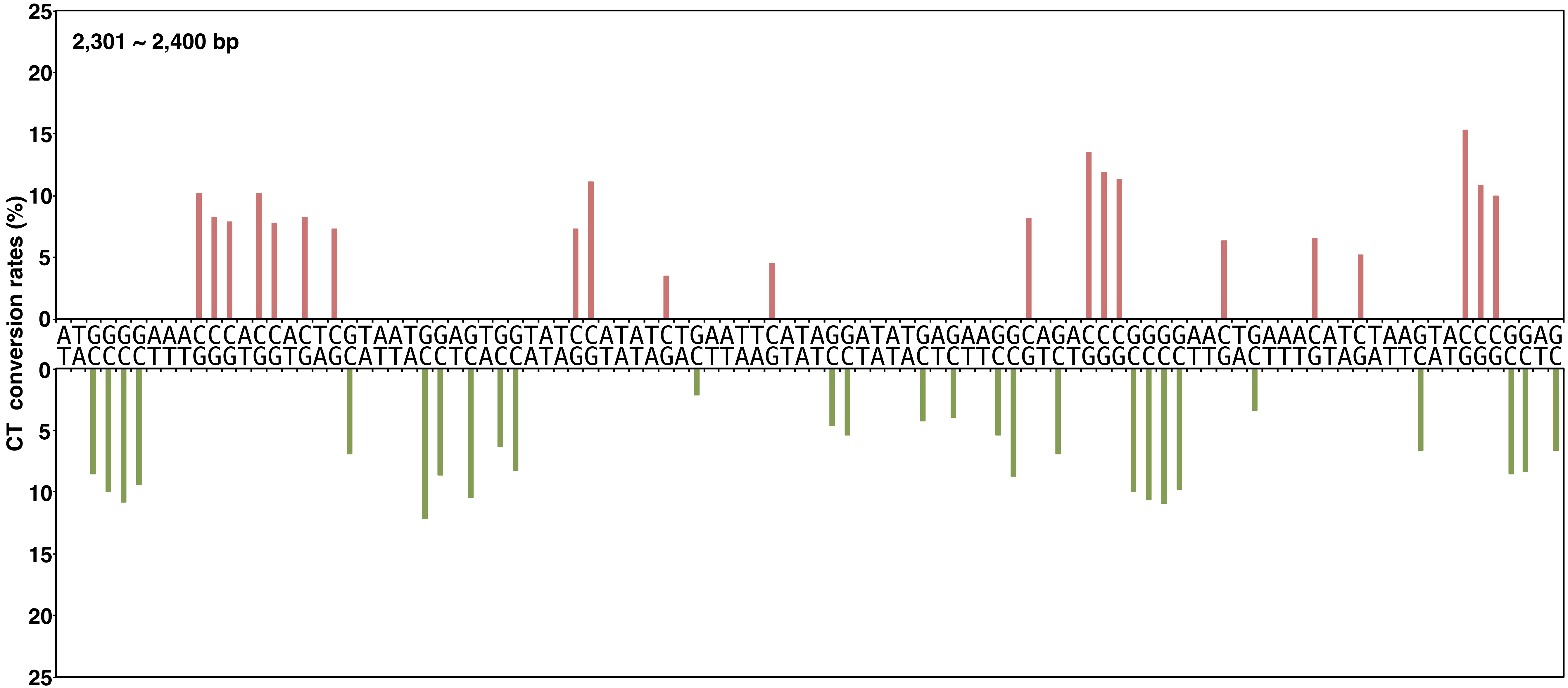


**Yano, Noguchi, and Niki Figure S2, continued**

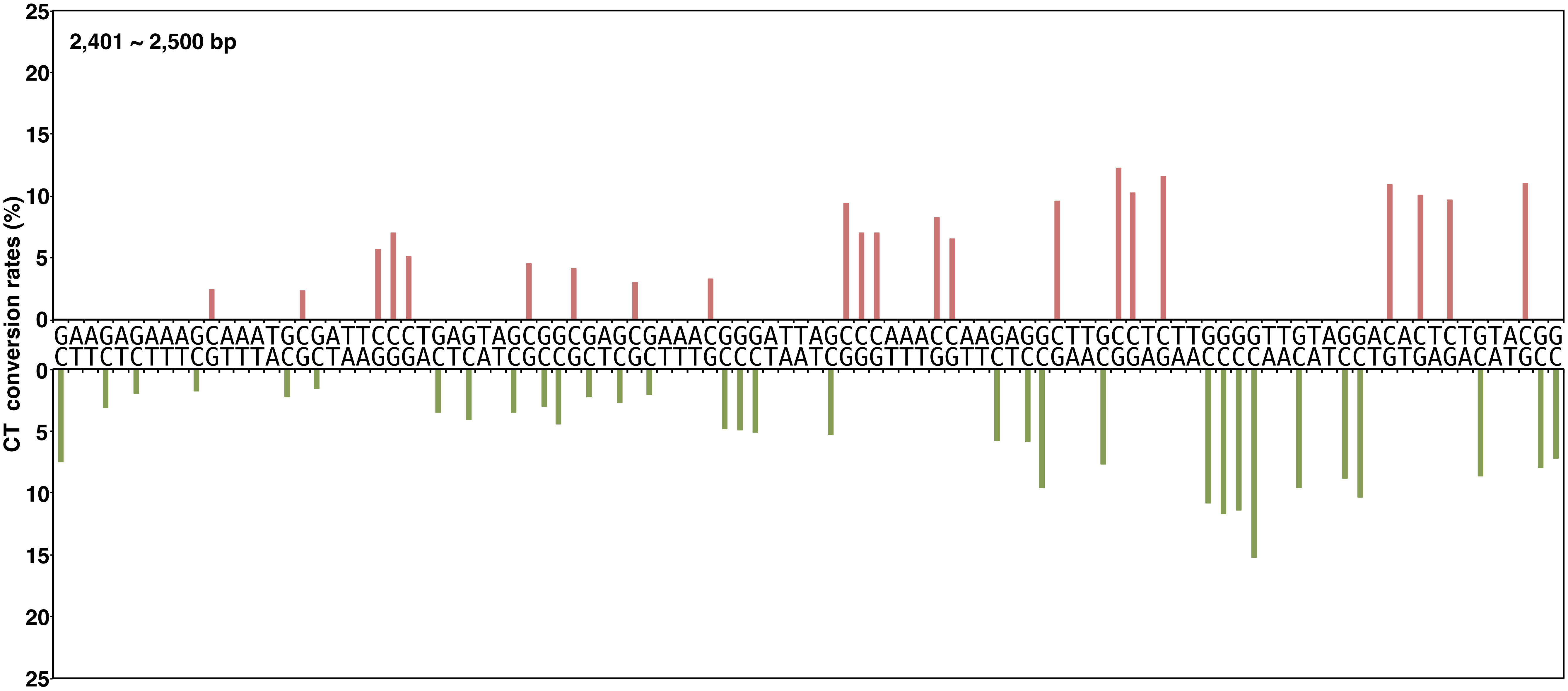


**Yano, Noguchi, and Niki Figure S2, continued**

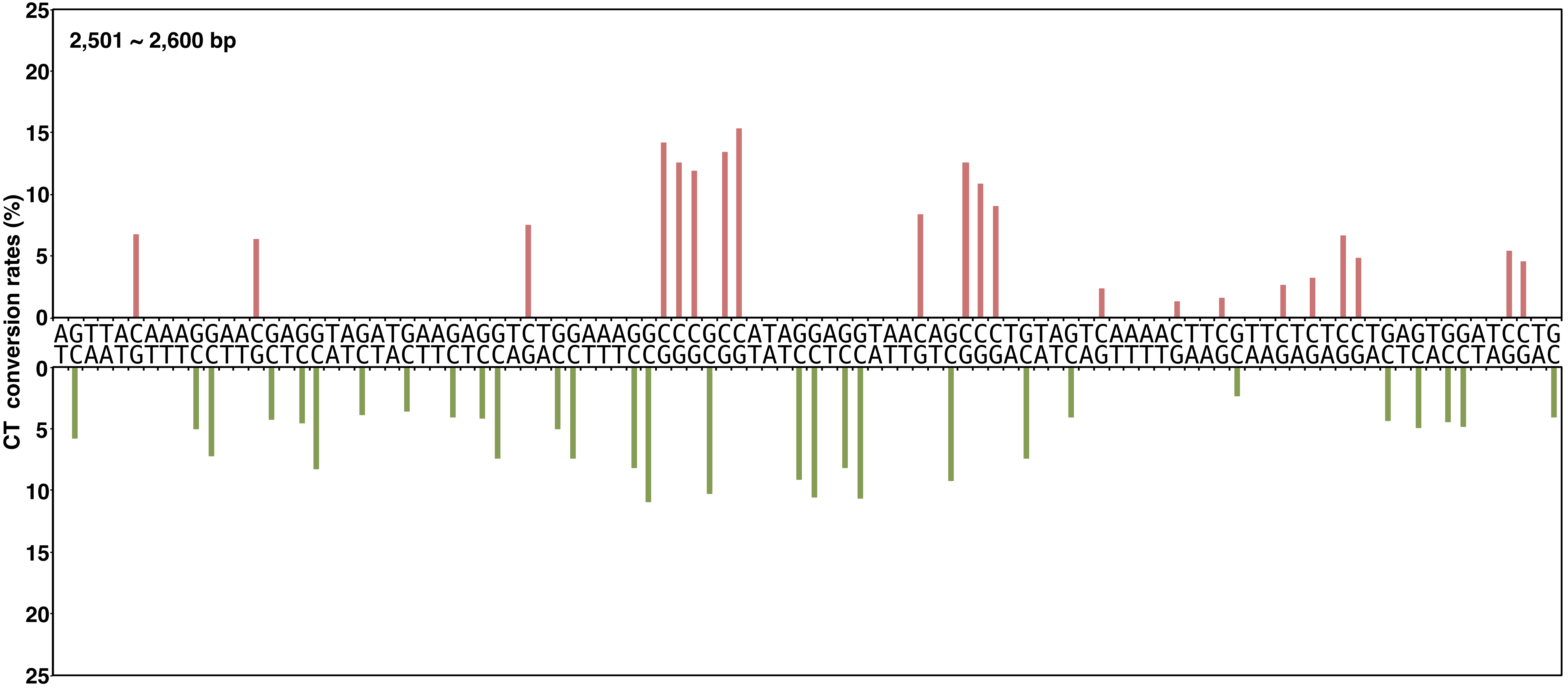


**Yano, Noguchi, and Niki Figure S2, continued**

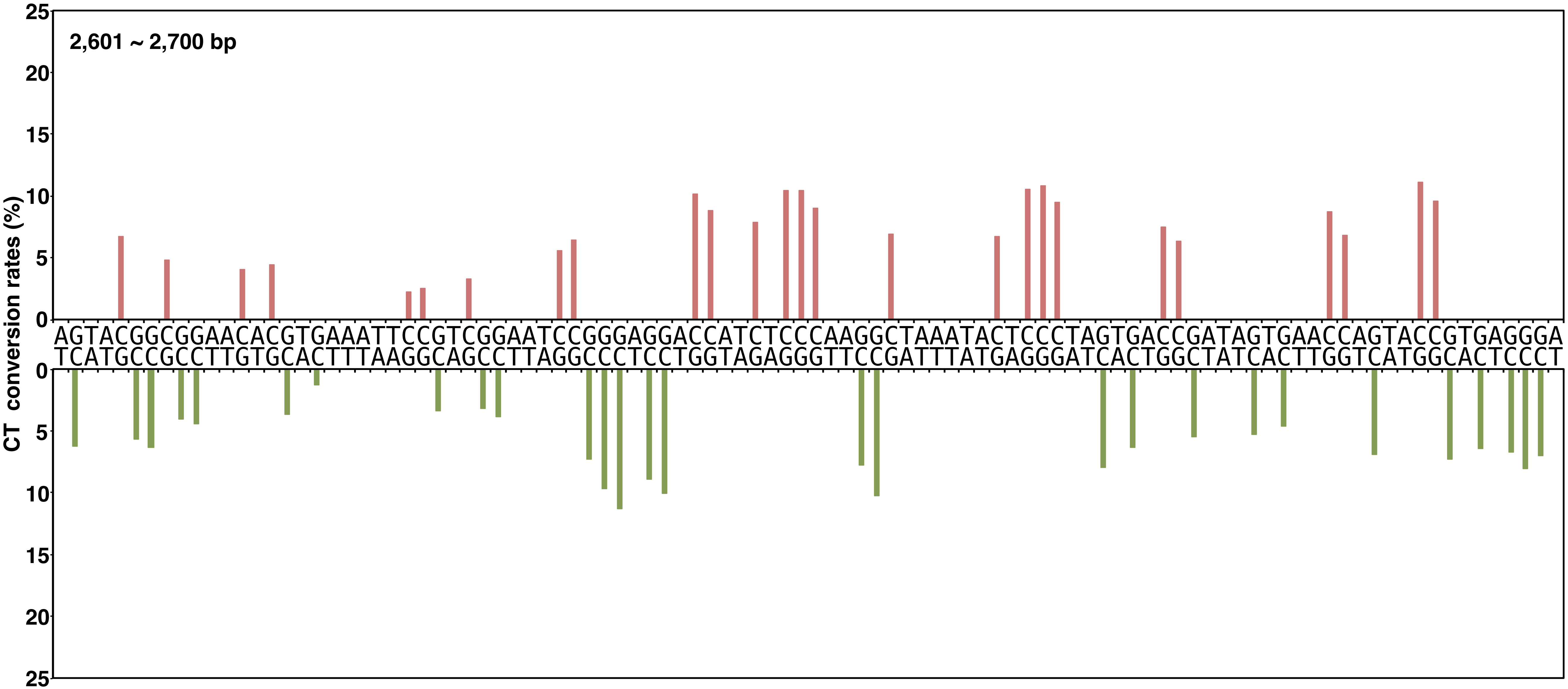


**Yano, Noguchi, and Niki Figure S2, continued**

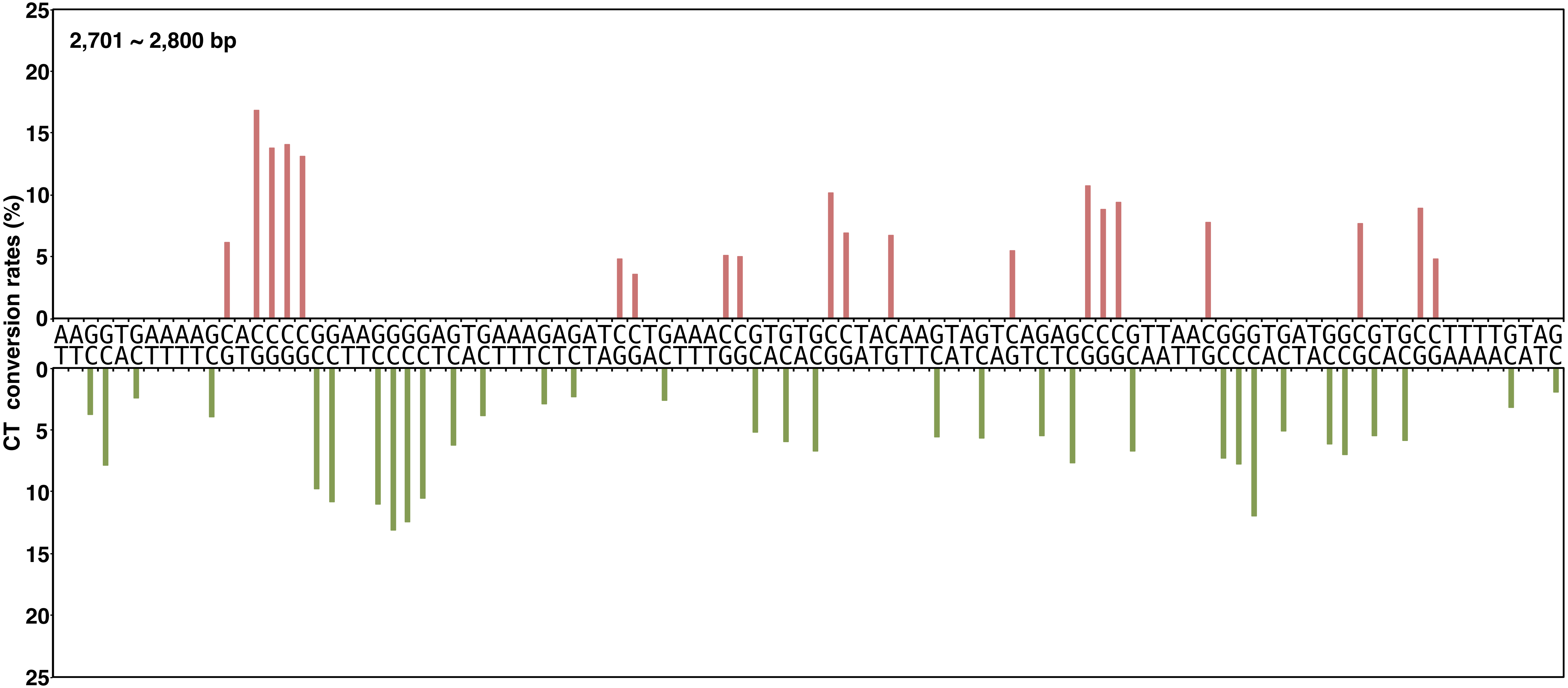


**Yano, Noguchi, and Niki Figure S2, continued**

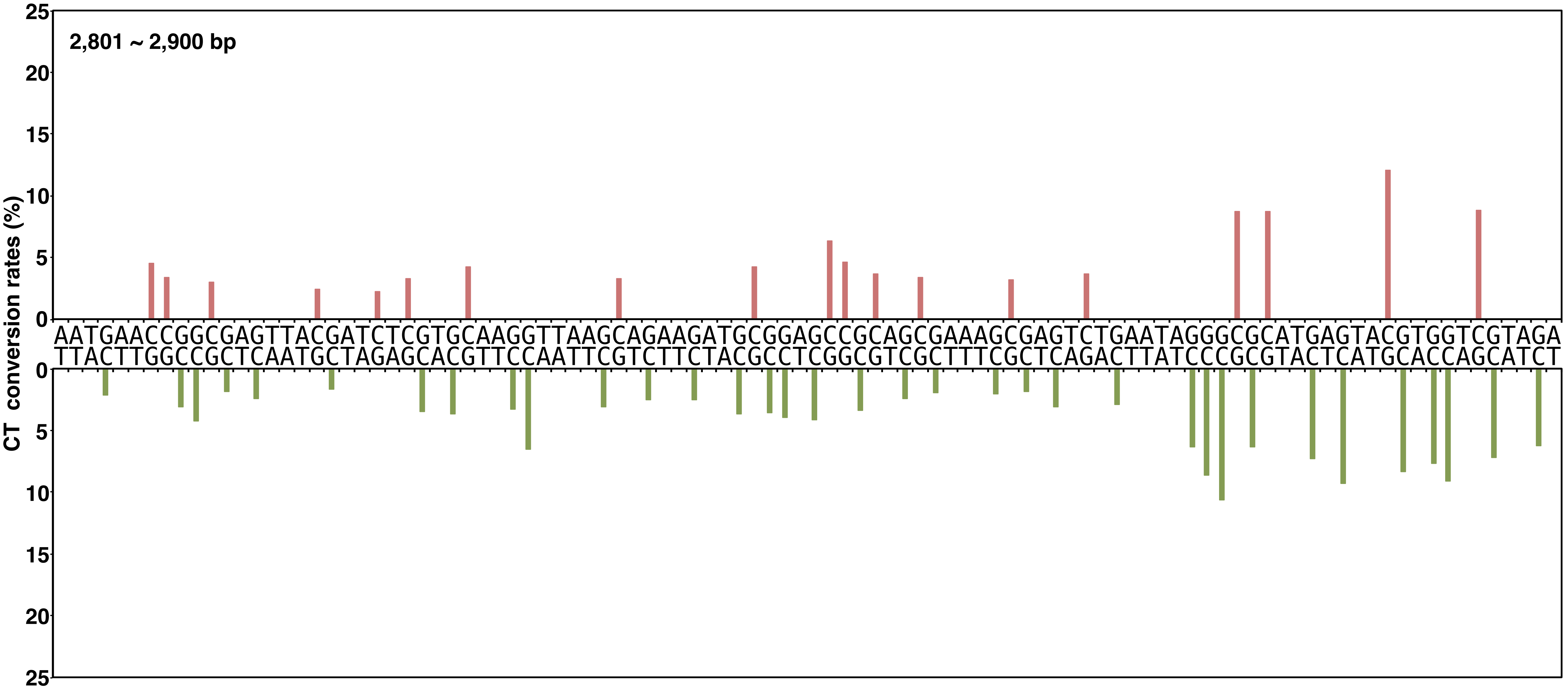


**Yano, Noguchi, and Niki Figure S2, continued**

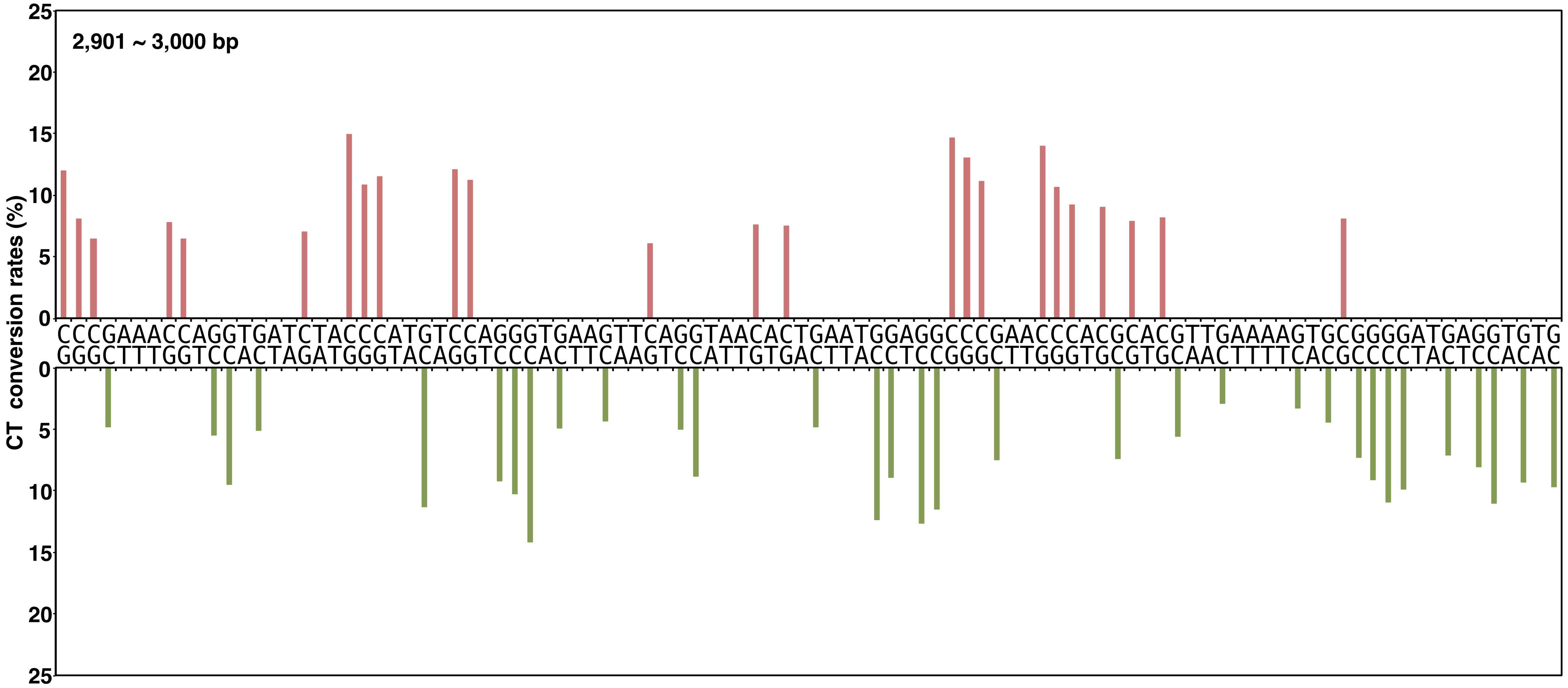


**Yano, Noguchi, and Niki Figure S2, continued**

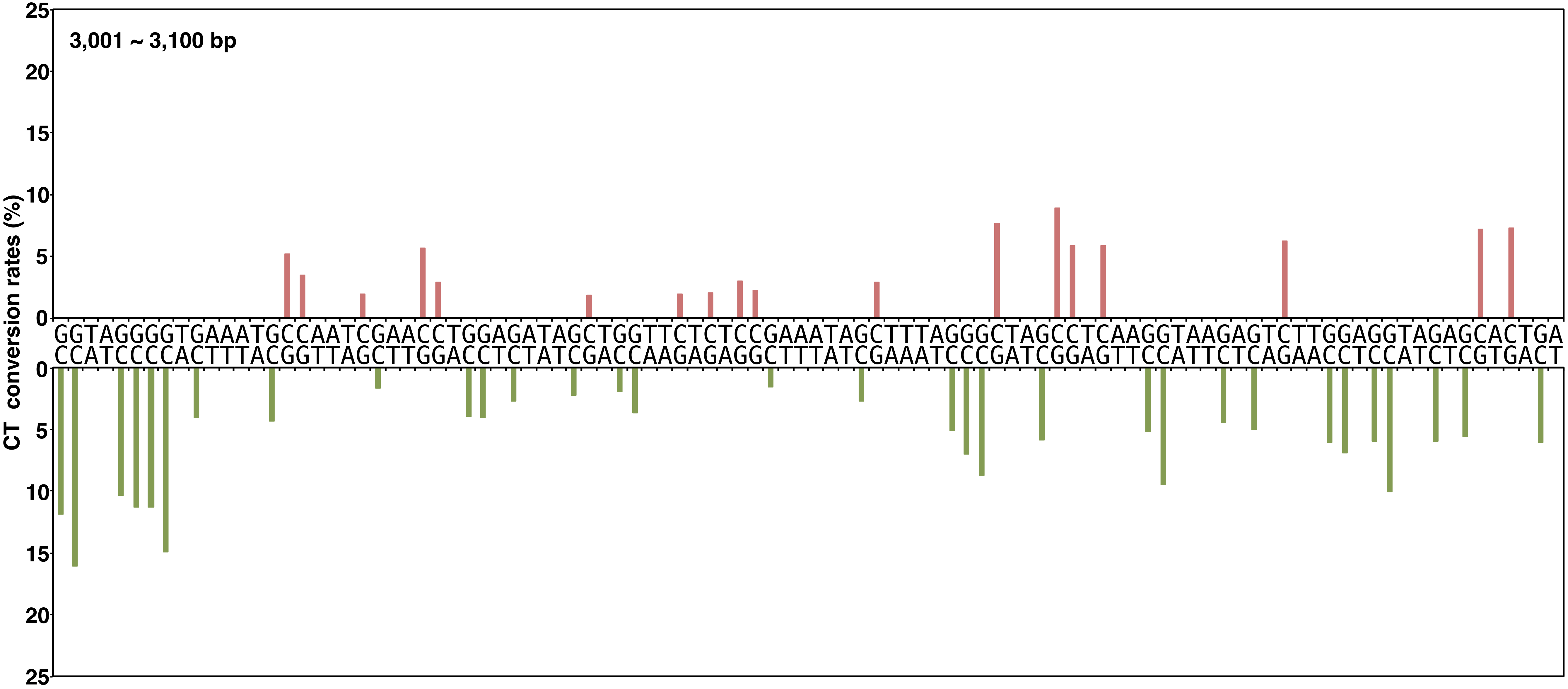


**Yano, Noguchi, and Niki Figure S2, continued**

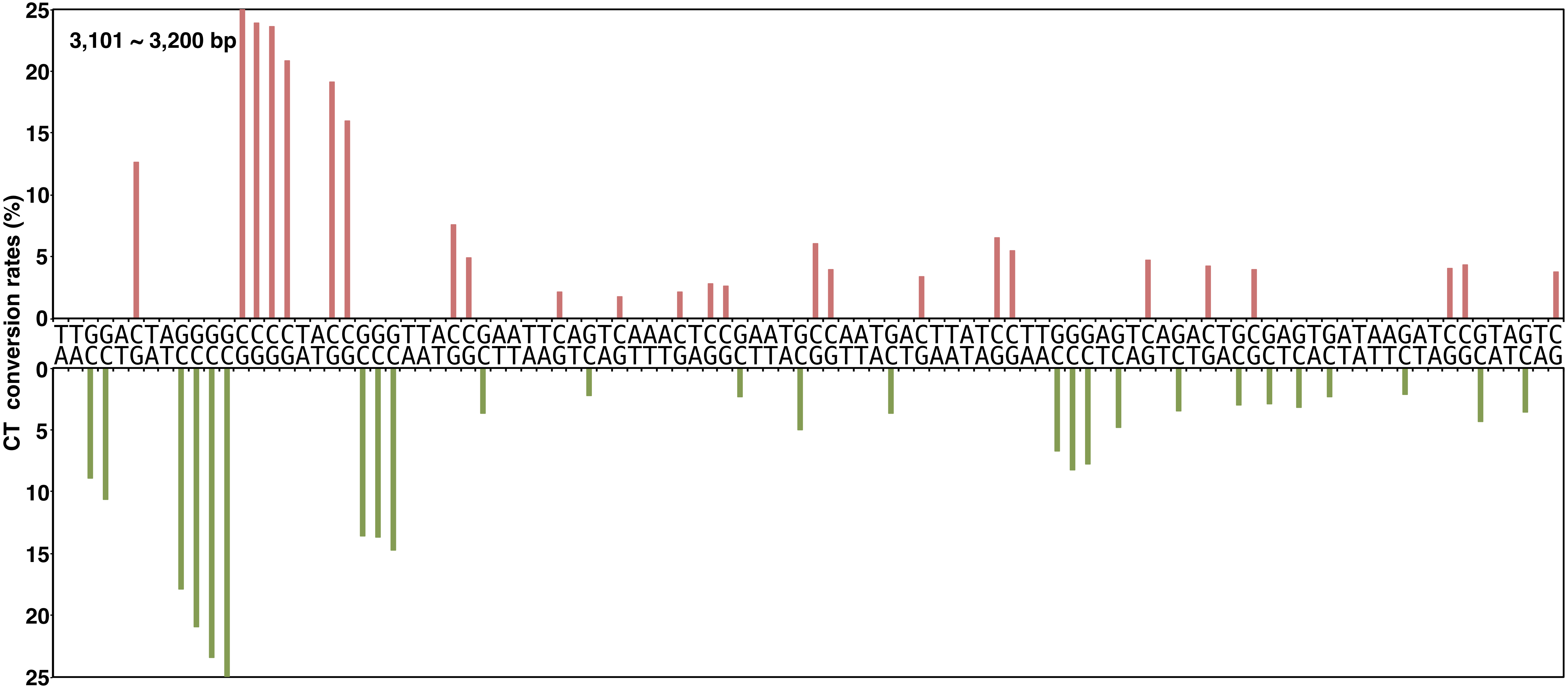


**Yano, Noguchi, and Niki Figure S2, continued**

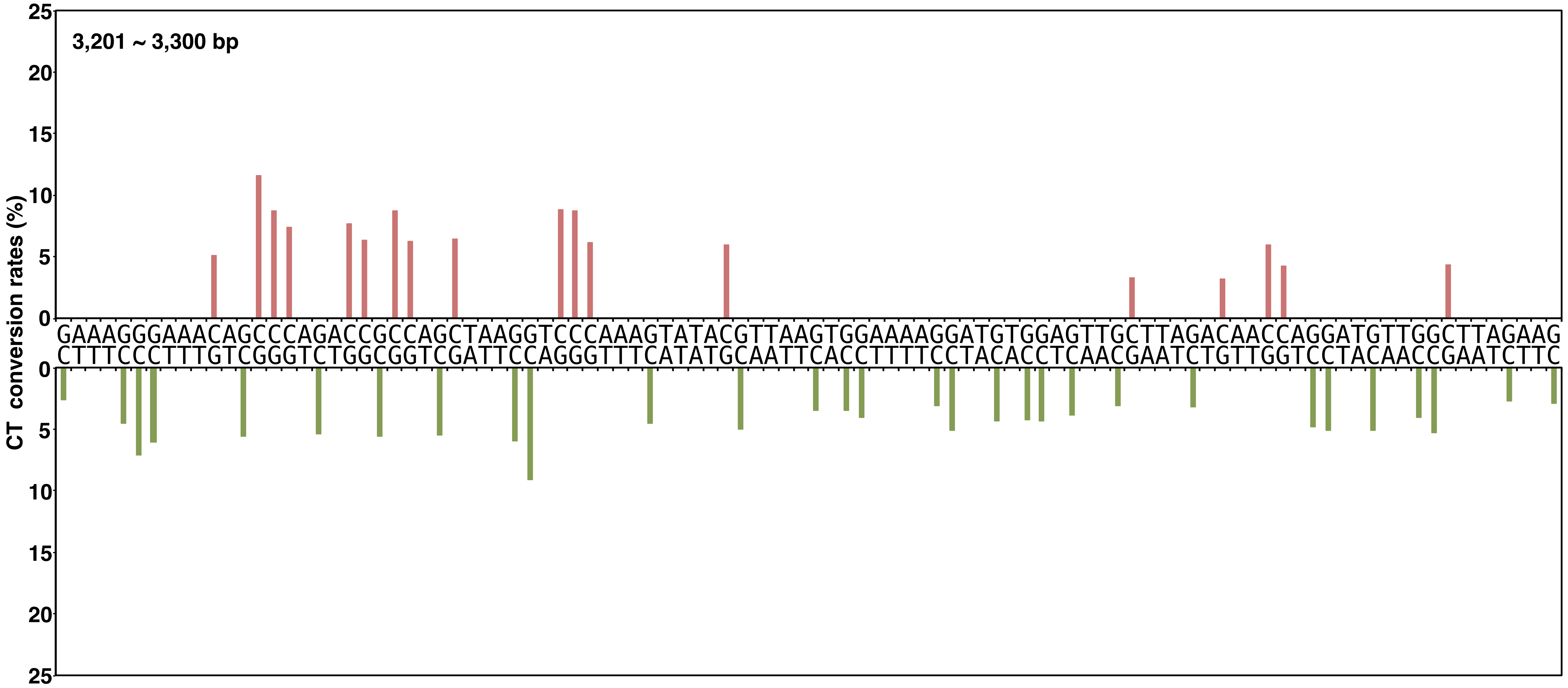


**Yano, Noguchi, and Niki Figure S2, continued**

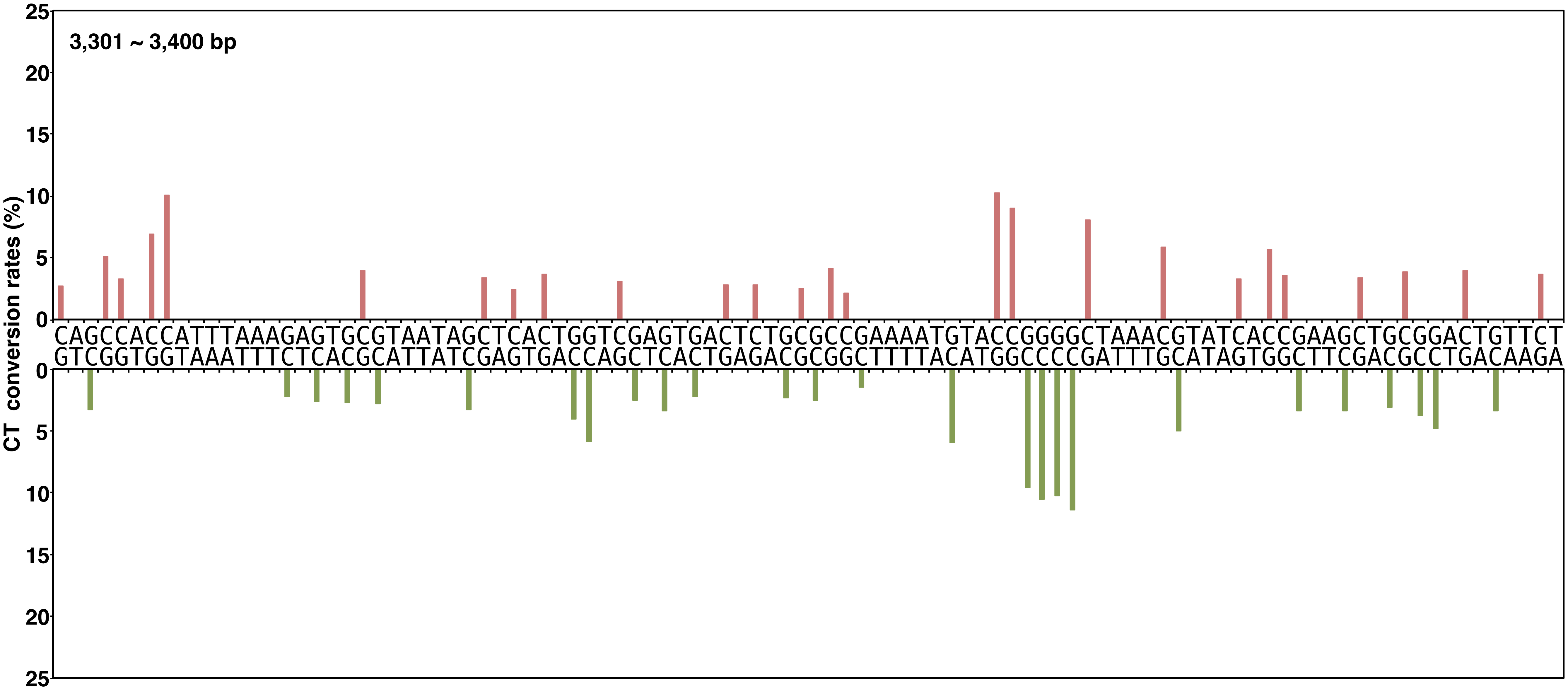


**Yano, Noguchi, and Niki Figure S2, continued**

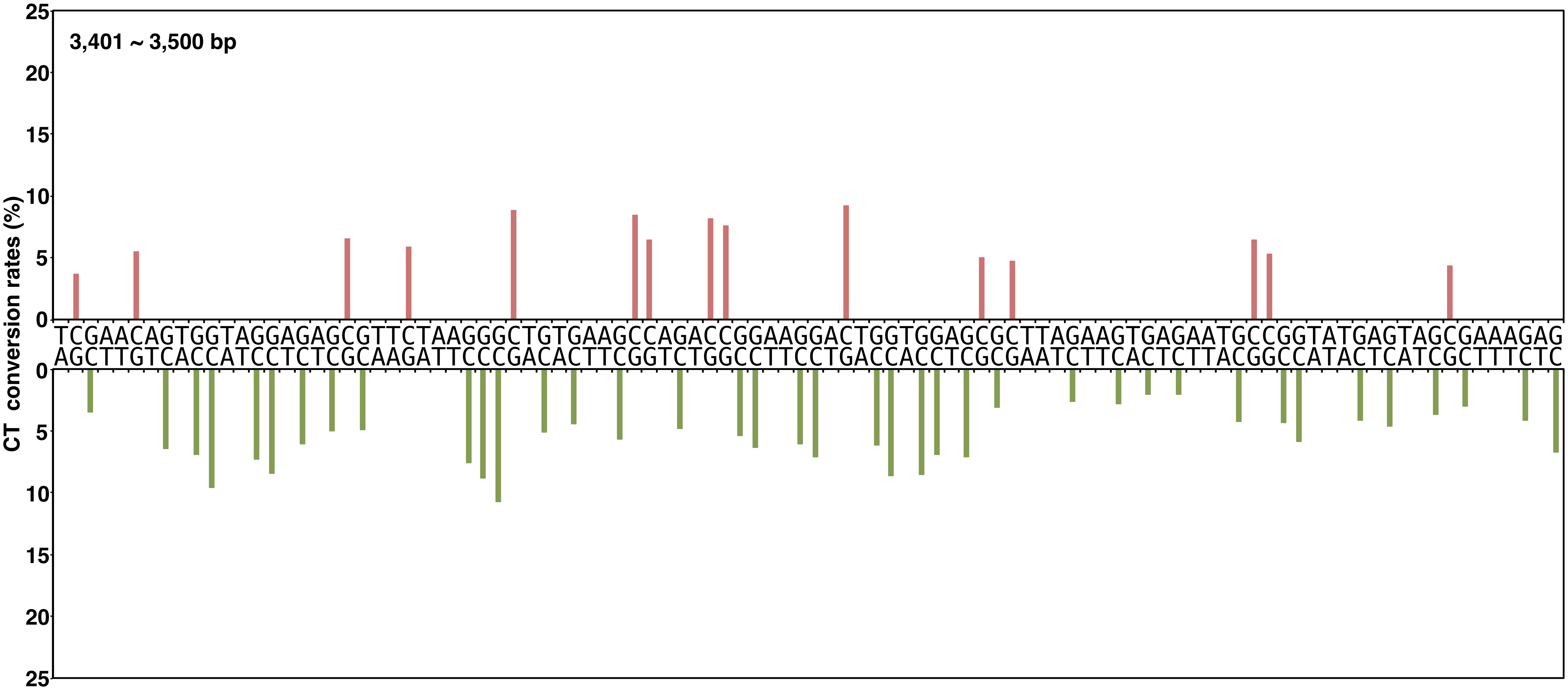


**Yano, Noguchi, and Niki Figure S2, continued**

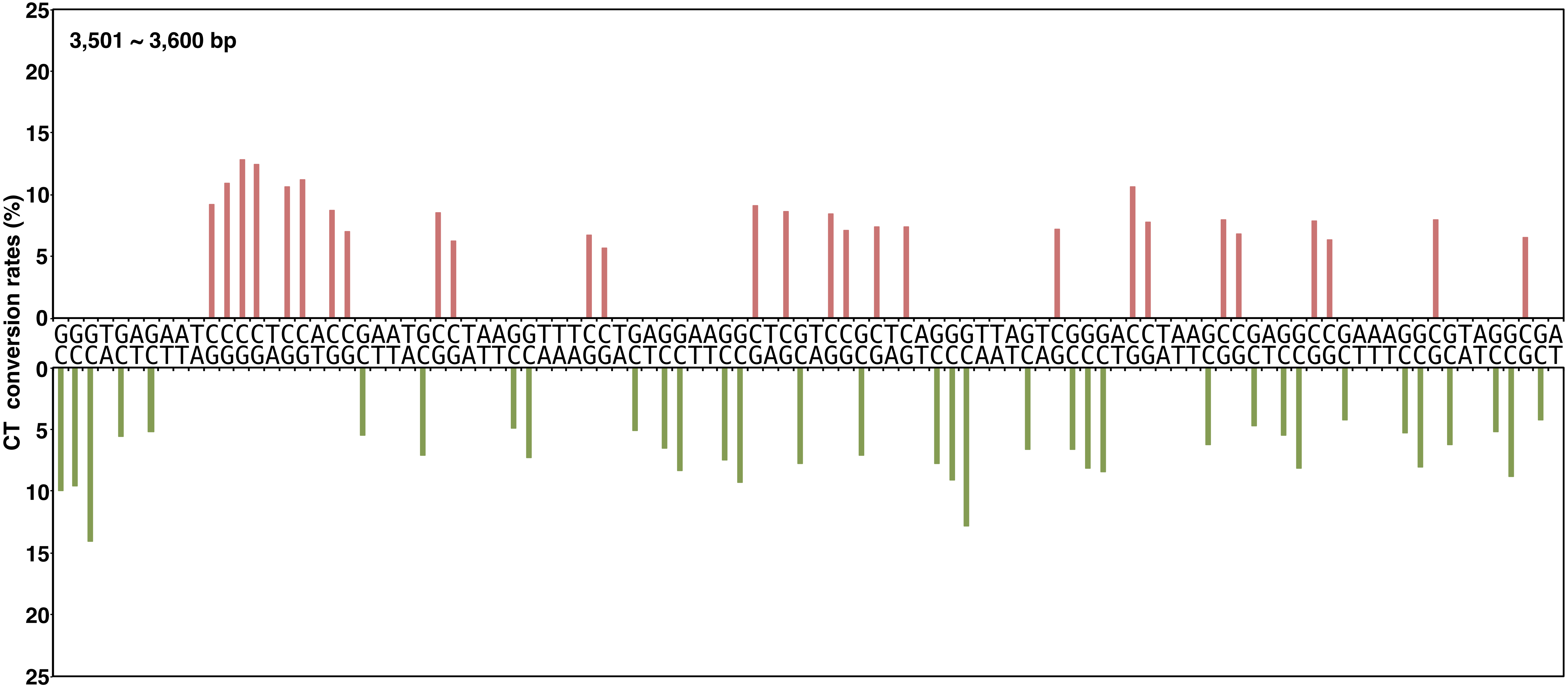


**Yano, Noguchi, and Niki Figure S2, continued**

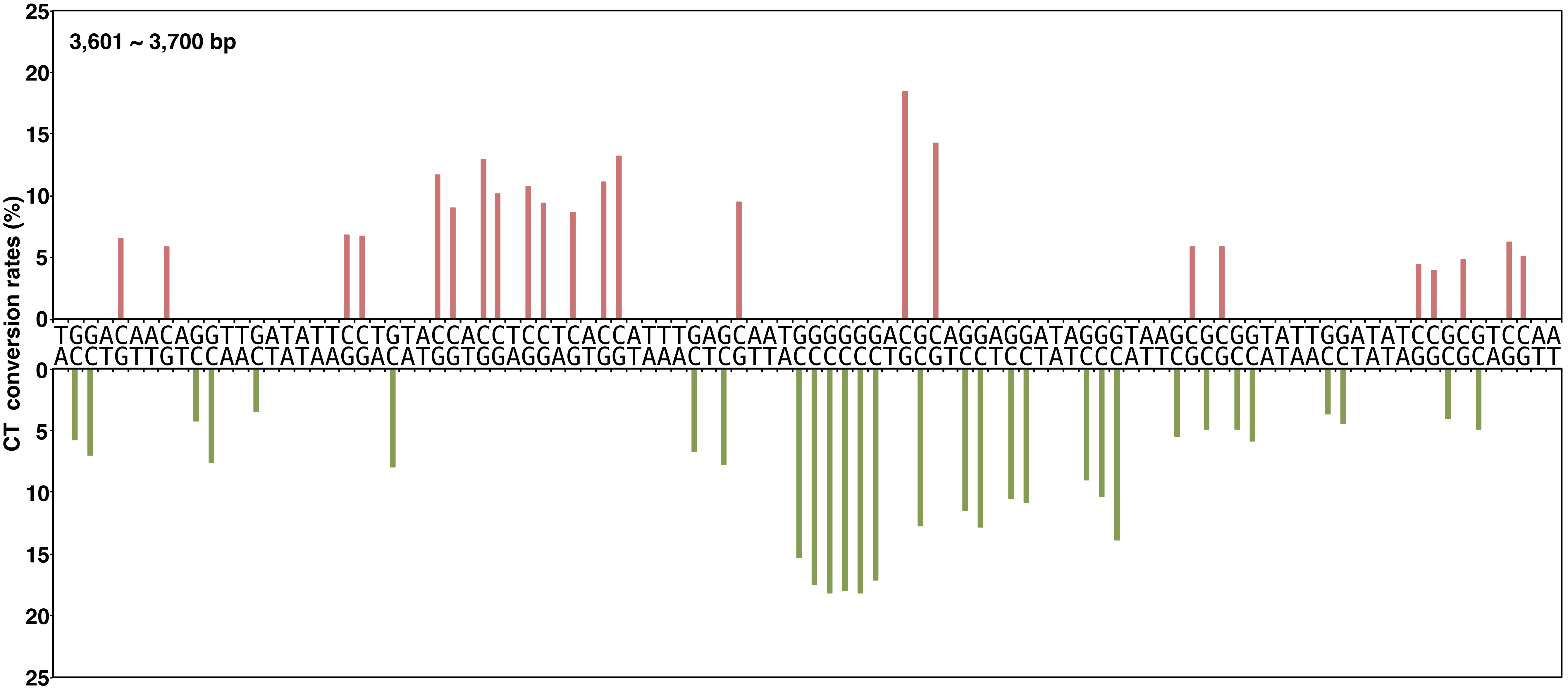


**Yano, Noguchi, and Niki Figure S2, continued**

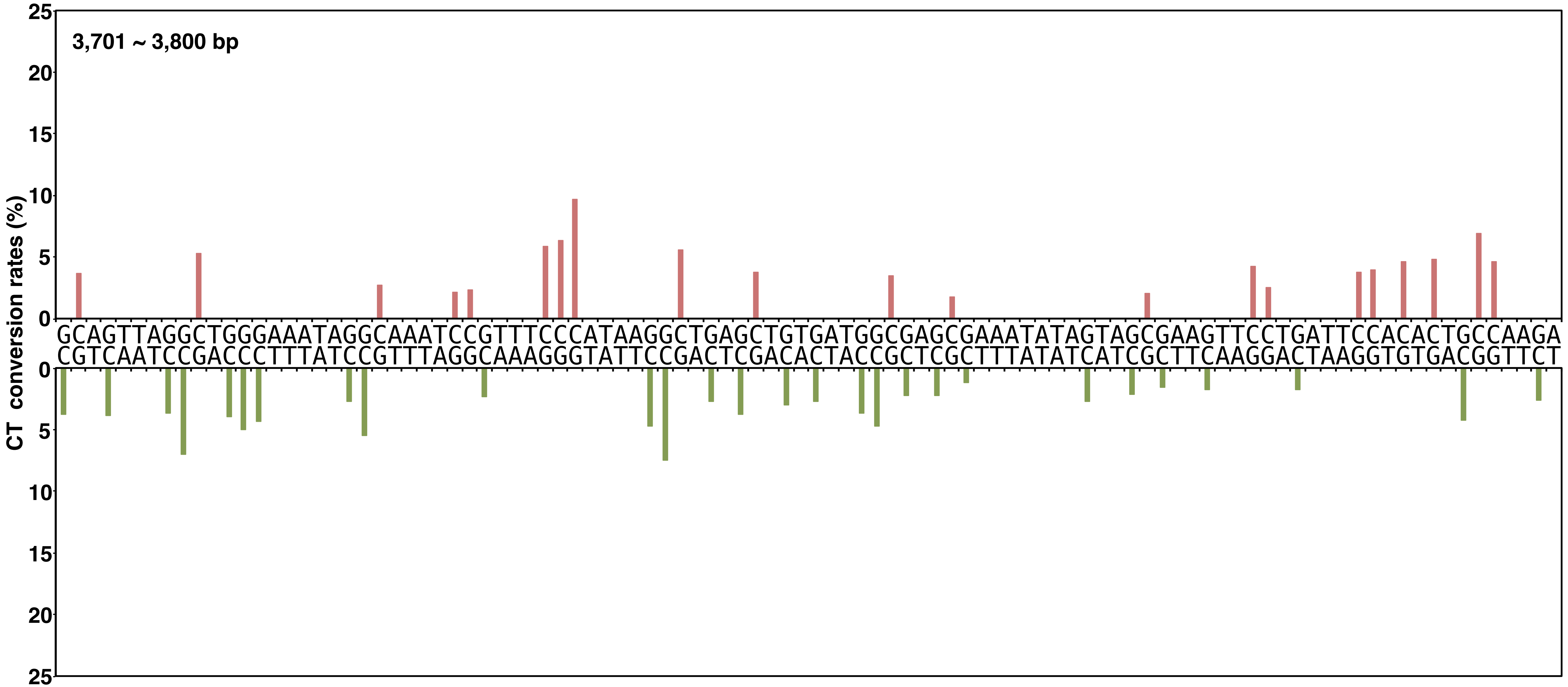


**Yano, Noguchi, and Niki Figure S2, continued**

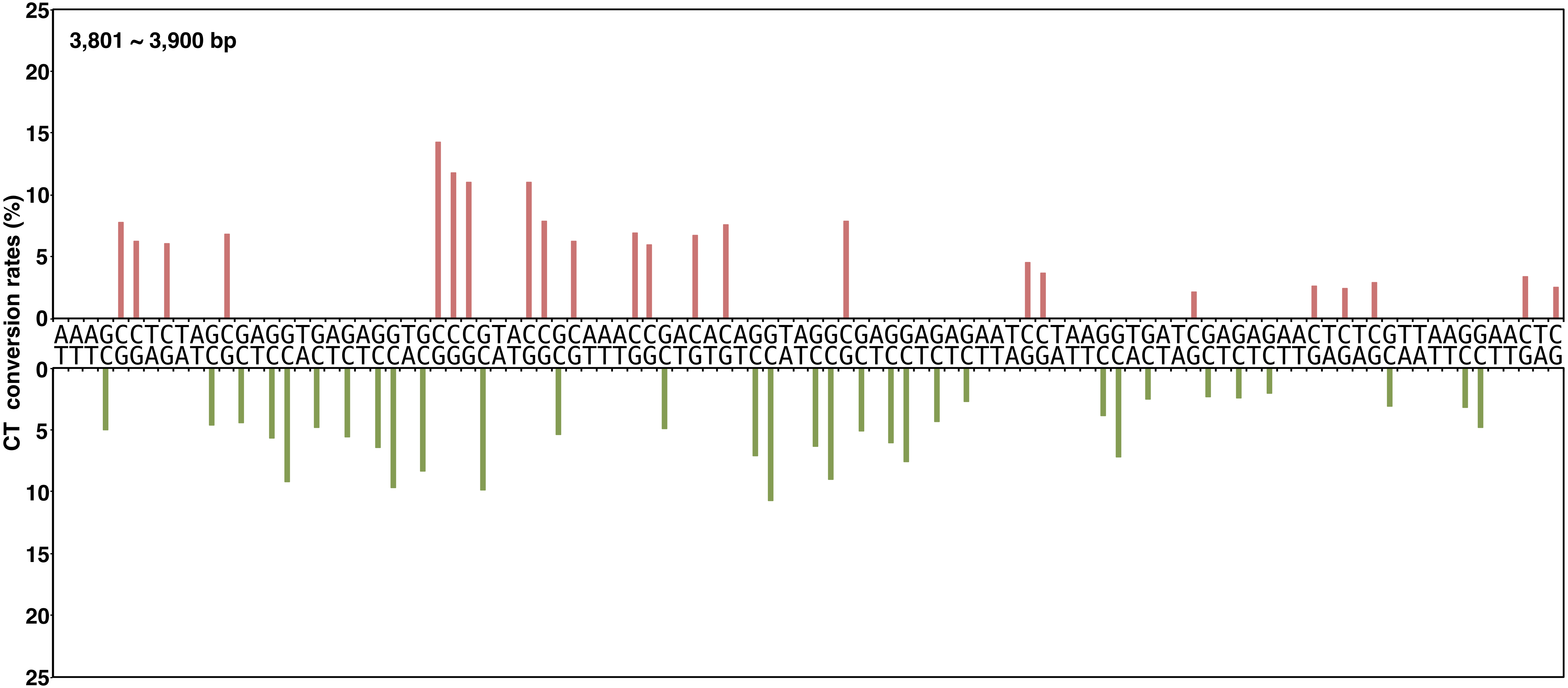


**Yano, Noguchi, and Niki Figure S2, continued**

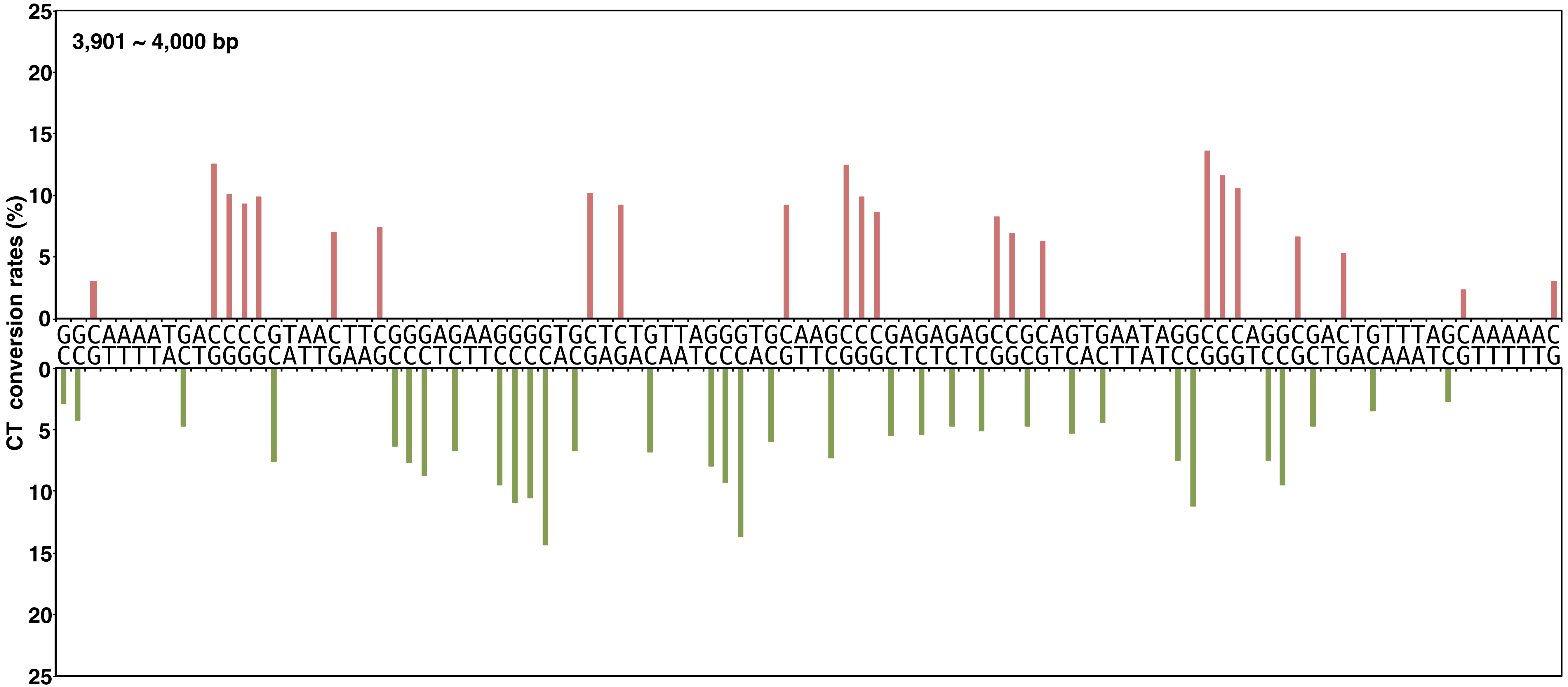


**Yano, Noguchi, and Niki Figure S2, continued**

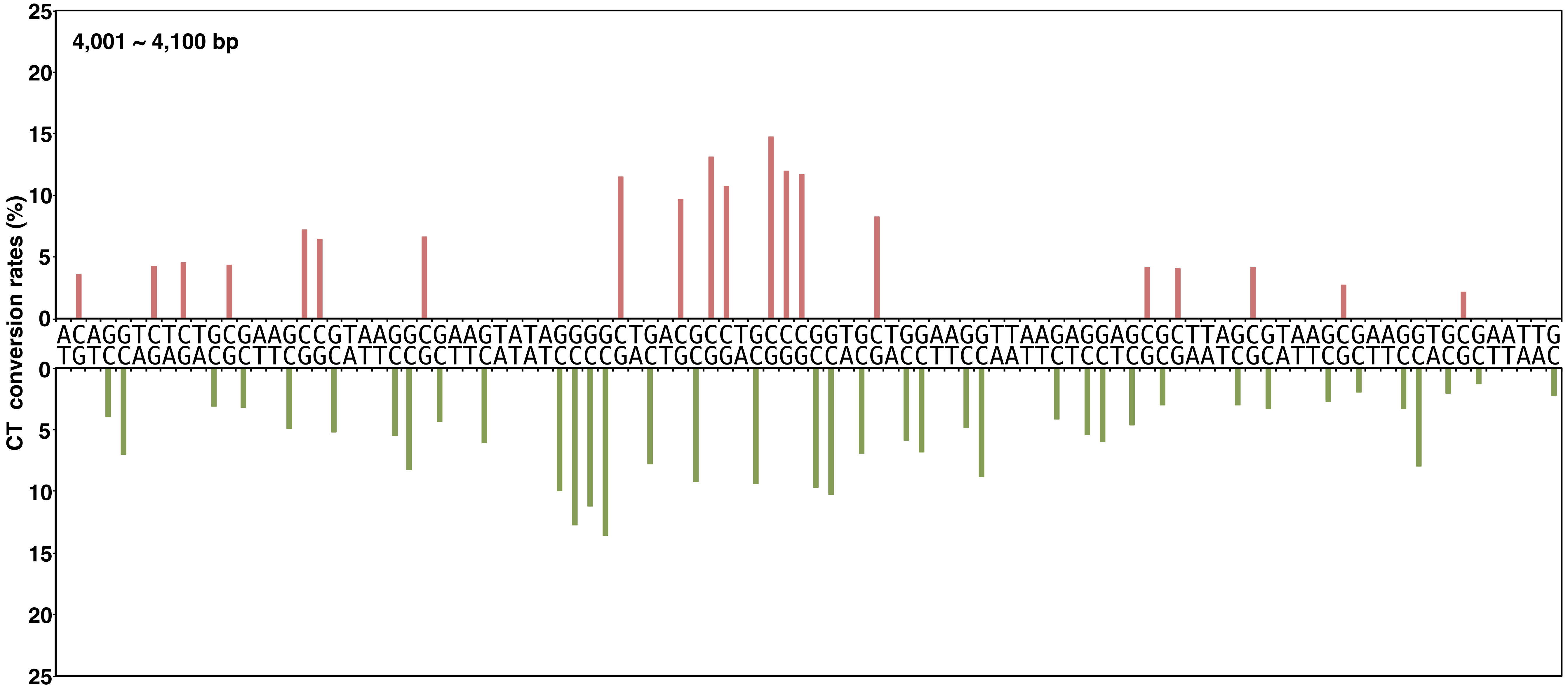


**Yano, Noguchi, and Niki Figure S2, continued**

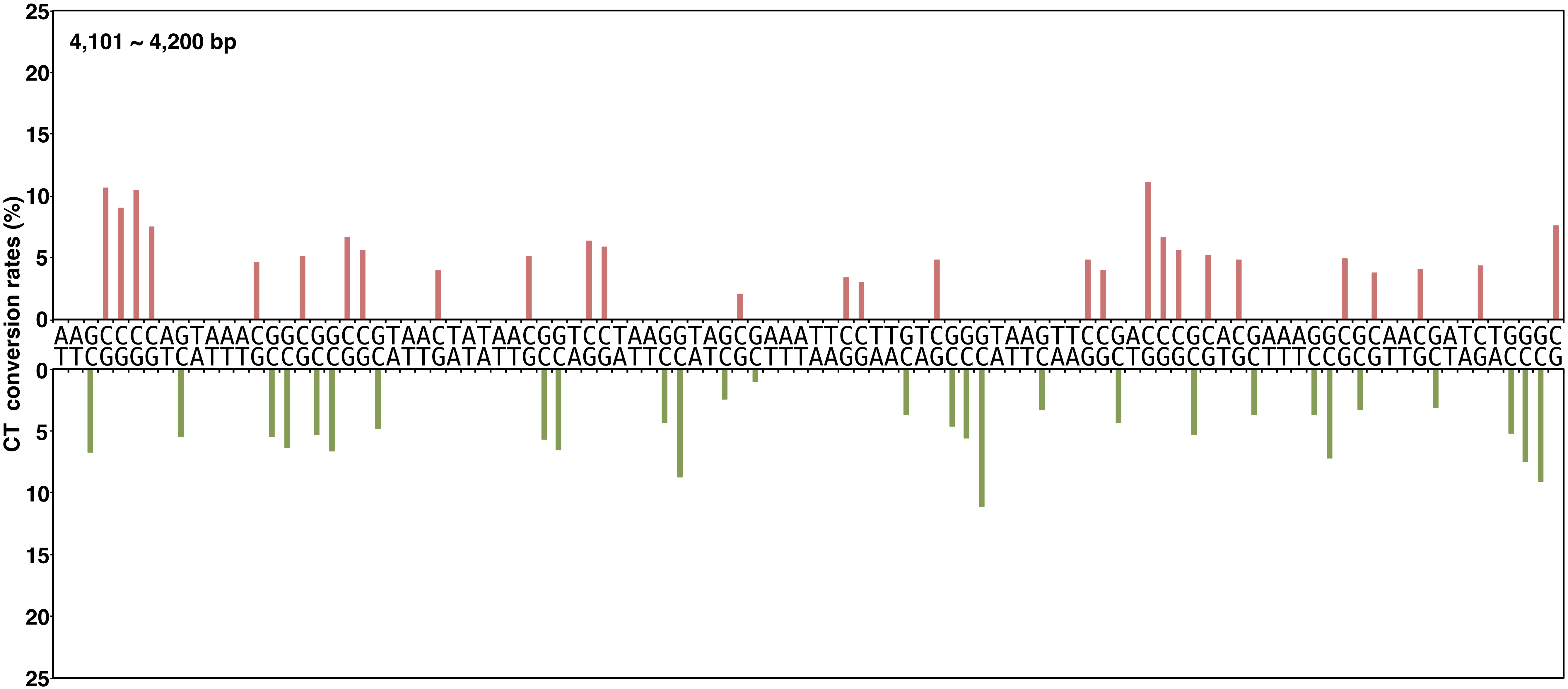


**Yano, Noguchi, and Niki Figure S2, continued**

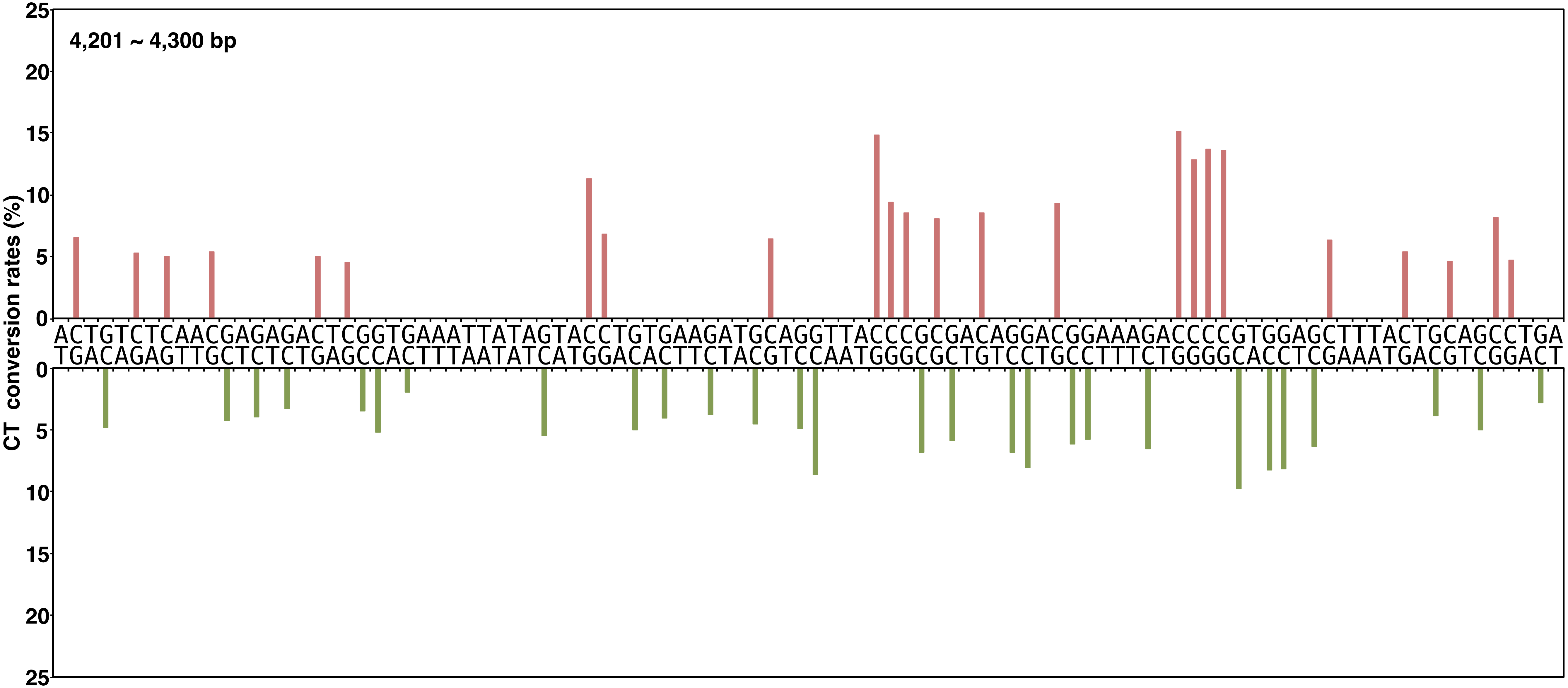


**Yano, Noguchi, and Niki Figure S2, continued**

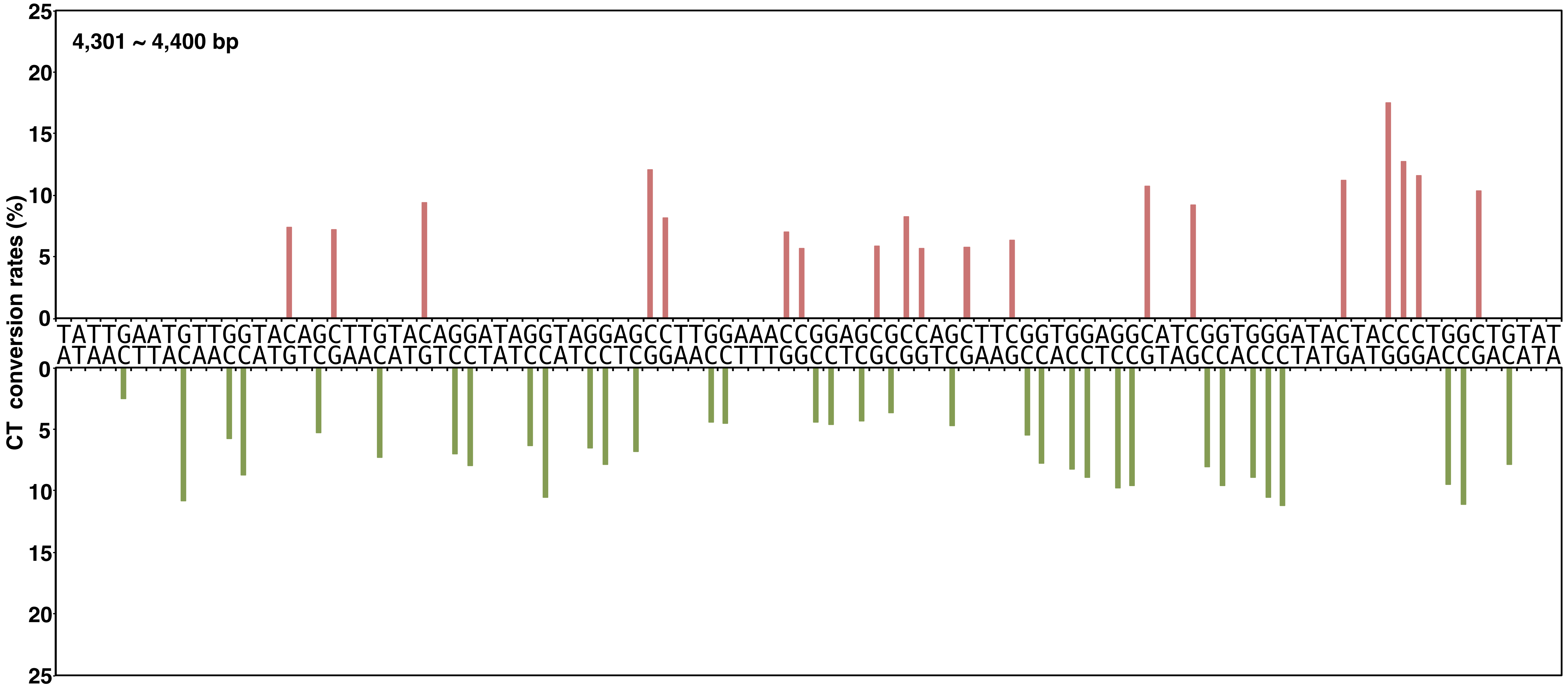


Yano, Noguchi, and Niki Figure S2, continued

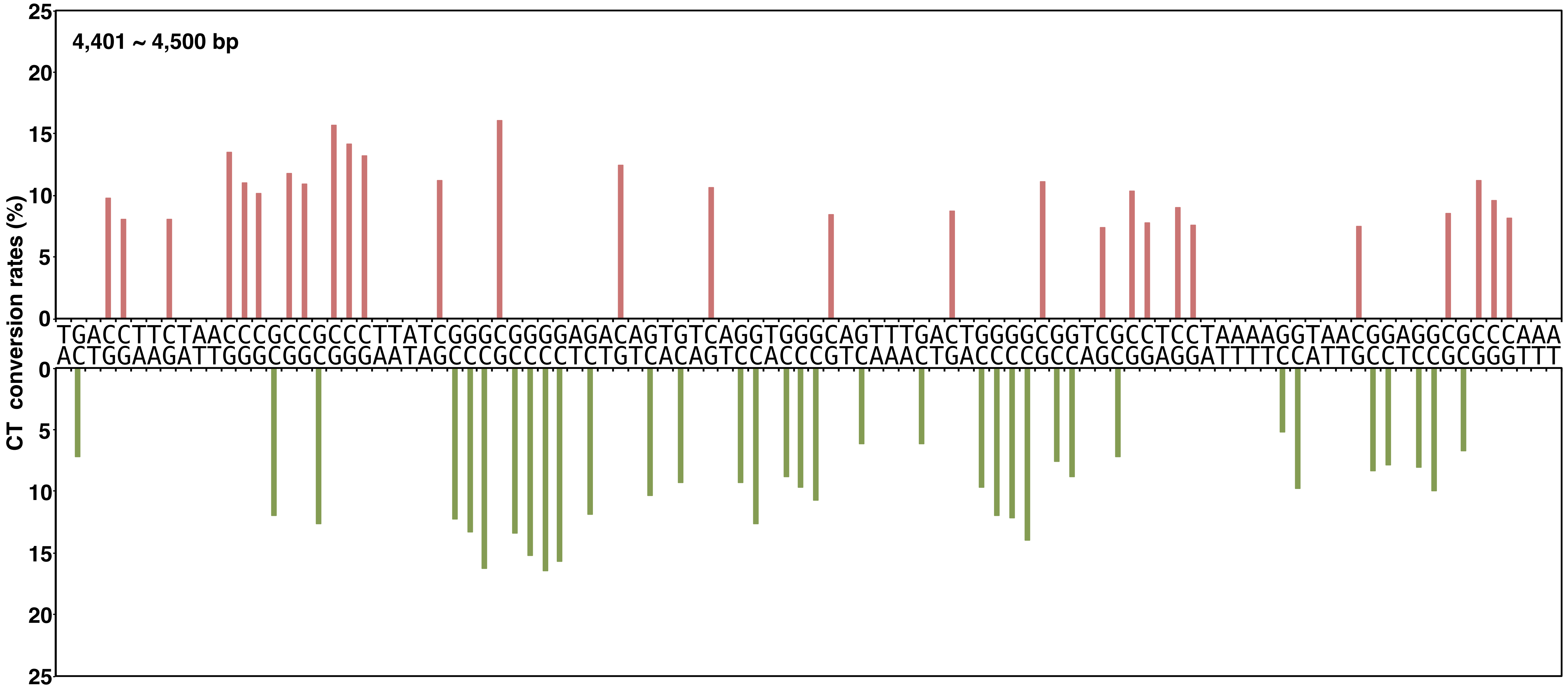


**Yano, Noguchi, and Niki Figure S2, continued**

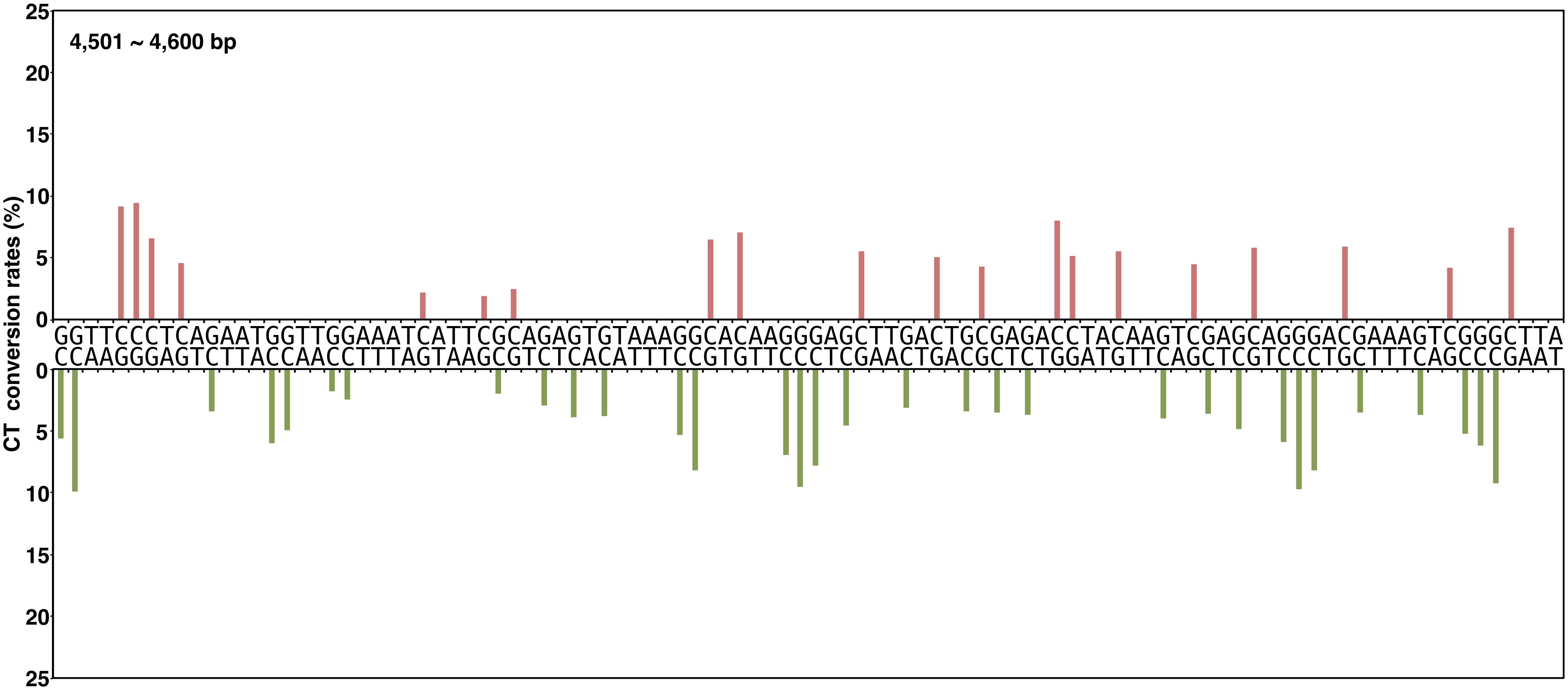


**Yano, Noguchi, and Niki Figure S2, continued**

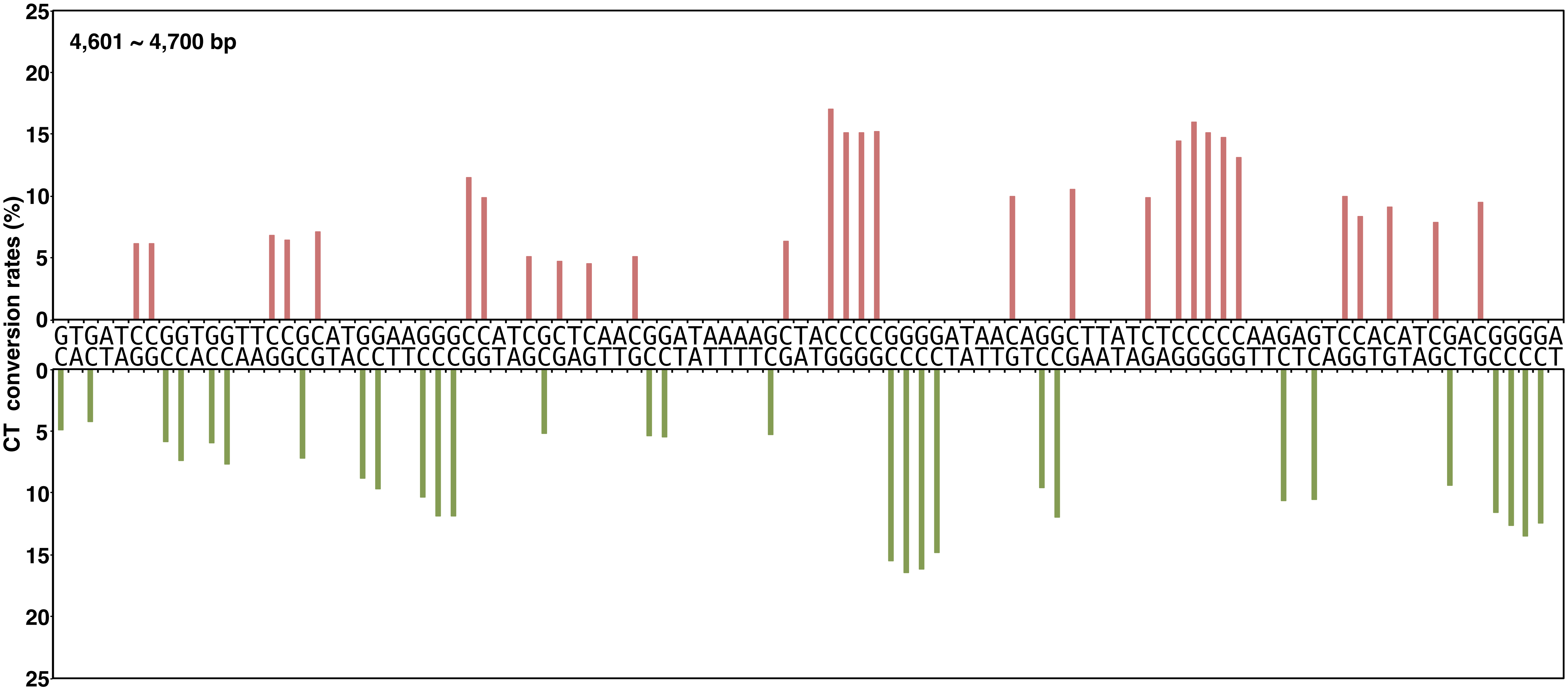


**Yano, Noguchi, and Niki Figure S2, continued**

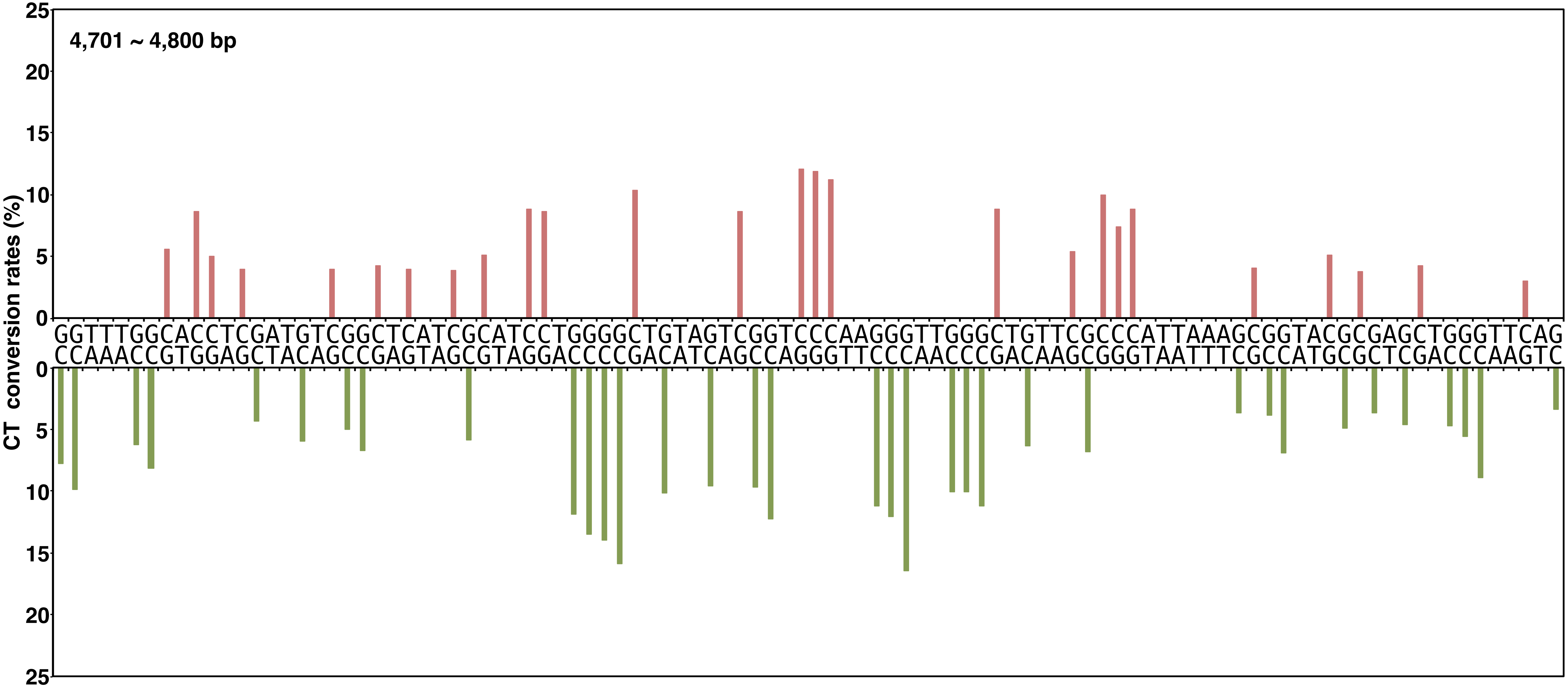


Yano, Noguchi, and Niki Figure S2, continued

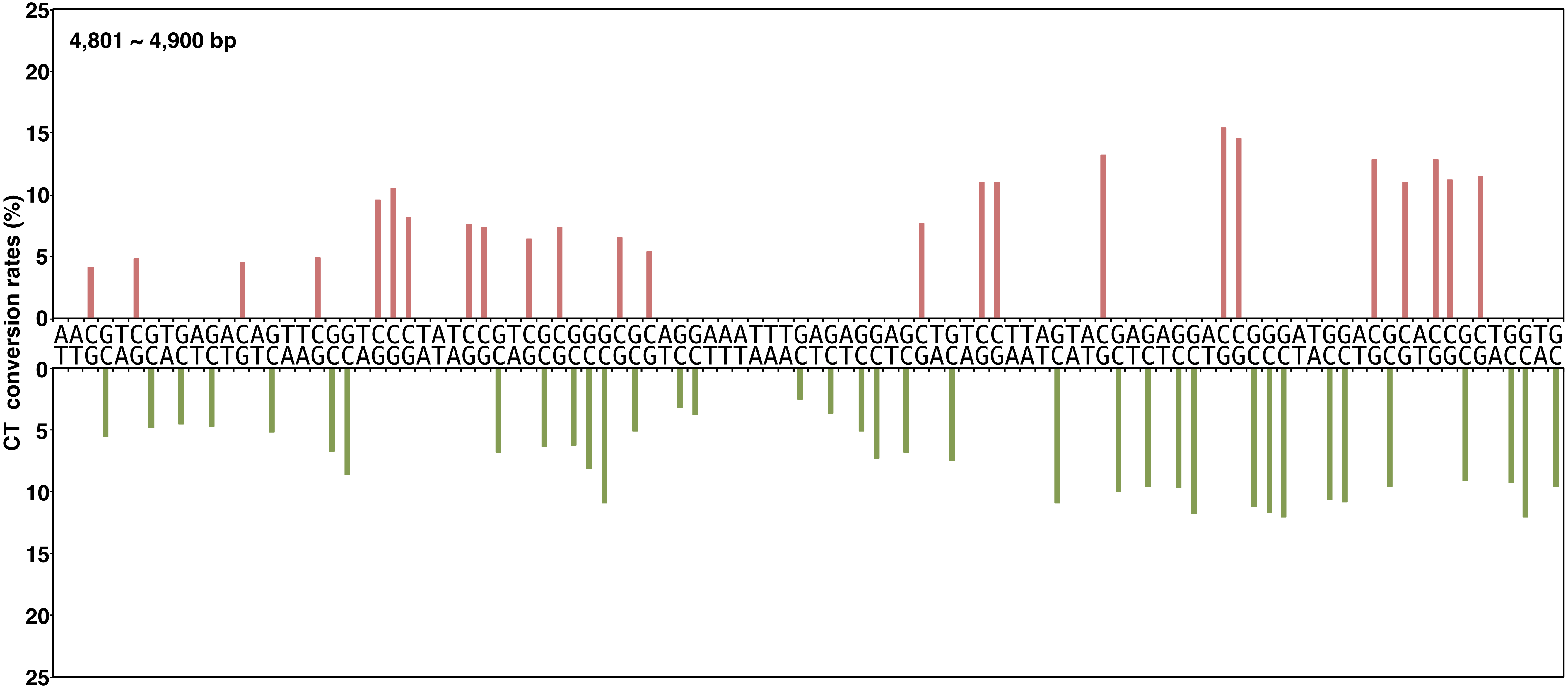


**Yano, Noguchi, and Niki Figure S2, continued**

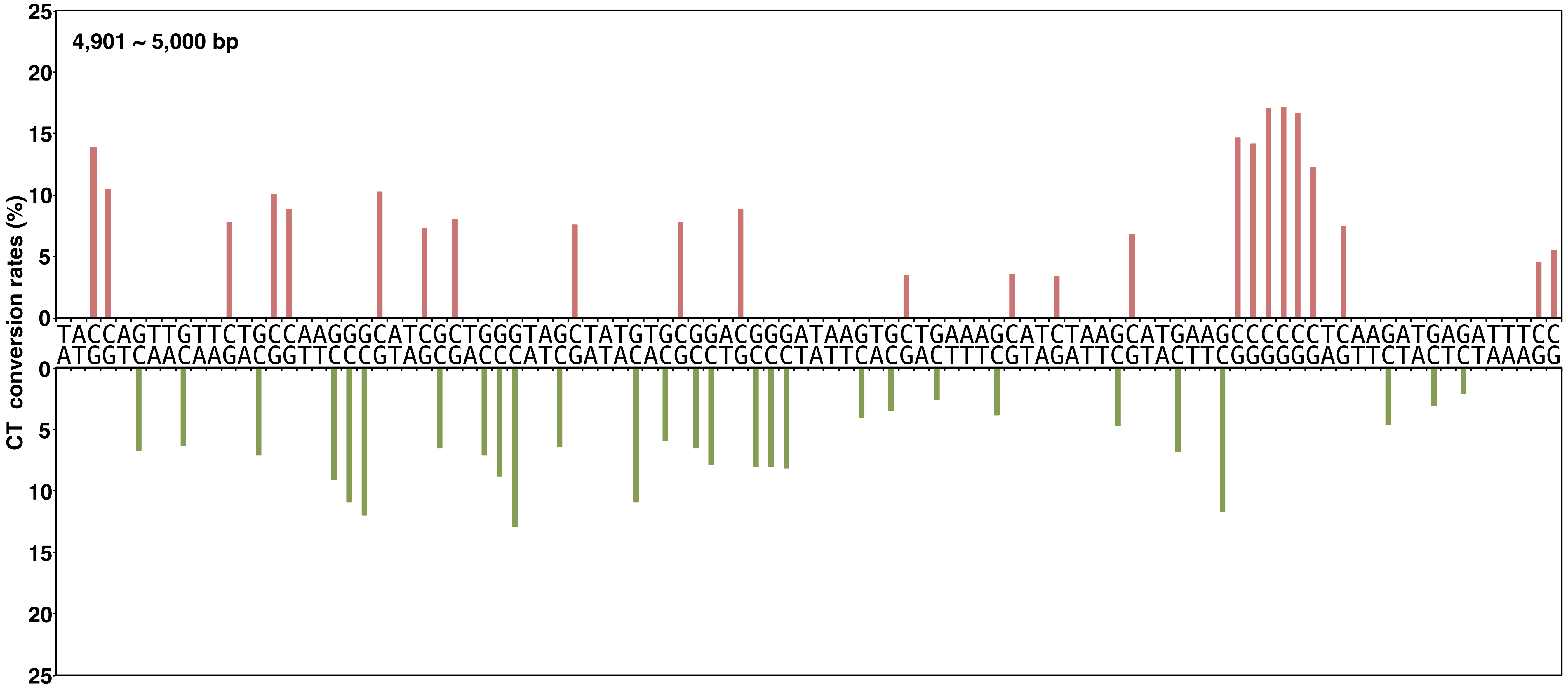


Yano, Noguchi, and Niki Figure S2, continued

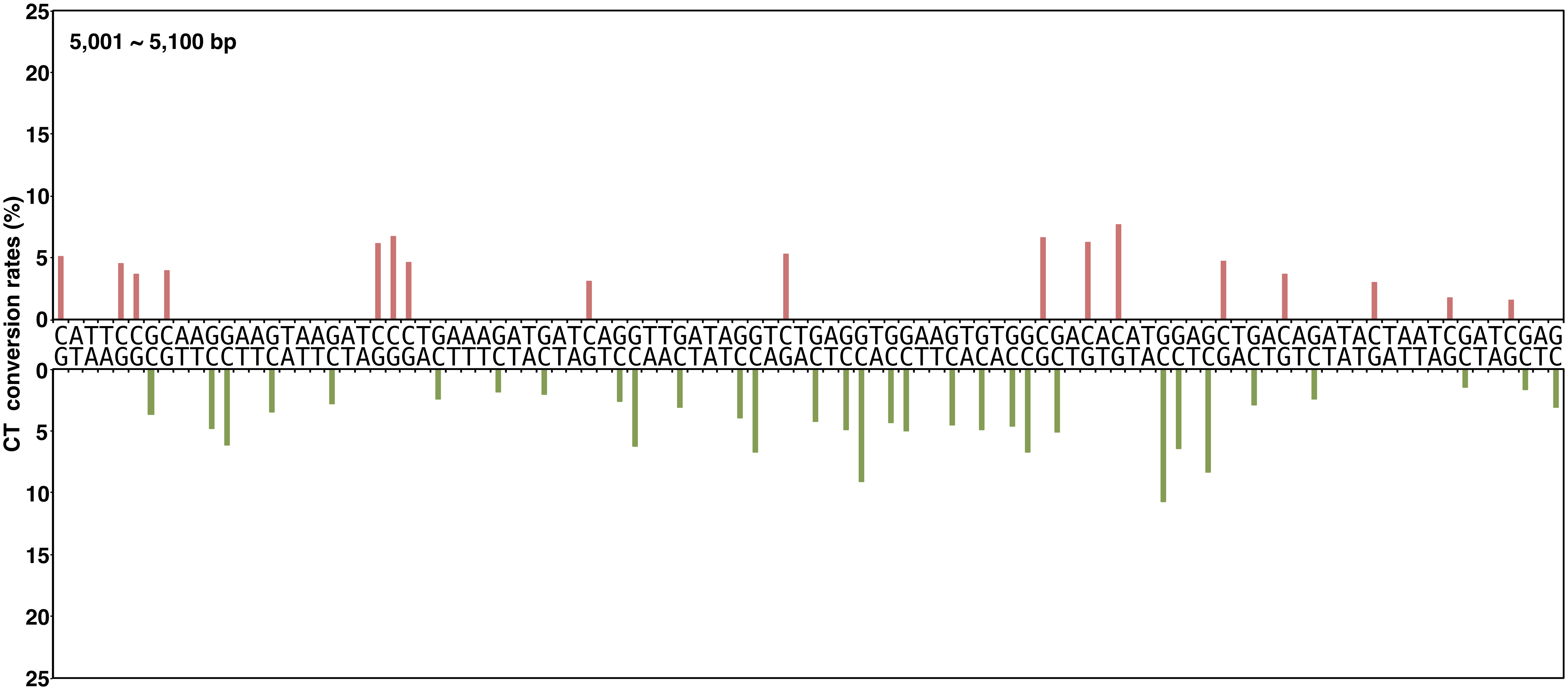


**Yano, Noguchi, and Niki Figure S2, continued**

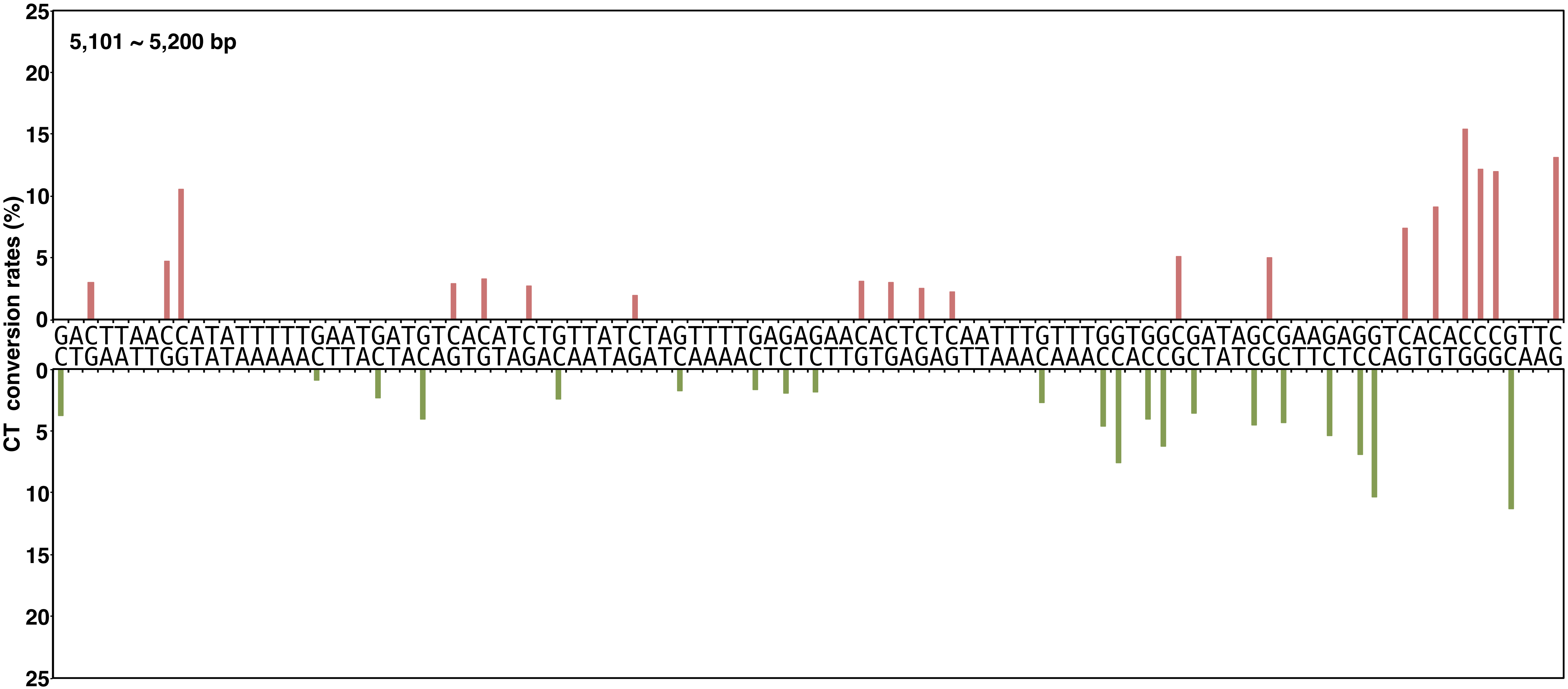


**Yano, Noguchi, and Niki Figure S2, continued**

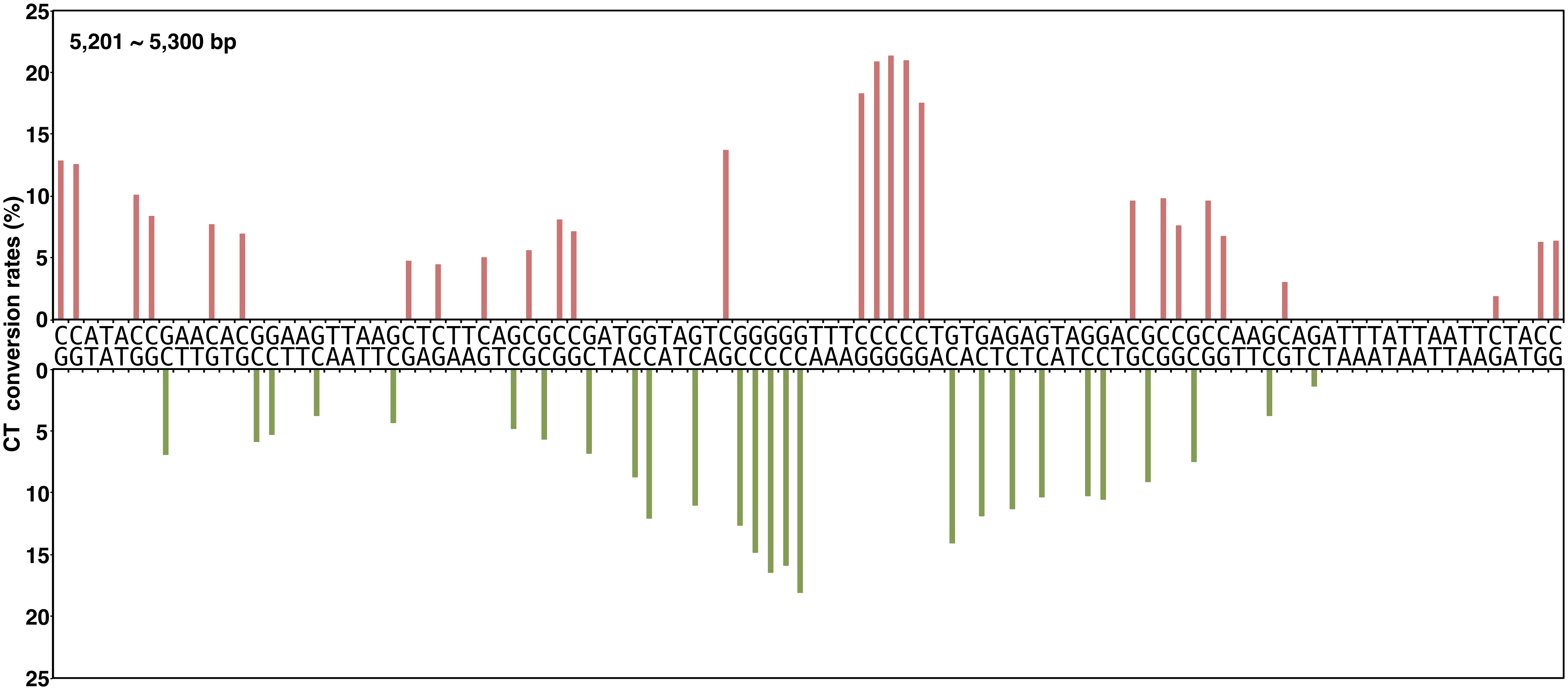


**Yano, Noguchi, and Niki Figure S2, continued**

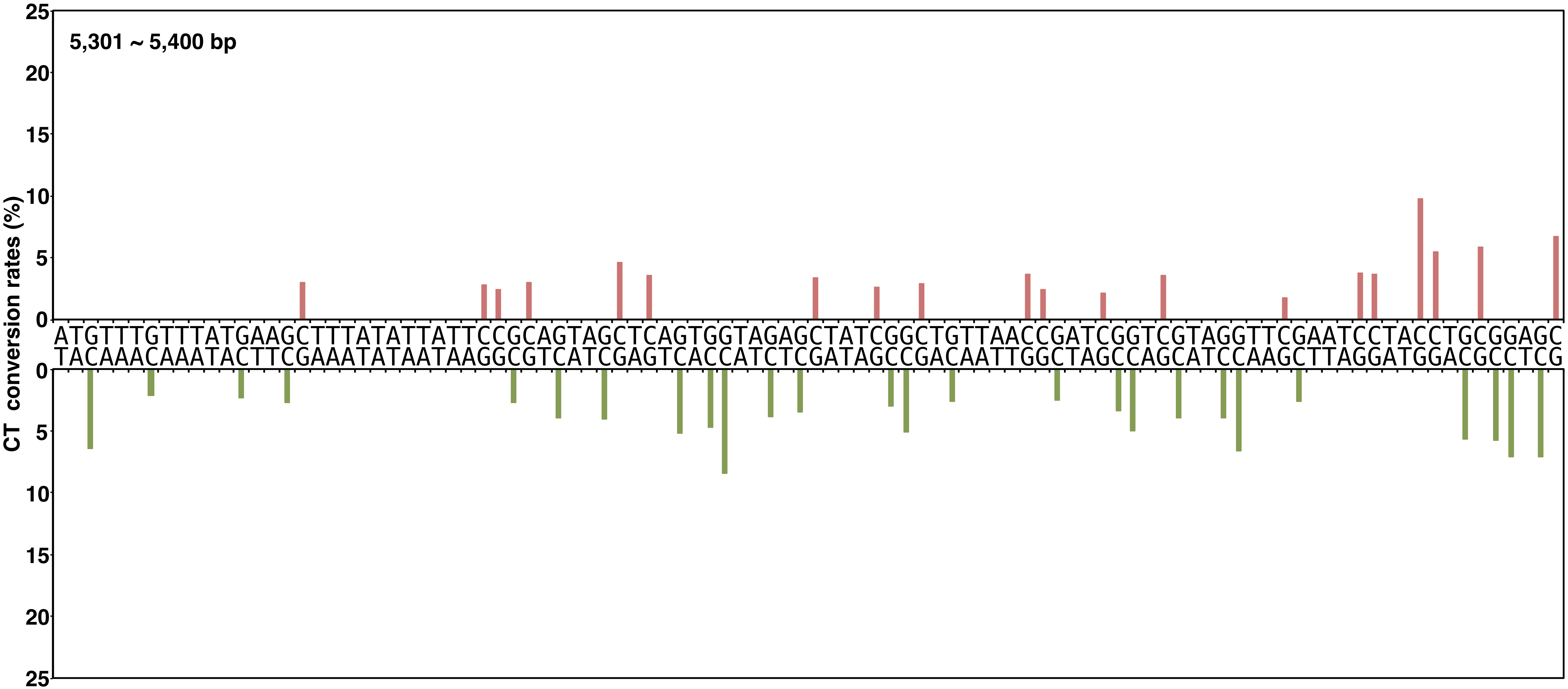


Yano, Noguchi, and Niki Figure S2, continued

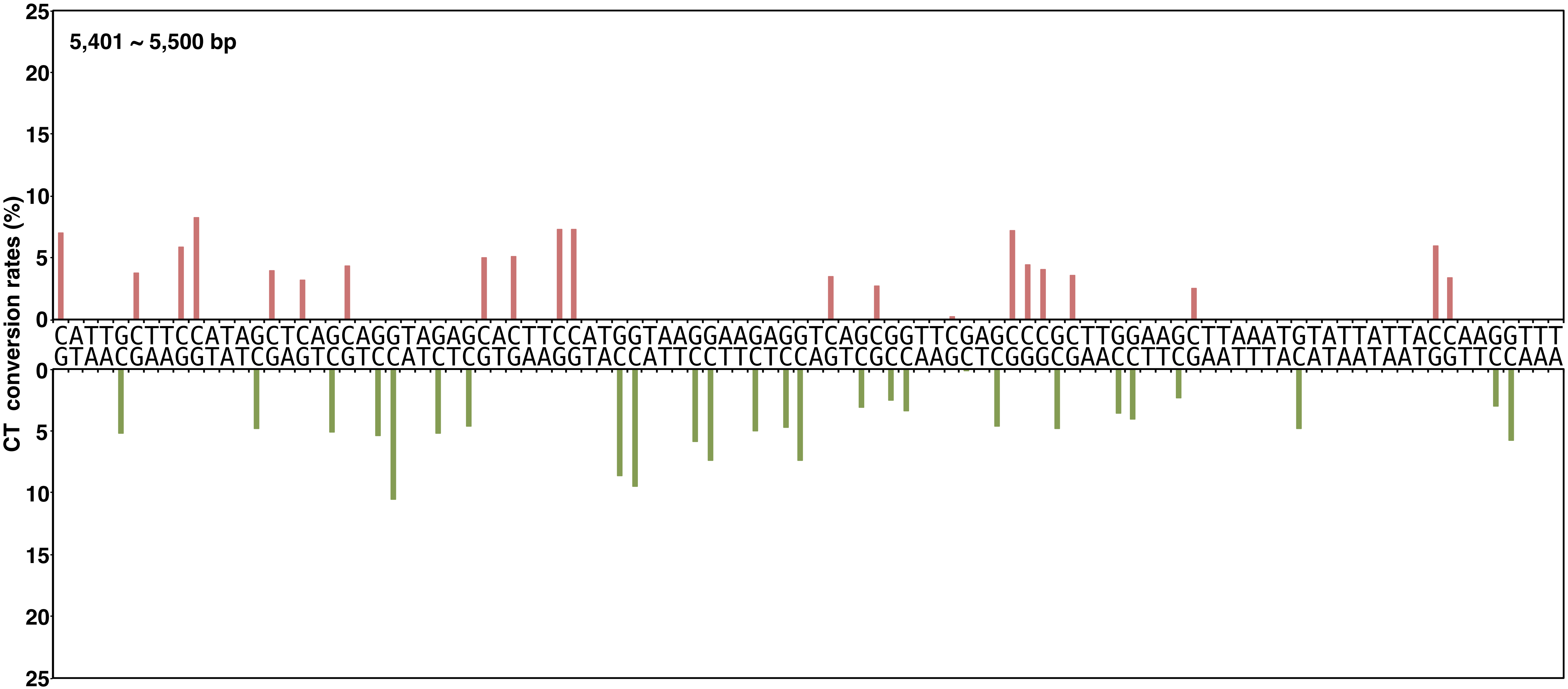


**Yano, Noguchi, and Niki Figure S2, continued**

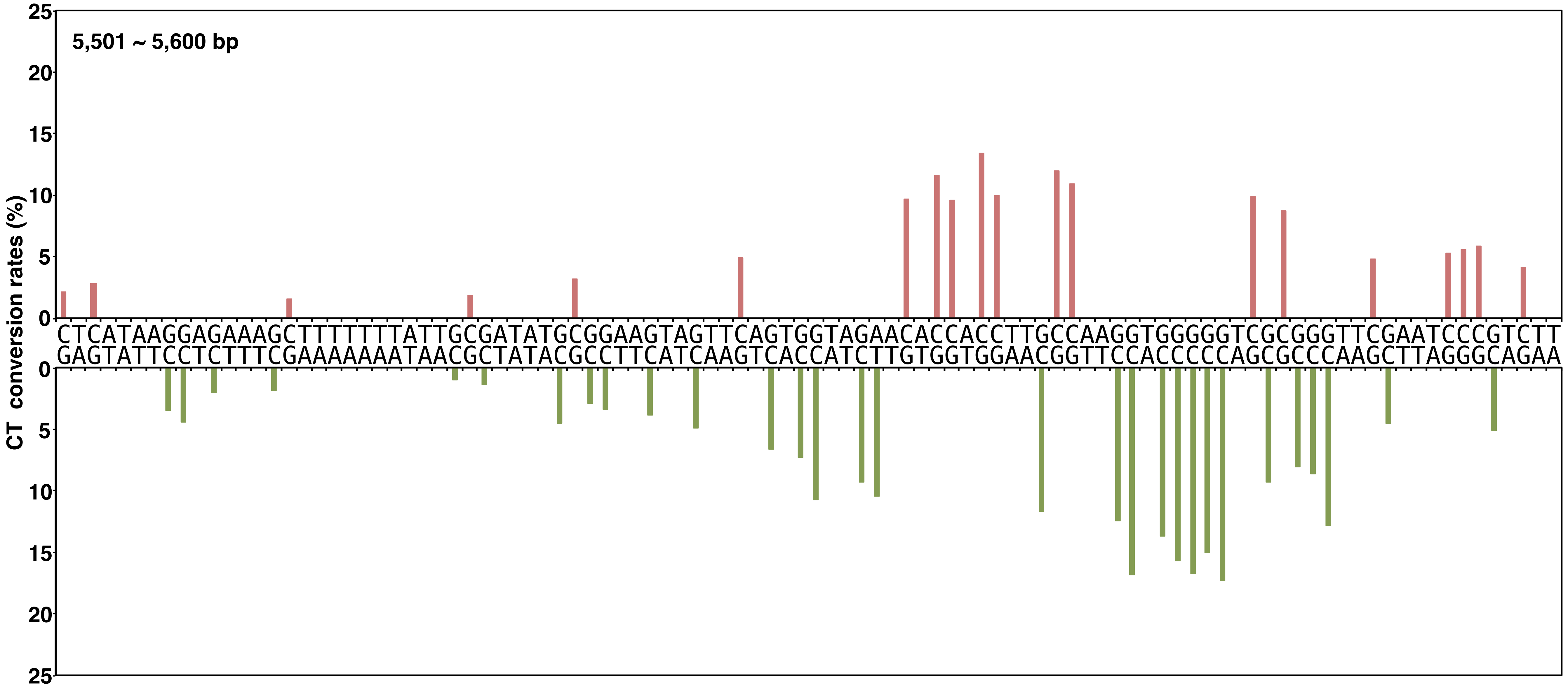


**Yano, Noguchi, and Niki Figure S2, continued**

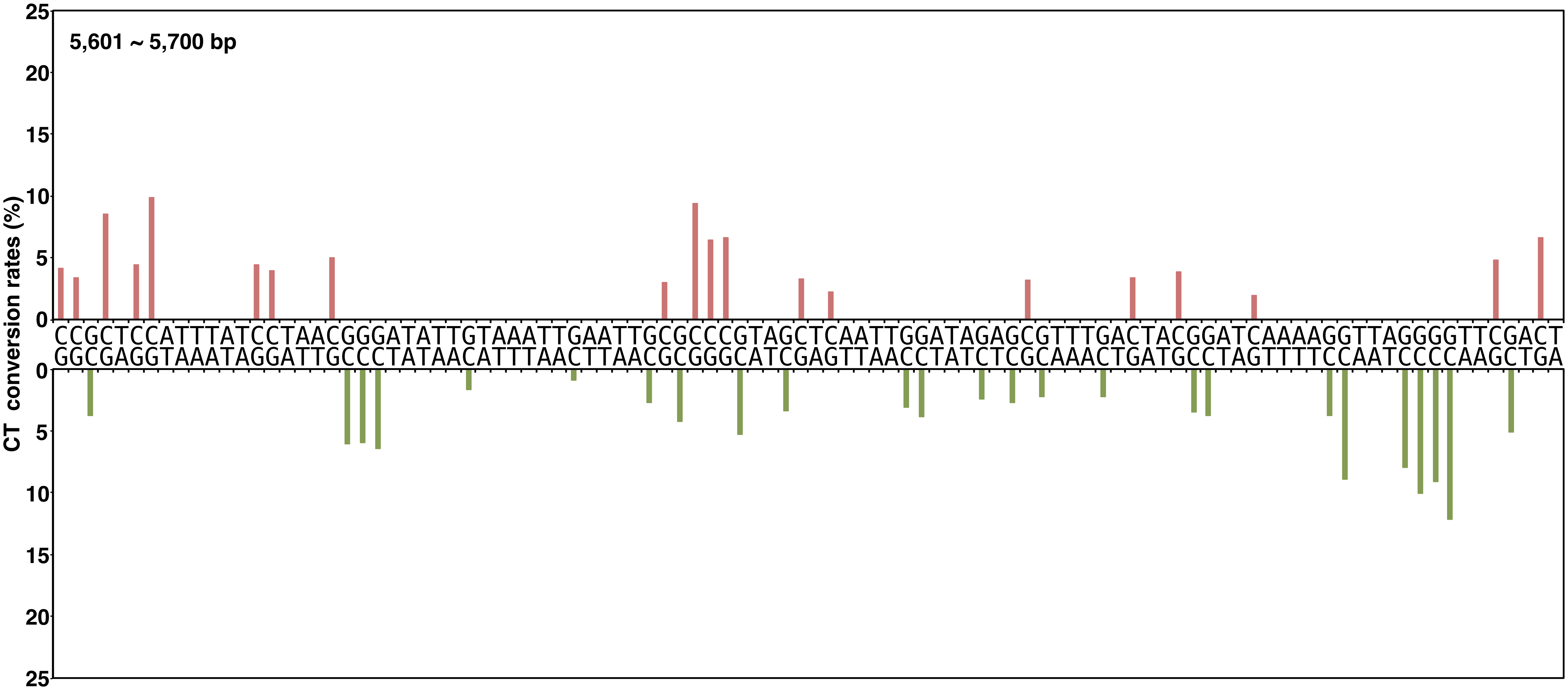


**Yano, Noguchi, and Niki Figure S2, continued**

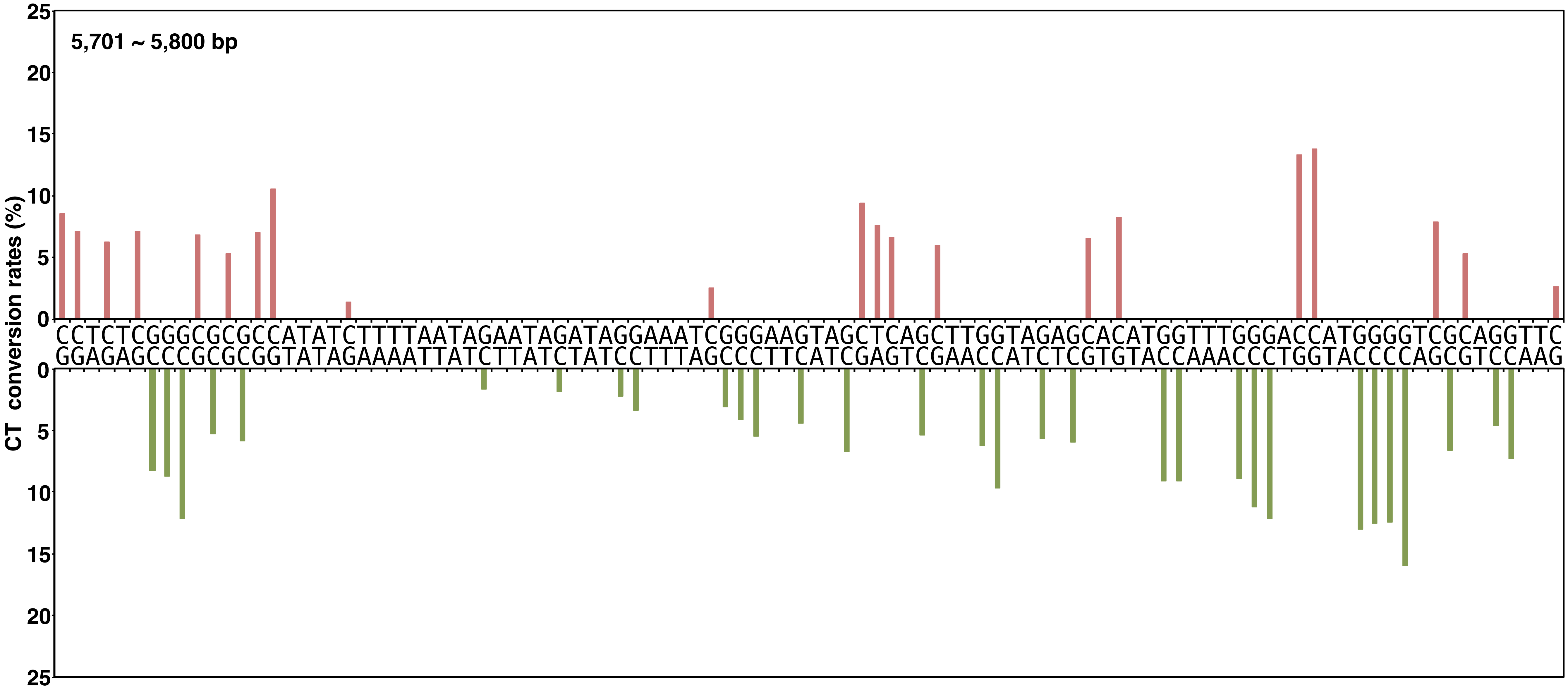


**Yano, Noguchi, and Niki Figure S2, continued**

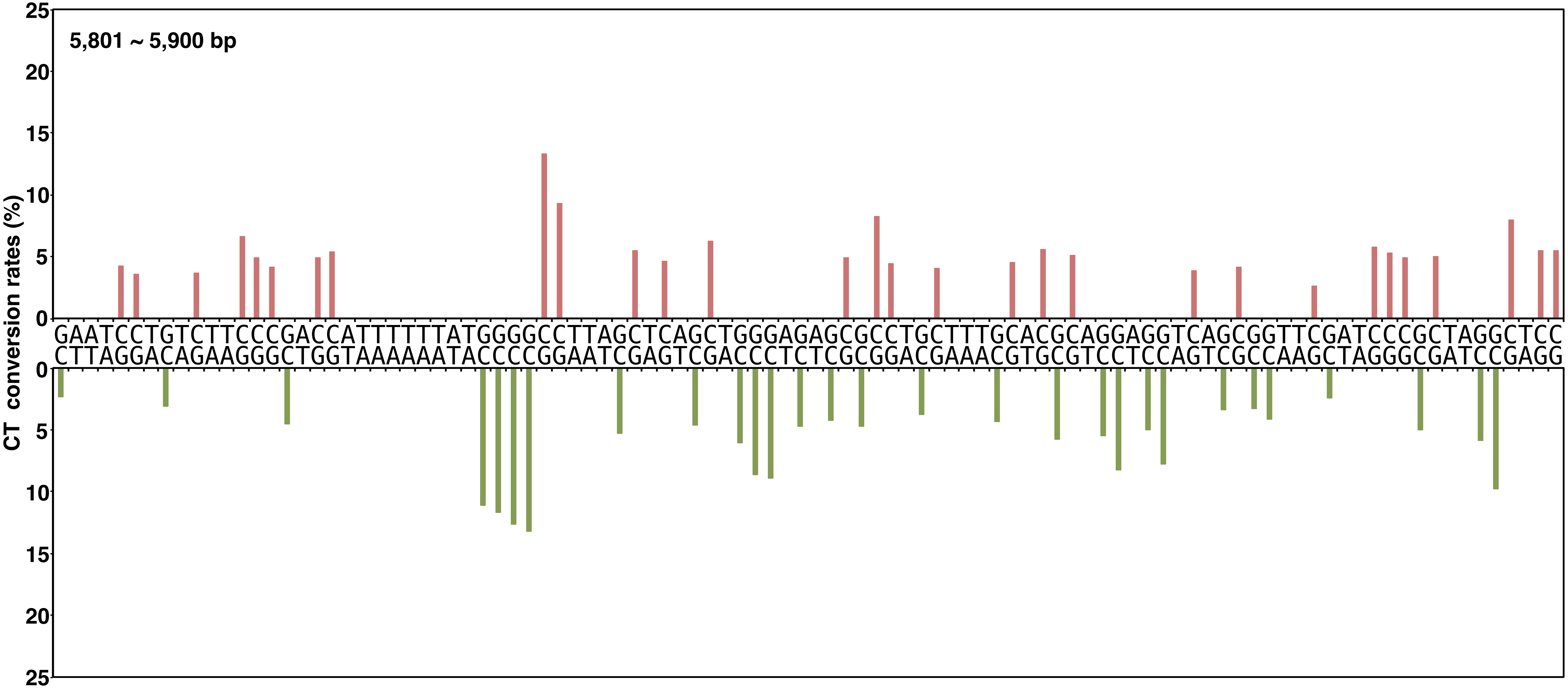


**Yano, Noguchi, and Niki Figure S2, continued**

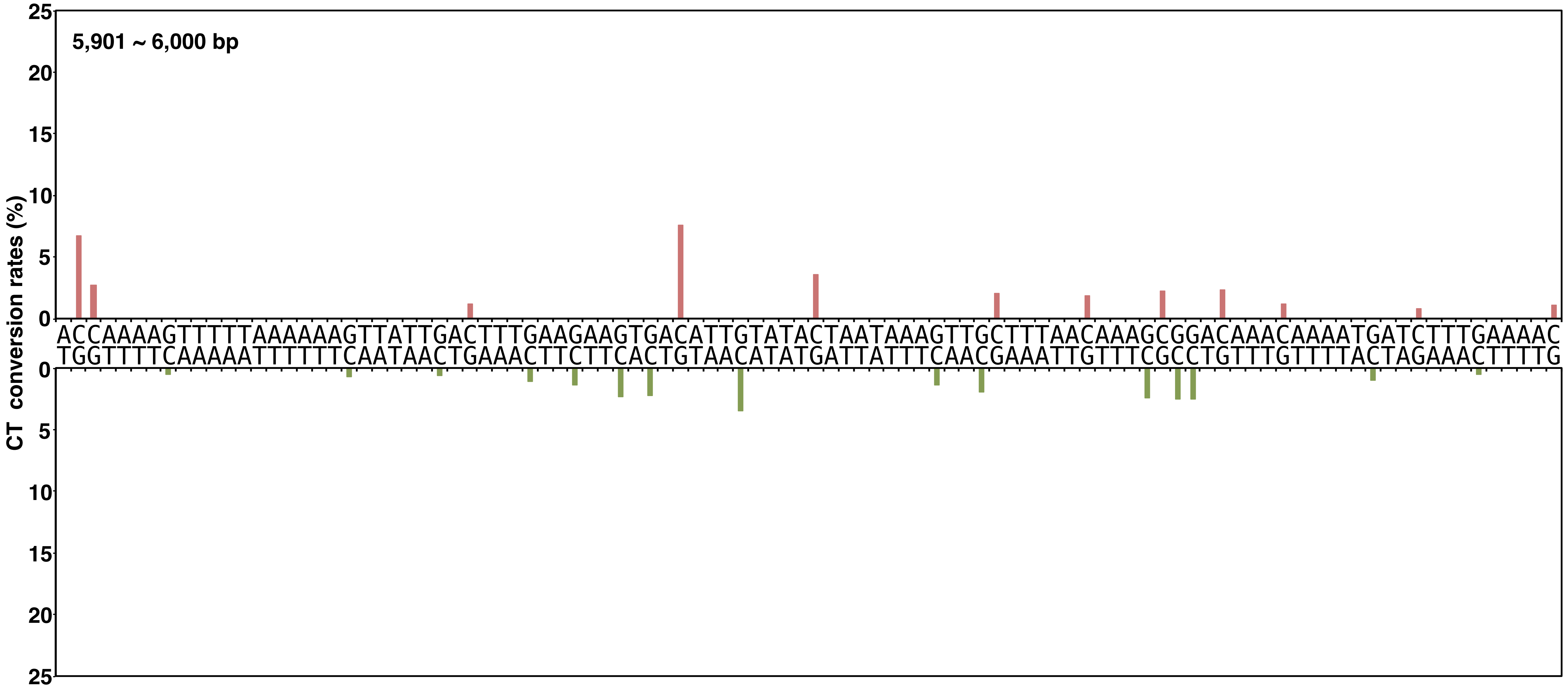


**Yano, Noguchi, and Niki Figure S2, continued**

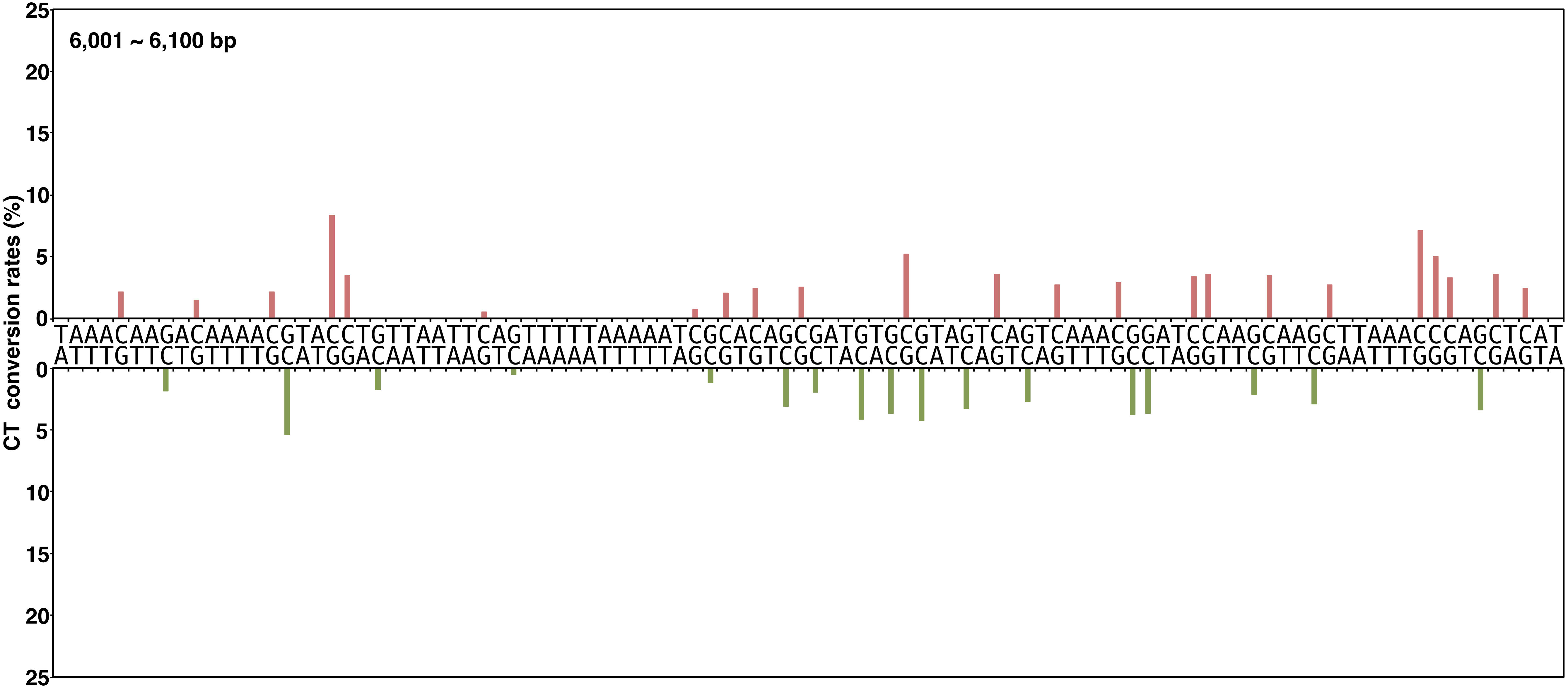


**Yano, Noguchi, and Niki Figure S2, continued**

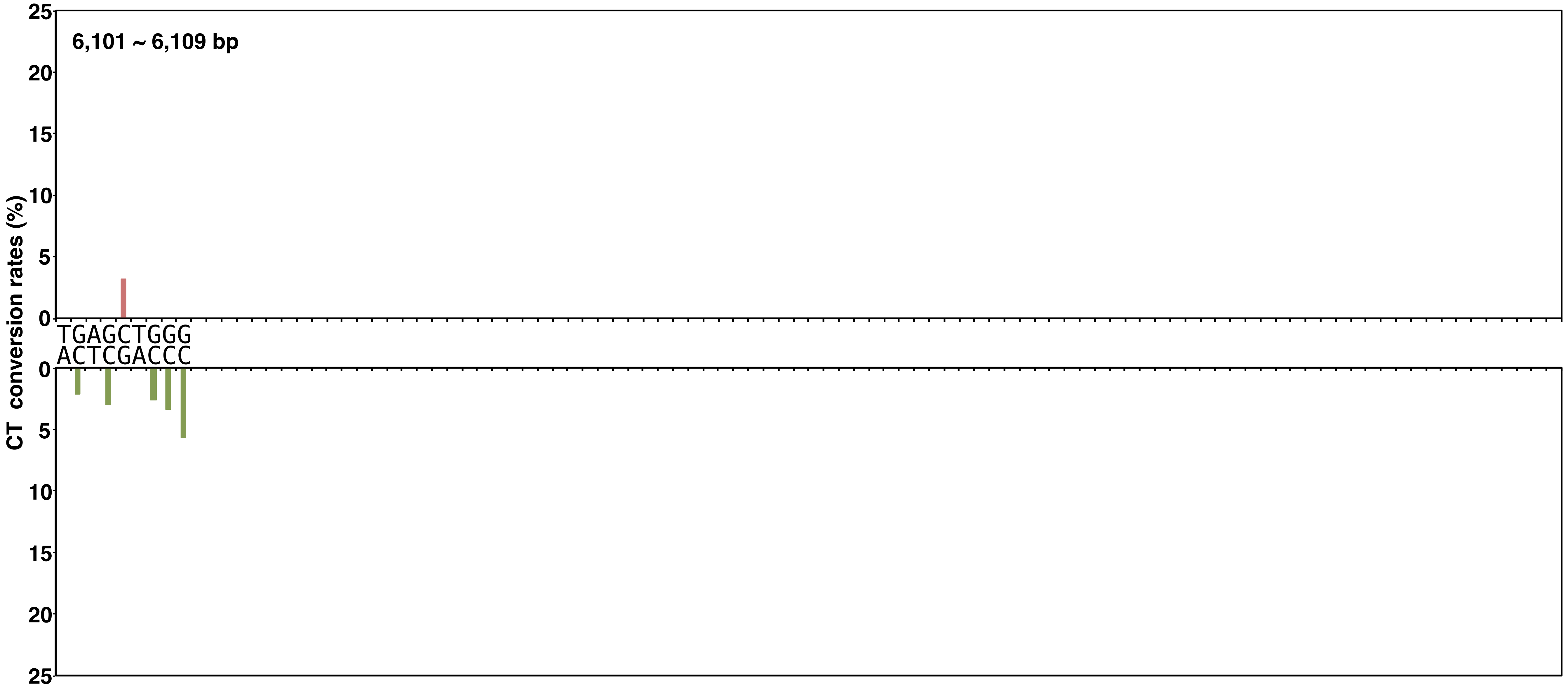


Yano, Noguchi, and Niki Figure S2, continued
