## Supplementary material for "Profiling a single-stranded DNA region within an rDNA segment that is a loading site for bacterial condensin": Yano et al_supplemental files: Yano et al_supplemental files.pdf

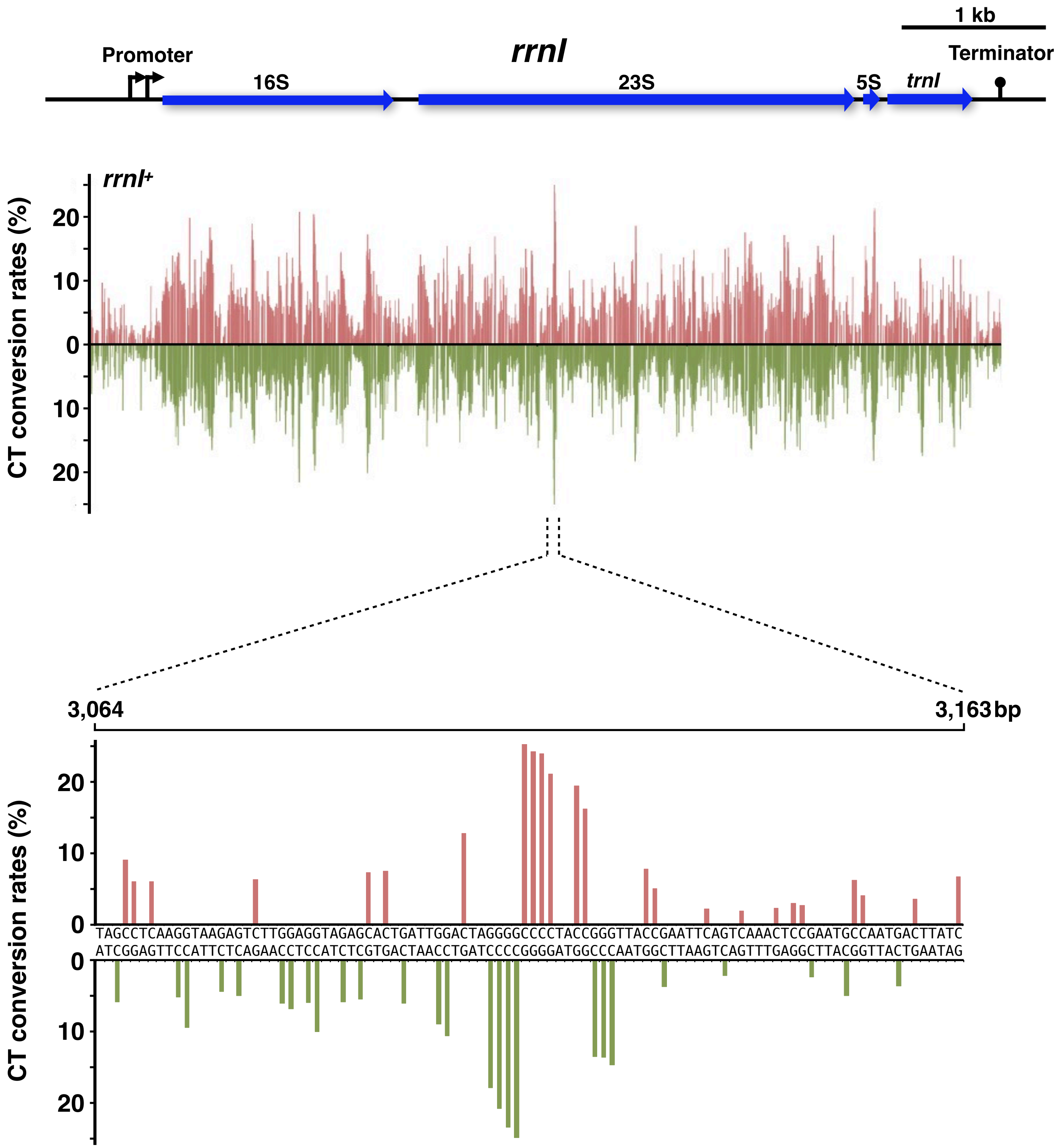


**Yano, Noguchi, and Niki Figure S1**

See the separate PDF file.

Yano, Noguchi, and Niki Figure S2

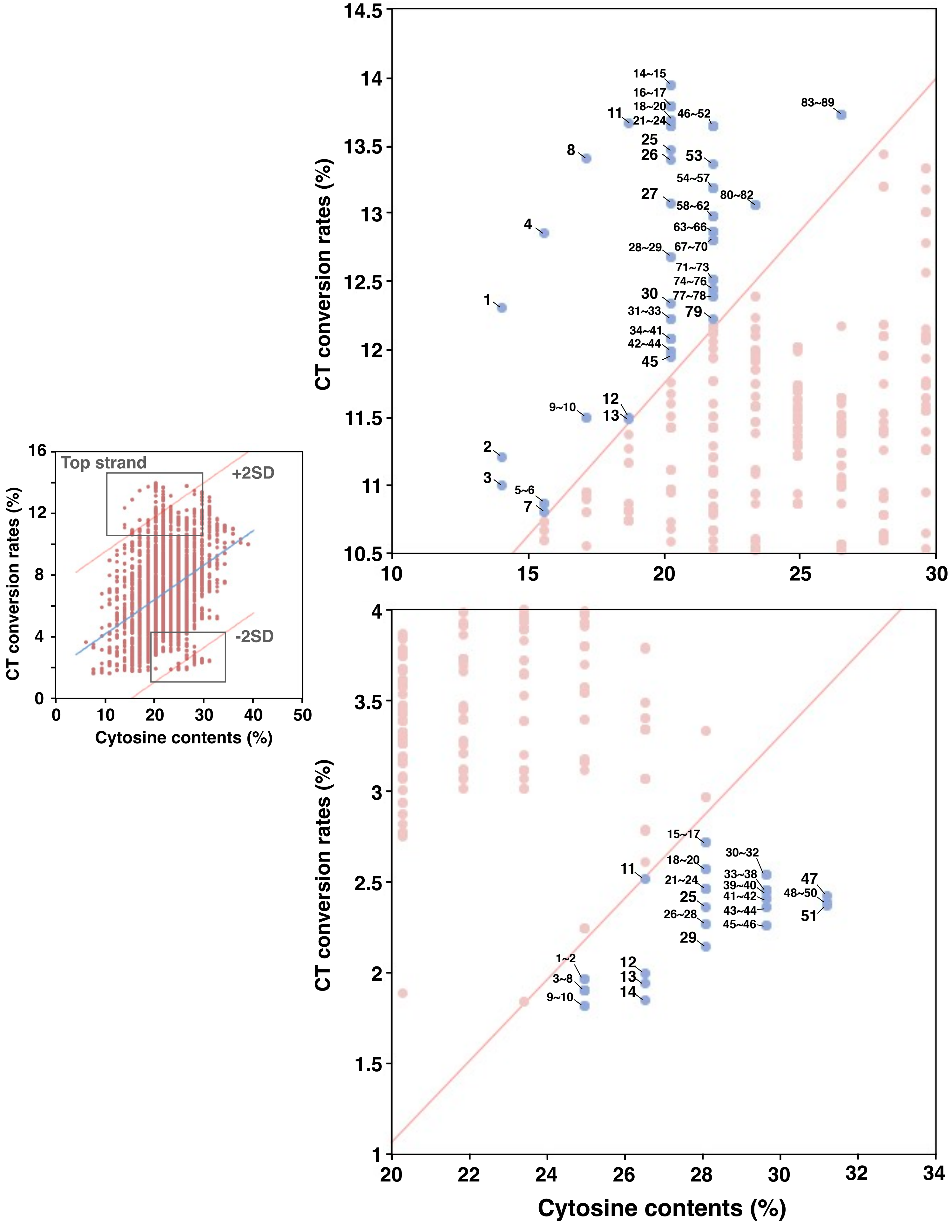


Yano, Noguchi, and Niki Figure S3

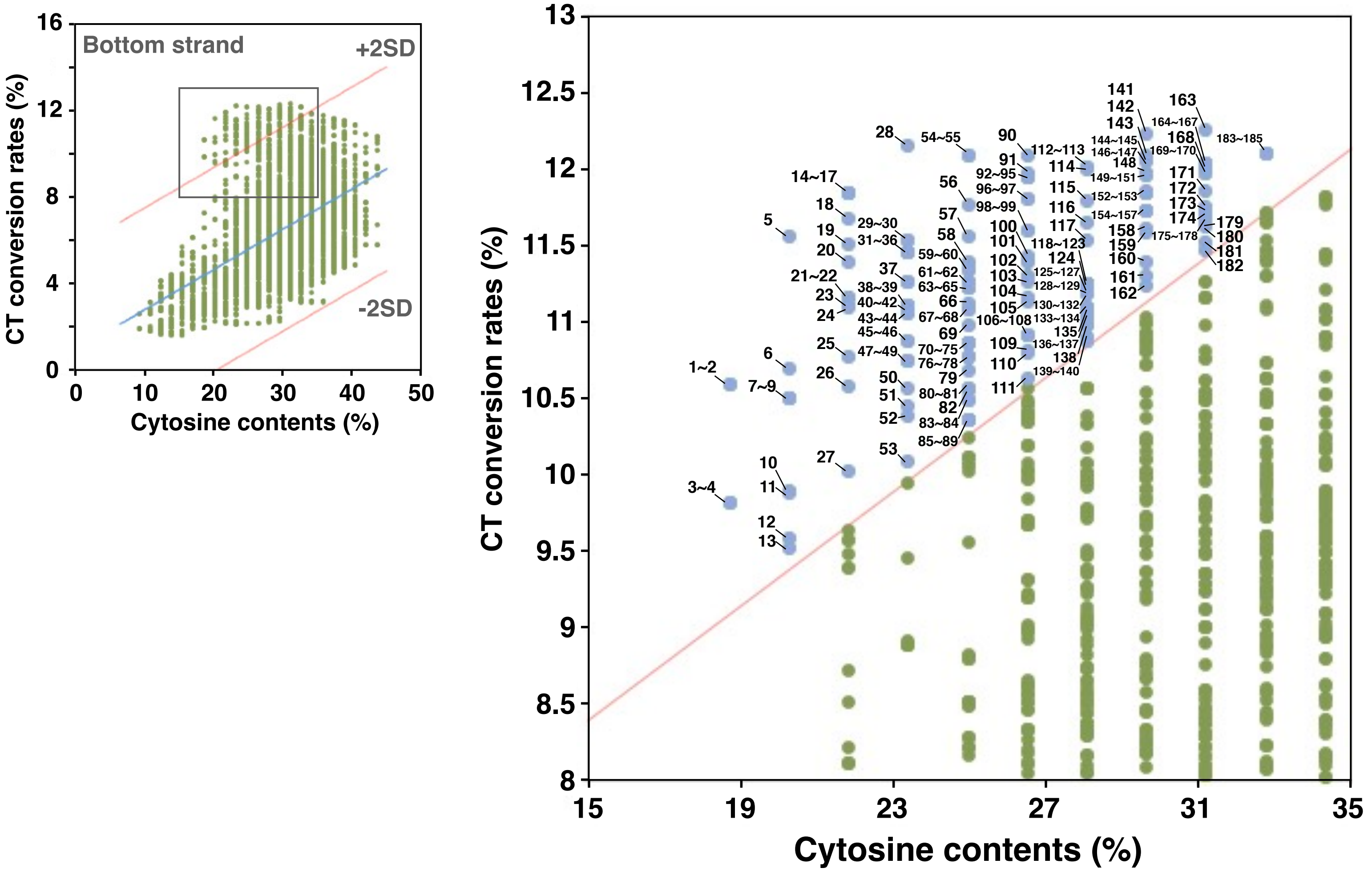


Yano, Noguchi, and Niki Figure S4

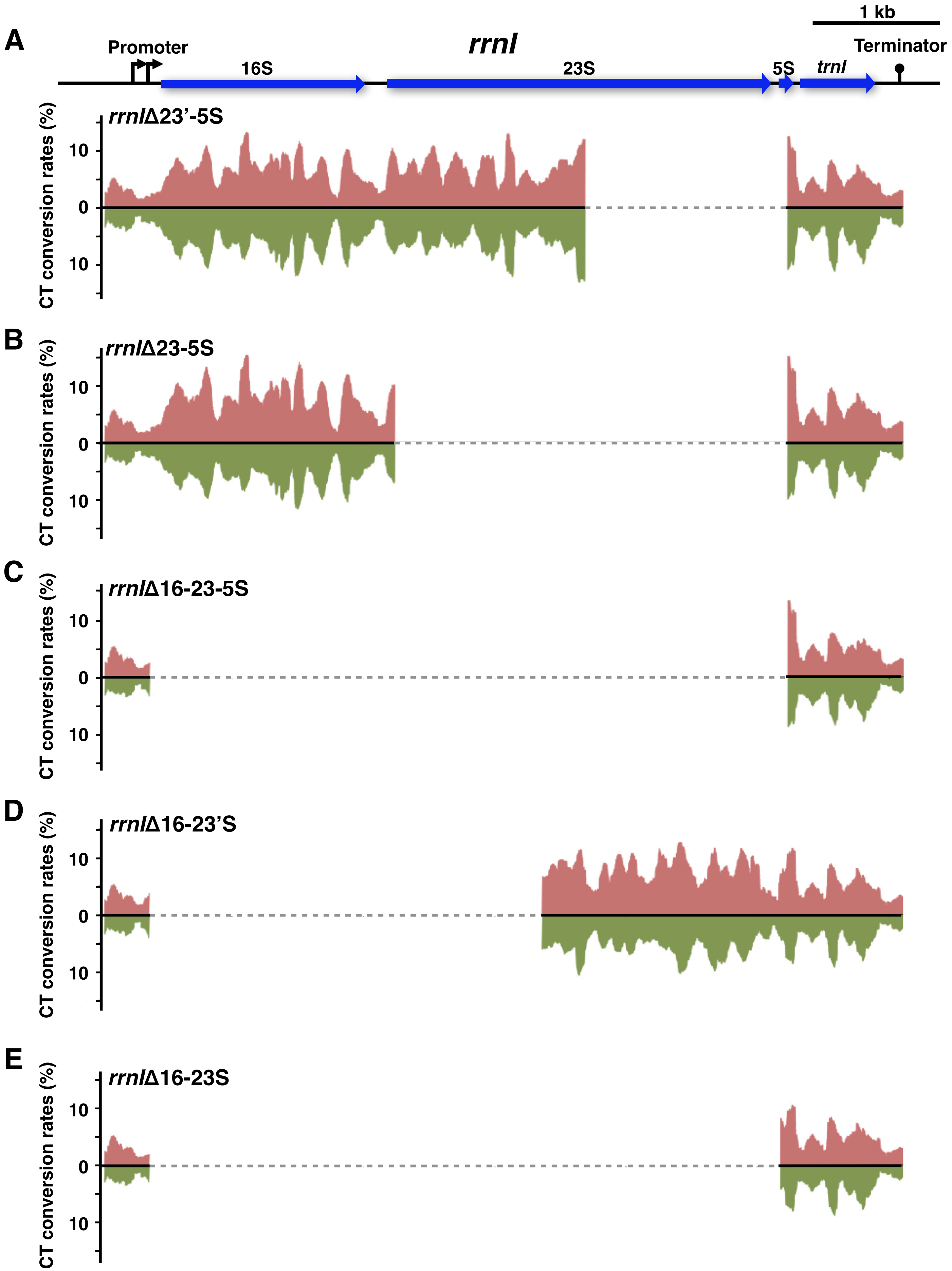


**Yano, Noguchi, and Niki Figure S5**

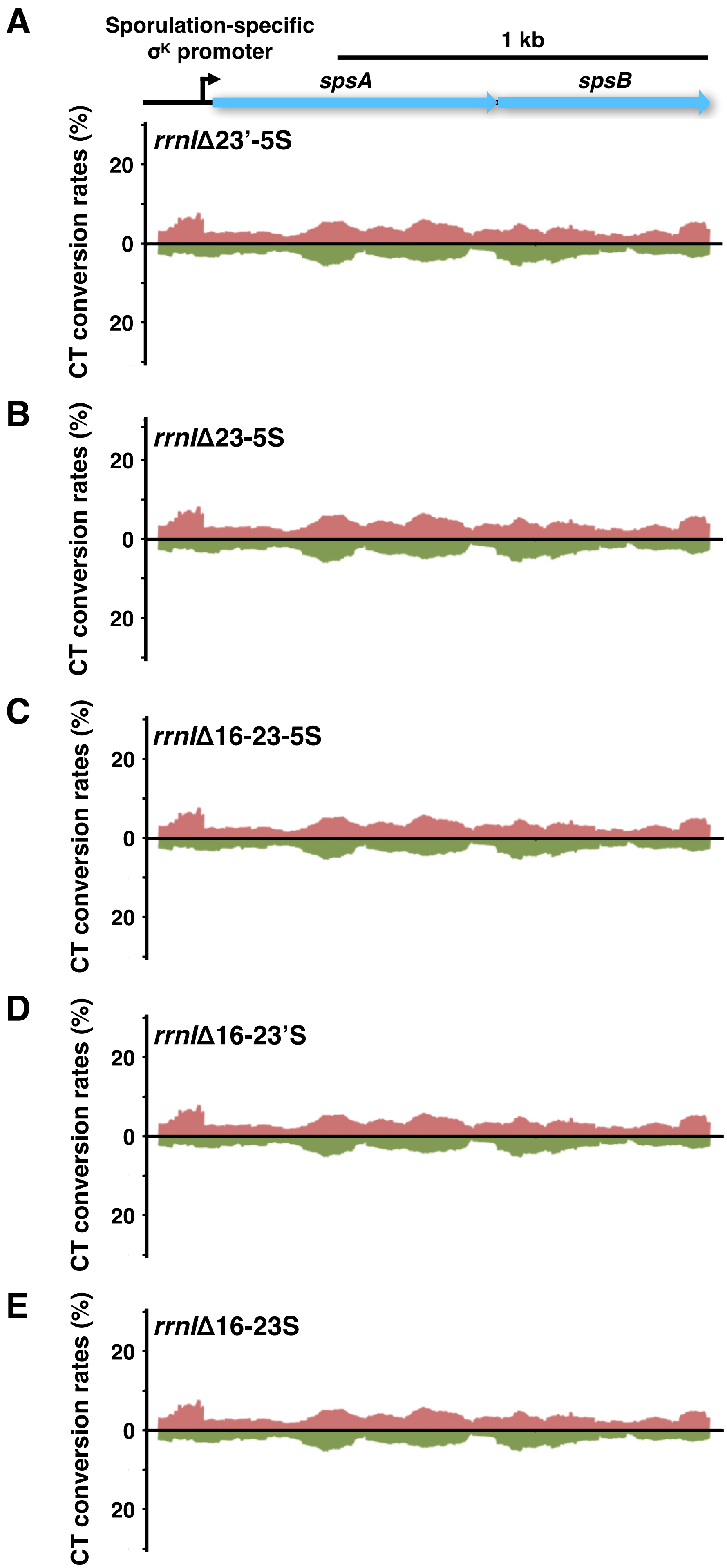


**Yano, Noguchi, and Niki Figure S6**

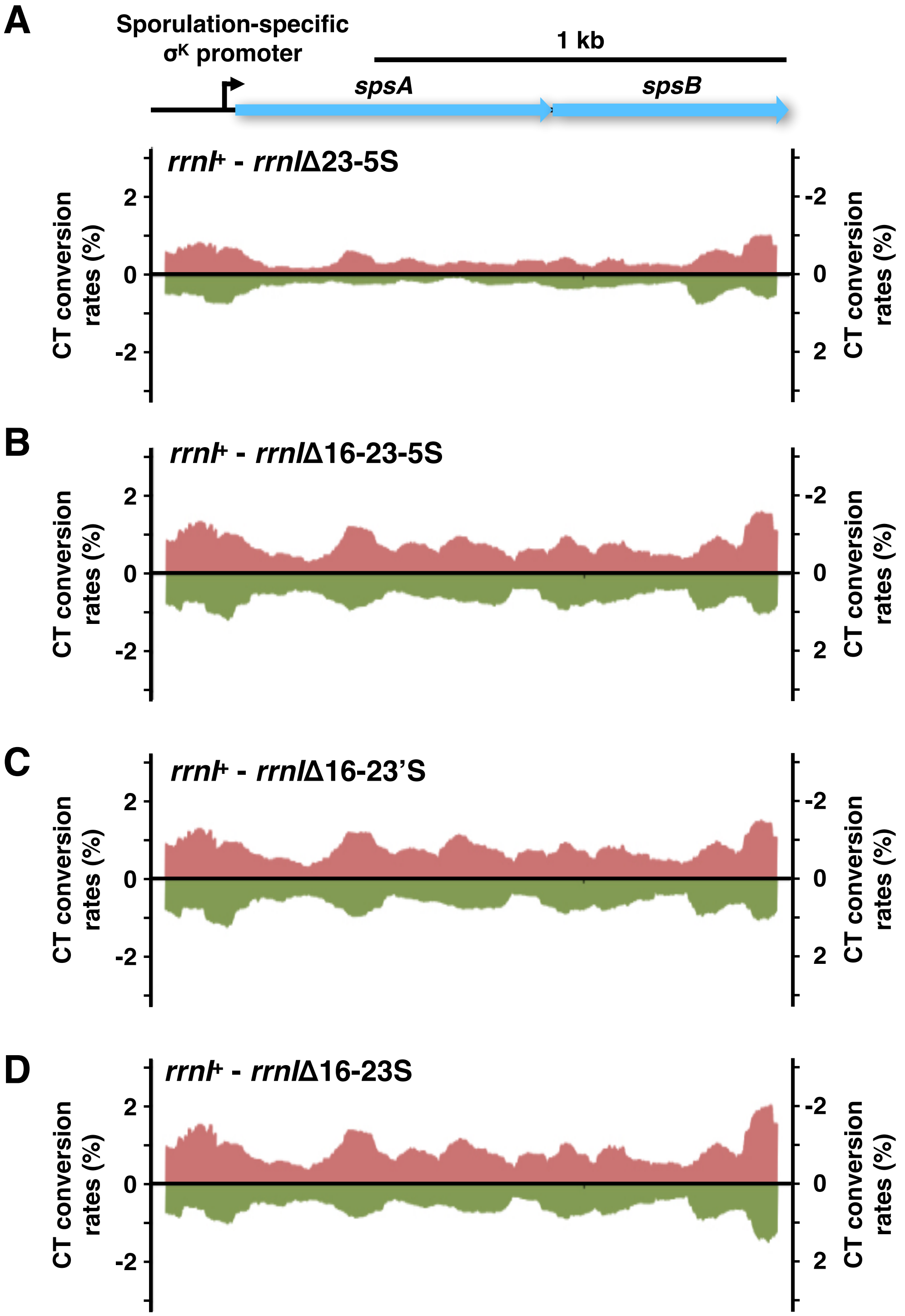


**Yano, Noguchi, and Niki Figure S7**

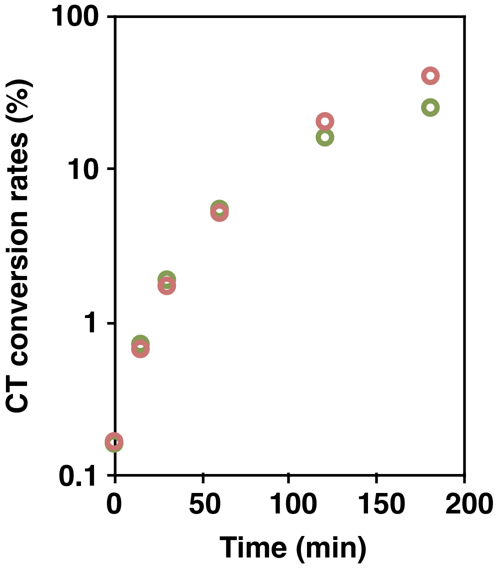


Yano, Noguchi, and Niki Figure S8

A

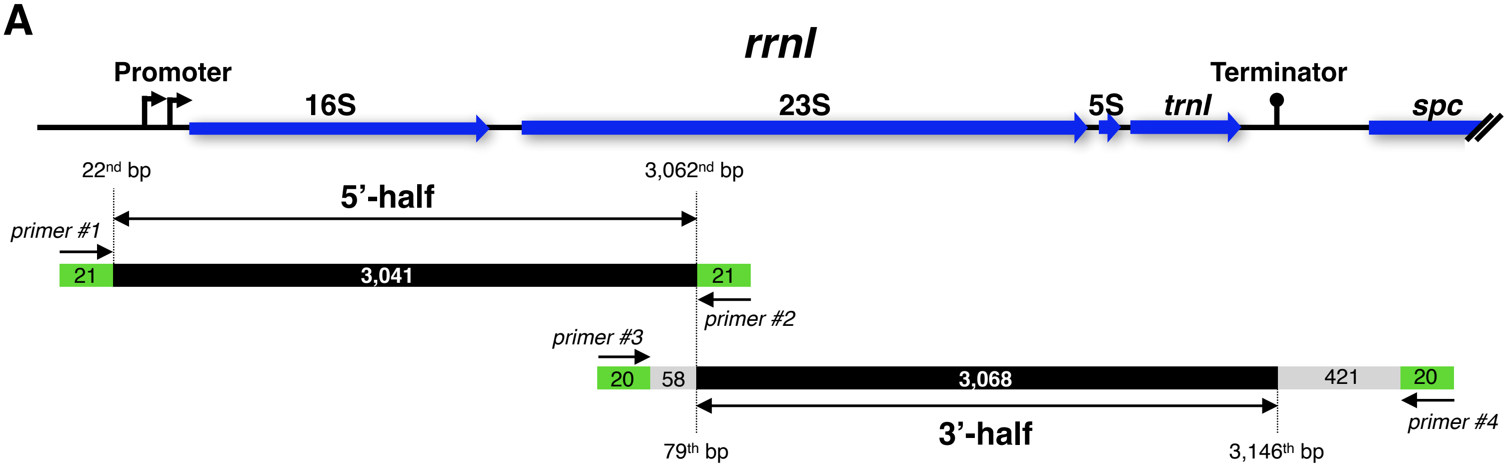


B

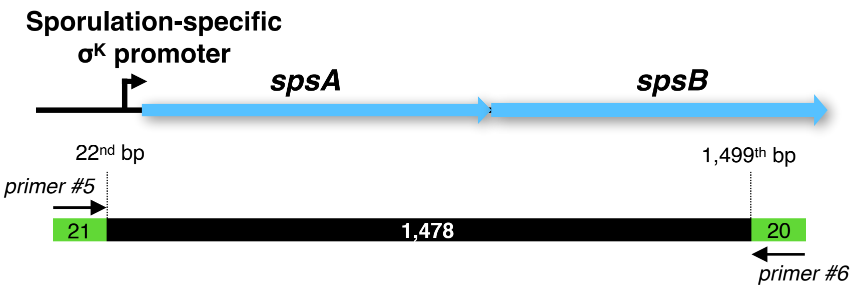


**Yano, Noguchi, and Niki Figure S9**

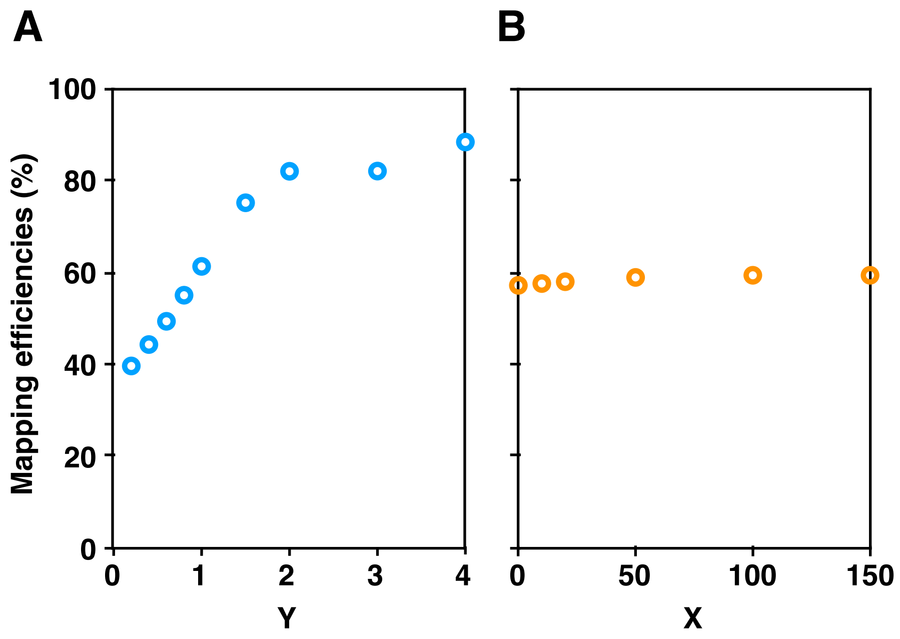


Yano, Noguchi, and Niki Figure S10

**Table S1. Chromosomal region, length, number of cytosines, and CT conversion rates for both top and bottom strands, related to Figures 2 and 7.**

| [a]Chromosomal region | Length (bp) | Number of Cytosines | | CT conversion rates (%) | | | | | | |
|---|---|---|---|---|---|---|---|---|---|---|
| | | | | Both | Top | | | Bottom | | |
| | | Top | Bottom | Mean | Mean | Maximum | Minimum | Mean | Maximum | Minimum |
| **<u>without bisulfite</u>** | | | | | | | | | | |
| 5'-half of *rrnI* (#1 and #2) | 3,041 | 654 | 870 | 0.165 | 0.168 | 0.479 | 0.074 | 0.163 | 1.015 | 0.069 |
| **<u>with bisulfite</u>** | | | | | | | | | | |
| 5'-half of *rrnI* (#1 and #2) | 3,041 | 654 | 870 | 6.89 | 7.14 | 20.7 | 0.838 | 6.70 | 21.5 | 0.808 |
| 3'-half of *rrnI* (#3 and #4) | 3,068 | 686 | 898 | 6.66 | 6.96 | 25.6 | 0.236 | 6.43 | 24.9 | 0.142 |
| full-length of *rrnI* (#1 and #2, #3 and #4) | 6,109 | 1,340 | 1,768 | 6.77 | 7.05 | 25.6 | 0.236 | 6.56 | 24.9 | 0.142 |
| *spsAB* (#5 and #6) | 1,478 | 289 | 348 | 3.90 | 4.20 | 29.3 | 0.514 | 3.65 | 14.5 | 0.448 |

[a] The numbers with # in parentheses indicate the primers which were used for amplification of chromosomal region listed in Table S8.

**Table S2. Nucleotide position, CT conversion rates, and neighboring sequence of the top strand of *rrnl*, related to Figure 2.**

| Nucleotide position | CT conversion rates (%) | Sequence (5' - 3')[a] |
|---|---|---|
| 3113 | 25.56 | GGACTAGGGGCCCCTACCGGG |
| 3114 | 23.93 | GACTAGGGGCCCCTACCGGGT |
| 3115 | 23.66 | ACTAGGGGCCCCTACCGGGTT |
| 5256 | 21.34 | GGGGGTTTCCCCCTGTGAGAG |
| 5257 | 20.96 | GGGGTTTCCCCCTGTGAGAGT |
| 3116 | 20.90 | CTAGGGGCCCCTACCGGGTTA |
| 5255 | 20.83 | CGGGGGTTTCCCCCTGTGAGA |
| 1404 | 20.71 | TTGACGGGGGCCCGCACAAGC |
| 1500 | 20.37 | ATAGGACGTCCCCTTCGGGGG |
| 1501 | 19.88 | TAGGACGTCCCCTTCGGGGGC |
| 669 | 19.75 | ATGGTTCAAACATAAAAGGTG |
| 1499 | 19.29 | GATAGGACGTCCCCTTCGGGG |
| 3119 | 19.18 | GGGGCCCCTACCGGGTTACCG |
| 1086 | 18.92 | TGTGAAAGCCCCCGGCTCAAC |
| 3657 | 18.51 | AATGGGGGGACGCAGGAGGAT |
| 5254 | 18.31 | TCGGGGGTTTCCCCCTGTGAG |
| 805 | 18.26 | TGAGACACGGCCCAGACTCCT |
| 1405 | 18.11 | TGACGGGGGCCCGCACAAGCG |
| 1502 | 17.85 | AGGACGTCCCCTTCGGGGGCA |
| 1087 | 17.76 | GTGAAAGCCCCCGGCTCAACC |
| 5258 | 17.56 | GGGTTTCCCCCTGTGAGAGTA |
| 1505 | 17.53 | ACGTCCCCTTCGGGGGCAGAG |
| 4389 | 17.50 | TGGGATACTACCCTGGCTGTA |
| 1861 | 17.20 | CGTTCCCGGGCCTTGTACACA |
| 4982 | 17.10 | CATGAAGCCCCCCTCAAGATG |
| 4652 | 17.08 | ATAAAAGCTACCCCGGGGATA |
| 4981 | 17.00 | GCATGAAGCCCCCCTCAAGAT |
| 2714 | 16.88 | GTGAAAAGCACCCCGGAAGGG |
| 806 | 16.66 | GAGACACGGCCCAGACTCCTA |
| 4983 | 16.63 | ATGAAGCCCCCCTCAAGATGA |
| 1085 | 16.44 | ATGTGAAAGCCCCCGGCTCAA |
| 1096 | 16.37 | CCCGGCTCAACCGGGGAGGGT |
| 817 | 16.31 | CAGACTCCTACGGGAGGCAGC |
| 4430 | 16.08 | CCTTATCGGGCGGGGAGACAG |
| 4676 | 16.01 | GGCTTATCTCCCCCCAAGAGTC |
| 3120 | 15.96 | GGGCCCCTACCGGGTTACCGA |
| 813 | 15.96 | GGCCCAGACTCCTACGGGAGG |
| 1088 | 15.95 | TGAAAGCCCCCGGCTCAACCG |
| 4419 | 15.71 | TAACCCGCCGCCCTTATCGGG |
| 1511 | 15.64 | CCTTCGGGGGCAGAGTGACAG |
| 1406 | 15.45 | GACGGGGGCCCGCACAAGCGG |
| 4878 | 15.44 | TACGAGAGGACCGGGATGGAC |
| 5194 | 15.40 | AGAGGTCACACCCGTTCCCAT |
| 2394 | 15.37 | CATCTAAGTACCCGGAGGAAG |
| 1084 | 15.31 | GATGTGAAAGCCCCCGGCTCA |
| 2546 | 15.30 | AAAGGCCCGCCATAGGAGGTA |
| 4655 | 15.20 | AAAGCTACCCCGGGGATAACA |
| 4653 | 15.16 | TAAAAGCTACCCCGGGGATAA |
| 4275 | 15.15 | GACGGAAAGACCCCGTGGAGC |
| 4677 | 15.13 | GCTTATCTCCCCCCAAGAGTCC |
| 1097 | 15.12 | CCGGCTCAACCGGGGAGGGTC |
| 4654 | 15.11 | AAAAGCTACCCCGGGGATAAC |

[a] The cytosines whose CT conversion rates were determined are underlined.

**Yano, Noguchi, and Niki Table S2**

**Table S3. Nucleotide position, CT conversion rates, and neighboring sequence of the bottom strand of *rrnI*, related to Figure 2.**

| Nucletide position | CT conversion rates (%) | Sequence (5’ - 3’)[a] |
|---|---|---|
| 3112 | 24.93 | CCGGTAGGGGCCCCTAGTCCA |
| 3111 | 23.42 | CGGTAGGGGCCCCTAGTCCAA |
| 1403 | 21.48 | CTTGTGCGGGCCCCCGTCAAT |
| 3110 | 20.89 | GGTAGGGGCCCCTAGTCCAAT |
| 1860 | 20.01 | GTGTACAAGGCCCGGGAACGT |
| 1508 | 19.64 | TCACTCTGCCCCCGAAGGGGA |
| 1401 | 19.39 | TGTGCGGGCCCCCGTCAATTC |
| 1402 | 19.16 | TTGTGCGGGCCCCCGTCAATT |
| 1859 | 19.14 | TGTACAAGGCCCGGGAACGTA |
| 1510 | 18.87 | TGTCACTCTGCCCCCGAAGGG |
| 1507 | 18.83 | CACTCTGCCCCCGAAGGGGAC |
| 1509 | 18.25 | GTCACTCTGCCCCCGAAGGGG |
| 3654 | 18.20 | CTCCTGCGTCCCCCCATTGCT |
| 3652 | 18.19 | CCTGCGTCCCCCCATTGCTCA |
| 5250 | 18.09 | CAGGGGGAAACCCCCGACTAC |
| 3653 | 17.99 | TCCTGCGTCCCCCCATTGCTC |
| 3109 | 17.91 | GTAGGGGCCCCTAGTCCAATC |
| 3651 | 17.51 | CTGCGTCCCCCCATTGCTCAA |
| 1858 | 17.44 | GTACAAGGCCCGGGAACGTAT |
| 5578 | 17.34 | GAACCCGCGACCCCCACCTTG |
| 3655 | 17.11 | CCTCCTGCGTCCCCCCATTGC |
| 1497 | 17.08 | CCGAAGGGGACGTCCTATCTC |
| 1506 | 16.99 | ACTCTGCCCCCGAAGGGGACG |
| 1865 | 16.88 | GCGGTGTGTACAAGGCCCGGG |
| 5572 | 16.82 | GCGACCCCCACCTTGGCAAGG |
| 5576 | 16.70 | ACCCGCGACCCCCACCTTGGC |
| 5248 | 16.45 | GGGGGAAACCCCCGACTACCA |
| 820 | 16.44 | ACTGCTGCCTCCCGTAGGAGT |
| 4757 | 16.44 | AACAGCCCAACCCTTGGGACC |
| 4433 | 16.40 | ACACTGTCTCCCCGCCCGATA |
| 4657 | 16.39 | CCTGTTATCCCCGGGGTAGCT |
| 4429 | 16.21 | TGTCTCCCCGCCCGATAAGGG |
| 4658 | 16.19 | GCCTGTTATCCCCGGGGTAGC |
| 1400 | 16.17 | GTGCGGGCCCCCGTCAATTCC |
| 819 | 16.10 | CTGCTGCCTCCCGTAGGAGTC |
| 3002 | 16.10 | TTCACCCCTACCCACACCTCA |
| 5790 | 15.97 | GAACCTGCGACCCCATGGTCC |
| 1361 | 15.91 | GACCGTACTCCCCAGGCGGAG |
| 4738 | 15.91 | CCGACTACAGCCCCAGGATGC |
| 5249 | 15.88 | AGGGGGAAACCCCCGACTACC |
| 2258 | 15.88 | CTCCCCGAAGCATATCGGTGT |
| 4434 | 15.69 | GACACTGTCTCCCCGCCCGAT |
| 5575 | 15.69 | CCCGCGACCCCCACCTTGGCA |
| 590 | 15.68 | AGGCAGGTTACCCACGTGTTA |
| 1874 | 15.51 | GTGTGACGGGCGGTGTGTACA |
| 4656 | 15.46 | CTGTTATCCCCGGGGTAGCTT |
| 1100 | 15.41 | AATGACCCTCCCCGGTTGAGC |
| 1695 | 15.35 | ACGTGTGTAGCCCAGGTCATA |
| 3650 | 15.27 | TGCGTCCCCCCATTGCTCAAA |
| 2480 | 15.24 | TGTCCTACAACCCCAAGAGGC |
| 4432 | 15.20 | CACTGTCTCCCCGCCCGATAA |

[a] The cytosines whose CT conversion rates were determined are underlined.

**Yano, Noguchi, and Niki Table S3**

Table S4. Plot number, position, mean CT conversion rates, and mean cytosine contents for windows whose CT conversion rates are greater than the mean + 2SD of the *rrnl* top strand, related to Figure 2.

| Plot No. | Position (bp)[a] | Mean CT conversion rates (%) | Mean cytosine contents (%) |
|---|---|---|---|
| 1 | 1119 | 12.30 | 14.06 |
| 2 | 1120 | 11.19 | 14.06 |
| 3 | 2525 | 10.99 | 14.06 |
| 4 | 1118 | 12.85 | 15.63 |
| 5 | 2523 | 10.86 | 15.63 |
| 6 | 2524 | 10.86 | 15.63 |
| 7 | 2207 | 10.79 | 15.63 |
| 8 | 1117 | 13.40 | 17.19 |
| 9 | 2215 | 11.49 | 17.19 |
| 10 | 2216 | 11.49 | 17.19 |
| 11 | 1116 | 13.65 | 18.75 |
| 12 | 3083 | 11.49 | 18.75 |
| 13 | 4870 | 11.48 | 18.75 |
| 14 | 3100 | 13.93 | 20.31 |
| 15 | 3101 | 13.93 | 20.31 |
| 16 | 1104 | 13.78 | 20.31 |
| 17 | 1105 | 13.78 | 20.31 |
| 18 | 1106 | 13.78 | 20.31 |
| 19 | 1107 | 13.78 | 20.31 |
| 20 | 1108 | 13.78 | 20.31 |
| 21 | 1109 | 13.78 | 20.31 |
| 22 | 1110 | 13.78 | 20.31 |
| 23 | 1111 | 13.78 | 20.31 |
| 24 | 1112 | 13.78 | 20.31 |
| 25 | 1113 | 13.78 | 20.31 |
| 26 | 1114 | 13.78 | 20.31 |
| 27 | 1115 | 13.78 | 20.31 |
| 28 | 1097 | 13.68 | 20.31 |
| 29 | 1098 | 13.68 | 20.31 |
| 30 | 1099 | 13.68 | 20.31 |
| 31 | 3102 | 13.64 | 20.31 |
| 32 | 3103 | 13.64 | 20.31 |
| 33 | 3104 | 13.64 | 20.31 |
| 34 | 3105 | 13.64 | 20.31 |
| 35 | 3087 | 13.46 | 20.31 |
| 36 | 1091 | 13.39 | 20.31 |
| 37 | 1507 | 13.07 | 20.31 |
| 38 | 1516 | 12.67 | 20.31 |
| 39 | 1517 | 12.67 | 20.31 |
| 40 | 583 | 12.33 | 20.31 |
| 41 | 3084 | 12.21 | 20.31 |
| 42 | 3085 | 12.21 | 20.31 |
| 43 | 3086 | 12.21 | 20.31 |
| 44 | 4872 | 12.06 | 20.31 |
| 45 | 4873 | 12.06 | 20.31 |
| 46 | 4874 | 12.06 | 20.31 |
| 47 | 4875 | 12.06 | 20.31 |
| 48 | 4876 | 12.06 | 20.31 |
| 49 | 4877 | 12.06 | 20.31 |
| 50 | 4878 | 12.06 | 20.31 |
| 51 | 4879 | 12.06 | 20.31 |
| 52 | 1528 | 11.97 | 20.31 |
| 53 | 1529 | 11.97 | 20.31 |
| 54 | 1530 | 11.97 | 20.31 |
| 55 | 585 | 11.94 | 20.31 |
| 56 | 3088 | 13.64 | 21.88 |
| 57 | 3089 | 13.64 | 21.88 |
| 58 | 3090 | 13.64 | 21.88 |
| 59 | 3091 | 13.64 | 21.88 |
| 60 | 3092 | 13.64 | 21.88 |
| 61 | 3093 | 13.64 | 21.88 |
| 62 | 3094 | 13.64 | 21.88 |
| 63 | 3099 | 13.35 | 21.88 |
| 64 | 1100 | 13.18 | 21.88 |
| 65 | 1101 | 13.18 | 21.88 |
| 66 | 1102 | 13.18 | 21.88 |
| 67 | 1103 | 13.18 | 21.88 |
| 68 | 1092 | 12.97 | 21.88 |
| 69 | 1093 | 12.97 | 21.88 |
| 70 | 1094 | 12.97 | 21.88 |
| 71 | 1095 | 12.97 | 21.88 |
| 72 | 1096 | 12.97 | 21.88 |
| 73 | 1087 | 12.85 | 21.88 |
| 74 | 1088 | 12.85 | 21.88 |
| 75 | 1089 | 12.85 | 21.88 |
| 76 | 1090 | 12.85 | 21.88 |
| 77 | 3106 | 12.79 | 21.88 |
| 78 | 3107 | 12.79 | 21.88 |
| 79 | 3108 | 12.79 | 21.88 |
| 80 | 3109 | 12.79 | 21.88 |
| 81 | 1504 | 12.51 | 21.88 |
| 82 | 1505 | 12.51 | 21.88 |
| 83 | 1506 | 12.51 | 21.88 |
| 84 | 1508 | 12.43 | 21.88 |
| 85 | 1509 | 12.43 | 21.88 |
| 86 | 1510 | 12.43 | 21.88 |
| 87 | 1511 | 12.38 | 21.88 |
| 88 | 1512 | 12.38 | 21.88 |
| 89 | 1515 | 12.22 | 21.88 |
| 90 | 3096 | 13.06 | 23.44 |
| 91 | 3097 | 13.06 | 23.44 |
| 92 | 3098 | 13.06 | 23.44 |
| 93 | 799 | 13.71 | 26.56 |
| 94 | 800 | 13.71 | 26.56 |
| 95 | 801 | 13.71 | 26.56 |
| 96 | 802 | 13.71 | 26.56 |
| 97 | 803 | 13.71 | 26.56 |
| 98 | 804 | 13.71 | 26.56 |
| 99 | 805 | 13.71 | 26.56 |

[a] Position of 32nd nucleotide in the 64bp window is shown.

**Table S5. Plot number, position, mean CT conversion rates, and mean cytosine contents for windows whose CT conversion rates are smaller than the mean - 2SD of the *rrnI* top strand, related to Figure 2.**

| Plot No. | Position (bp)[a] | Mean CT conversion rates (%) | Mean cytosine contents (%) |
|---|---|---|---|
| 1 | 1801 | 1.96 | 25.00 |
| 2 | 1802 | 1.96 | 25.00 |
| 3 | 1796 | 1.90 | 25.00 |
| 4 | 1797 | 1.90 | 25.00 |
| 5 | 1798 | 1.90 | 25.00 |
| 6 | 1799 | 1.90 | 25.00 |
| 7 | 1789 | 1.90 | 25.00 |
| 8 | 1790 | 1.90 | 25.00 |
| 9 | 1793 | 1.81 | 25.00 |
| 10 | 1794 | 1.81 | 25.00 |
| 11 | 1807 | 2.51 | 26.56 |
| 12 | 1788 | 1.99 | 26.56 |
| 13 | 1800 | 1.93 | 26.56 |
| 14 | 1795 | 1.84 | 26.56 |
| 15 | 1751 | 2.71 | 28.13 |
| 16 | 1752 | 2.71 | 28.13 |
| 17 | 1753 | 2.71 | 28.13 |
| 18 | 1755 | 2.56 | 28.13 |
| 19 | 1756 | 2.56 | 28.13 |
| 20 | 1757 | 2.56 | 28.13 |
| 21 | 1772 | 2.46 | 28.13 |
| 22 | 1773 | 2.46 | 28.13 |
| 23 | 1774 | 2.46 | 28.13 |
| 24 | 1775 | 2.46 | 28.13 |
| 25 | 1782 | 2.35 | 28.13 |
| 26 | 1783 | 2.26 | 28.13 |
| 27 | 1784 | 2.26 | 28.13 |
| 28 | 1787 | 2.14 | 28.13 |
| 29 | 1758 | 2.54 | 29.69 |
| 30 | 1759 | 2.54 | 29.69 |
| 31 | 1760 | 2.54 | 29.69 |
| 32 | 1776 | 2.45 | 29.69 |
| 33 | 1777 | 2.45 | 29.69 |
| 34 | 1778 | 2.45 | 29.69 |
| 35 | 1779 | 2.45 | 29.69 |
| 36 | 1780 | 2.45 | 29.69 |
| 37 | 1781 | 2.45 | 29.69 |
| 38 | 1770 | 2.44 | 29.69 |
| 39 | 1771 | 2.44 | 29.69 |
| 40 | 1761 | 2.40 | 29.69 |
| 41 | 1762 | 2.40 | 29.69 |
| 42 | 1764 | 2.35 | 29.69 |
| 43 | 1765 | 2.35 | 29.69 |
| 44 | 1785 | 2.26 | 29.69 |
| 45 | 1786 | 2.26 | 29.69 |
| 46 | 1769 | 2.41 | 31.25 |
| 47 | 1766 | 2.38 | 31.25 |
| 48 | 1767 | 2.38 | 31.25 |
| 49 | 1768 | 2.38 | 31.25 |
| 50 | 1763 | 2.36 | 31.25 |

[a] **Position of 32nd nucleotide in the 64bp window is shown.**

Table S6. Plot number, position, mean CT conversion rates, and mean cytosine contents for windows whose CT conversion rates are greater than the mean + 2SD of the *rrnI* bottom strand, related to Figure 2.

| Plot No. | Position (bp)[a] | Mean CT conversion rates (%) | Mean cytosine contents (%) | Plot No. | Position (bp)[a] | Mean CT conversion rates (%) | Mean cytosine contents (%) |
|---|---|---|---|---|---|---|---|
| 1 | 4659 | 10.58 | 18.75 | 51 | 1477 | 10.43 | 23.44 |
| 2 | 4660 | 10.58 | 18.75 | 52 | 3141 | 10.37 | 23.44 |
| 3 | 1692 | 9.80 | 18.75 | 53 | 1689 | 10.08 | 23.44 |
| 4 | 1693 | 9.80 | 18.75 | 54 | 1483 | 12.07 | 25.00 |
| 5 | 1878 | 11.54 | 20.31 | 55 | 1484 | 12.07 | 25.00 |
| 6 | 4658 | 10.68 | 20.31 | 56 | 4673 | 11.75 | 25.00 |
| 7 | 4661 | 10.49 | 20.31 | 57 | 1481 | 11.55 | 25.00 |
| 8 | 4662 | 10.49 | 20.31 | 58 | 5586 | 11.37 | 25.00 |
| 9 | 4663 | 10.49 | 20.31 | 59 | 4667 | 11.33 | 25.00 |
| 10 | 1691 | 9.88 | 20.31 | 60 | 4668 | 11.33 | 25.00 |
| 11 | 1698 | 9.87 | 20.31 | 61 | 5584 | 11.25 | 25.00 |
| 12 | 1695 | 9.57 | 20.31 | 62 | 5585 | 11.25 | 25.00 |
| 13 | 1694 | 9.51 | 20.31 | 63 | 3121 | 11.21 | 25.00 |
| 14 | 3131 | 11.83 | 21.88 | 64 | 3122 | 11.21 | 25.00 |
| 15 | 3132 | 11.83 | 21.88 | 65 | 3123 | 11.21 | 25.00 |
| 16 | 3133 | 11.83 | 21.88 | 66 | 5588 | 11.10 | 25.00 |
| 17 | 3134 | 11.83 | 21.88 | 67 | 3124 | 11.07 | 25.00 |
| 18 | 3135 | 11.67 | 21.88 | 68 | 3125 | 11.07 | 25.00 |
| 19 | 3136 | 11.50 | 21.88 | 69 | 1478 | 10.96 | 25.00 |
| 20 | 1875 | 11.38 | 21.88 | 70 | 3139 | 10.84 | 25.00 |
| 21 | 1876 | 11.14 | 21.88 | 71 | 3140 | 10.84 | 25.00 |
| 22 | 1877 | 11.14 | 21.88 | 72 | 1866 | 10.84 | 25.00 |
| 23 | 4665 | 11.09 | 21.88 | 73 | 1867 | 10.84 | 25.00 |
| 24 | 1879 | 11.08 | 21.88 | 74 | 1868 | 10.84 | 25.00 |
| 25 | 4657 | 10.76 | 21.88 | 75 | 1869 | 10.84 | 25.00 |
| 26 | 4664 | 10.57 | 21.88 | 76 | 1861 | 10.76 | 25.00 |
| 27 | 1690 | 10.01 | 21.88 | 77 | 1862 | 10.76 | 25.00 |
| 28 | 1482 | 12.14 | 23.44 | 78 | 1863 | 10.76 | 25.00 |
| 29 | 1479 | 11.52 | 23.44 | 79 | 4653 | 10.66 | 25.00 |
| 30 | 1480 | 11.52 | 23.44 | 80 | 4650 | 10.55 | 25.00 |
| 31 | 3126 | 11.44 | 23.44 | 81 | 4651 | 10.55 | 25.00 |
| 32 | 3127 | 11.44 | 23.44 | 82 | 1686 | 10.53 | 25.00 |
| 33 | 3128 | 11.44 | 23.44 | 83 | 1882 | 10.48 | 25.00 |
| 34 | 3129 | 11.44 | 23.44 | 84 | 1883 | 10.48 | 25.00 |
| 35 | 3130 | 11.44 | 23.44 | 85 | 4644 | 10.34 | 25.00 |
| 36 | 5587 | 11.44 | 23.44 | 86 | 4645 | 10.34 | 25.00 |
| 37 | 4666 | 11.25 | 23.44 | 87 | 4646 | 10.34 | 25.00 |
| 38 | 3137 | 11.24 | 23.44 | 88 | 4647 | 10.34 | 25.00 |
| 39 | 3138 | 11.24 | 23.44 | 89 | 4648 | 10.34 | 25.00 |
| 40 | 1870 | 11.09 | 23.44 | 90 | 4413 | 12.07 | 26.56 |
| 41 | 1871 | 11.09 | 23.44 | 91 | 4414 | 11.96 | 26.56 |
| 42 | 1872 | 11.09 | 23.44 | 92 | 1485 | 11.94 | 26.56 |
| 43 | 1873 | 11.04 | 23.44 | 93 | 1486 | 11.94 | 26.56 |
| 44 | 1874 | 11.04 | 23.44 | 94 | 1487 | 11.94 | 26.56 |
| 45 | 1880 | 10.86 | 23.44 | 95 | 1488 | 11.94 | 26.56 |
| 46 | 1881 | 10.86 | 23.44 | 96 | 3635 | 11.79 | 26.56 |
| 47 | 4654 | 10.73 | 23.44 | 97 | 3636 | 11.79 | 26.56 |
| 48 | 4655 | 10.73 | 23.44 | 98 | 4680 | 11.59 | 26.56 |
| 49 | 4657 | 10.73 | 23.44 | 99 | 4681 | 11.59 | 26.56 |
| 50 | 4649 | 10.55 | 23.44 | 100 | 4674 | 11.42 | 26.56 |

Table S6. Continued.

| Plot No. | Position (bp)[a] | Mean CT conversion rates (%) | Mean cytosine contents (%) |
|---|---|---|---|
| 101 | 4672 | 11.38 | 26.56 |
| 102 | 3631 | 11.28 | 26.56 |
| 103 | 5583 | 11.25 | 26.56 |
| 104 | 3120 | 11.15 | 26.56 |
| 105 | 4669 | 11.11 | 26.56 |
| 106 | 5272 | 10.89 | 26.56 |
| 107 | 5273 | 10.89 | 26.56 |
| 108 | 5274 | 10.89 | 26.56 |
| 109 | 5589 | 10.80 | 26.56 |
| 110 | 3629 | 10.78 | 26.56 |
| 111 | 3628 | 10.62 | 26.56 |
| 112 | 4415 | 12.00 | 28.13 |
| 113 | 4416 | 12.00 | 28.13 |
| 114 | 4412 | 11.99 | 28.13 |
| 115 | 1489 | 11.78 | 28.13 |
| 116 | 3637 | 11.63 | 28.13 |
| 117 | 3634 | 11.52 | 28.13 |
| 118 | 3632 | 11.24 | 28.13 |
| 119 | 4675 | 11.24 | 28.13 |
| 120 | 4676 | 11.24 | 28.13 |
| 121 | 4677 | 11.24 | 28.13 |
| 122 | 4678 | 11.24 | 28.13 |
| 123 | 4679 | 11.24 | 28.13 |
| 124 | 5582 | 11.22 | 28.13 |
| 125 | 4682 | 11.18 | 28.13 |
| 126 | 4683 | 11.18 | 28.13 |
| 127 | 4684 | 11.18 | 28.13 |
| 128 | 3112 | 11.17 | 28.13 |
| 129 | 3113 | 11.17 | 28.13 |
| 130 | 5268 | 11.08 | 28.13 |
| 131 | 5269 | 11.08 | 28.13 |
| 132 | 5270 | 11.08 | 28.13 |
| 133 | 4670 | 11.05 | 28.13 |
| 134 | 4671 | 11.05 | 28.13 |
| 135 | 5581 | 11.01 | 28.13 |
| 136 | 3117 | 10.96 | 28.13 |
| 137 | 5271 | 10.96 | 28.13 |
| 138 | 3630 | 10.89 | 28.13 |
| 139 | 3118 | 10.86 | 28.13 |
| 140 | 3119 | 10.86 | 28.13 |
| 141 | 1494 | 12.21 | 29.69 |
| 142 | 3642 | 12.07 | 29.69 |
| 143 | 3646 | 12.04 | 29.69 |
| 144 | 3643 | 11.96 | 29.69 |
| 145 | 3644 | 11.96 | 29.69 |
| 146 | 814 | 11.95 | 29.69 |
| 147 | 815 | 11.95 | 29.69 |
| 148 | 4409 | 11.84 | 29.69 |
| 149 | 4417 | 11.83 | 29.69 |
| 150 | 1490 | 11.83 | 29.69 |
| 151 | 1491 | 11.83 | 29.69 |
| 152 | 4410 | 11.82 | 29.69 |
| 153 | 4411 | 11.82 | 29.69 |
| 154 | 4404 | 11.72 | 29.69 |
| 155 | 4405 | 11.72 | 29.69 |
| 156 | 4406 | 11.72 | 29.69 |
| 157 | 4407 | 11.72 | 29.69 |
| 158 | 4403 | 11.60 | 29.69 |
| 159 | 3638 | 11.57 | 29.69 |
| 160 | 1380 | 11.38 | 29.69 |
| 161 | 4401 | 11.29 | 29.69 |
| 162 | 3633 | 11.22 | 29.69 |
| 163 | 1499 | 12.24 | 31.25 |
| 164 | 1495 | 12.03 | 31.25 |
| 165 | 1496 | 12.03 | 31.25 |
| 166 | 1497 | 12.03 | 31.25 |
| 167 | 1498 | 12.03 | 31.25 |
| 168 | 1493 | 11.99 | 31.25 |
| 169 | 809 | 11.96 | 31.25 |
| 170 | 810 | 11.96 | 31.25 |
| 171 | 813 | 11.84 | 31.25 |
| 172 | 816 | 11.74 | 31.25 |
| 173 | 4418 | 11.73 | 31.25 |
| 174 | 1492 | 11.70 | 31.25 |
| 175 | 3639 | 11.68 | 31.25 |
| 176 | 3640 | 11.68 | 31.25 |
| 177 | 3641 | 11.68 | 31.25 |
| 178 | 3647 | 11.68 | 31.25 |
| 179 | 4408 | 11.65 | 31.25 |
| 180 | 3645 | 11.61 | 31.25 |
| 181 | 4402 | 11.51 | 31.25 |
| 182 | 4429 | 11.45 | 31.25 |
| 183 | 1500 | 12.09 | 32.81 |
| 184 | 1501 | 12.09 | 32.81 |
| 185 | 1502 | 12.09 | 32.81 |

[a] Position of 32nd nucleotide in the 64bp window is shown.

**Table S7. Mean CT conversion rates on the top and bottom strands at upstream and downstream regions of transcription start site, related to Figures 2 and 7.**

| Chromosomal region | Top strand | | | | | | Bottom strand | | | | | |
|---|---|---|---|---|---|---|---|---|---|---|---|---|
| | Upstream region of transcription start site[a] | | Downstream region of transcription start site[b] | | Whole region[c] | | Upstream region of transcription start site[a] | | Downstream region of transcription start site[b] | | Whole region[c] | |
| | Mean CT conv. rate | Subtracted value[d] | Mean CT conv. rate | Subtracted value[d] | Mean CT conv. rate | Subtracted value[d] | Mean CT conv. rate | Subtracted value[d] | Mean CT conv. rate | Subtracted value[d] | Mean CT conv. rate | Subtracted value[d] |
| ***rrnI*** | | | | | | | | | | | | |
| Wild-type | 3.76 | — | 7.17 | — | 7.05 | — | 3.87 | — | 6.64 | — | 6.56 | — |
| Δpromoter | 3.24 | 0.52 | 5.11 | 2.06 | 5.06 | 1.99 | 3.16 | 0.71 | 5.94 | 0.70 | 5.87 | 0.69 |
| Δ5S | 3.39 | 0.37 | 6.65 | 0.52 | 6.54 | 0.51 | 3.78 | 0.09 | 6.46 | 0.18 | 6.38 | 0.18 |
| Δ23'-5S | 3.42 | 0.34 | 6.20 | 0.97 | 6.07 | 0.98 | 3.44 | 0.43 | 6.36 | 0.28 | 6.25 | 0.31 |
| Δ23-5S | 3.87 | -0.11 | 6.96 | 0.21 | 6.74 | 0.31 | 3.26 | 0.61 | 5.76 | 0.88 | 5.62 | 0.94 |
| Δ16-23-5S | 3.59 | 0.17 | 4.91 | 2.26 | 4.66 | 2.39 | 2.92 | 0.95 | 4.27 | 2.37 | 4.05 | 2.51 |
| Δ16-23'S | 3.52 | 0.24 | 7.05 | 0.12 | 6.80 | 0.25 | 3.08 | 0.79 | 5.41 | 1.23 | 5.28 | 1.28 |
| Δ16-23S | 3.42 | 0.34 | 4.89 | 2.28 | 4.63 | 2.42 | 2.87 | 1.00 | 4.20 | 2.44 | 3.99 | 2.57 |
| Average[e] | 3.49 | 0.27 | 5.97 | 1.20 | 5.79 | 1.26 | 3.22 | 0.65 | 5.49 | 1.15 | 5.35 | 1.21 |
| ***spsAB*** | | | | | | | | | | | | |
| *rrnI*$^+$ | 5.30 | — | 4.07 | — | 4.20 | — | 2.99 | — | 3.70 | — | 3.65 | — |
| *rrnI*Δp | 4.42 | 0.88 | 3.49 | 0.58 | 3.59 | 0.61 | 2.29 | 0.70 | 3.13 | 0.57 | 3.07 | 0.58 |
| *rrnI*Δ5S | 4.52 | 0.78 | 3.55 | 0.52 | 3.65 | 0.55 | 2.40 | 0.59 | 3.30 | 0.40 | 3.23 | 0.42 |
| *rrnI*Δ23'-5S | 4.38 | 0.92 | 3.48 | 0.59 | 3.57 | 0.63 | 2.44 | 0.55 | 3.25 | 0.45 | 3.20 | 0.45 |
| *rrnI*Δ23-5S | 4.64 | 0.66 | 3.67 | 0.40 | 3.77 | 0.43 | 2.53 | 0.46 | 3.44 | 0.26 | 3.37 | 0.28 |
| *rrnI*Δ16-23-5S | 4.28 | 1.02 | 3.32 | 0.75 | 3.42 | 0.78 | 2.26 | 0.73 | 3.05 | 0.65 | 2.99 | 0.66 |
| *rrnI*Δ16-23'S | 4.28 | 1.02 | 3.30 | 0.77 | 3.39 | 0.81 | 2.22 | 0.77 | 3.04 | 0.66 | 2.98 | 0.67 |
| *rrnI*Δ16-23S | 4.16 | 1.14 | 3.20 | 0.87 | 3.30 | 0.90 | 2.33 | 0.66 | 3.06 | 0.64 | 3.00 | 0.65 |
| Average[e] | 4.38 | 0.92 | 3.43 | 0.64 | 3.53 | 0.67 | 2.35 | 0.64 | 3.18 | 0.52 | 3.12 | 0.53 |

[a] Positions of the nucleotide sequence are 1 ~ 261 bp and 1 ~ 129 bp for *rrnI* and *spsAB*, respectively.

[b] Positions of the nucleotide sequence are 262 ~ 6,109 bp and 130 ~ 1,478 bp for *rrnI* and *spsAB*, respectively.

[c] Positions of the nucleotide sequence are 1 ~ 6,109 bp and 1 ~ 1,478 bp for *rrnI* and *spsAB*, respectively.

[d] CT conversion rates of the *rrnI*-deletion mutant were subtracted from that of the wild-type.

[e] Means of the CT conversion rates of the seven *rrnI*-deletion mutants are indicated.

**Table S8. List of primers used in this study, related to STAR Methods.**

| # | Primer name | Sequence (5’ to 3’) | Region |
|---|---|---|---|
| **1** | **rrnI-check-kyF** | GGTCCGGTGTCAGTTGAAAAC | 5’ half of *rrnI* |
| **2** | **seq-rrn-R6** | CCCTAAAGCTATTTCGGAGAG | 5’ half of *rrnI* |
| **3** | **seq-bsrrnO-F6** | AACCCACGCACGTTGAAAAG | 3’ half of *rrnI* |
| **4** | **spc-inR** | CTGTTAATGCGTAAACCACC | 3’ half of *rrnI* |
| **5** | **spsA-upF** | GCTGATGCTTGGATTATGCTG | *spsAB* |
| **6** | **spsB-inR** | GAGGCCATTGATACAGTTGA | *sps*AB |
